# Microstructural and geochemical evidence offers a solution to the cephalopod cameral deposits riddle

**DOI:** 10.1101/2025.02.25.639475

**Authors:** Alexander Pohle, René Hoffmann, Alexander Nützel, Barbara Seuss, Martina Aubrechtová, Björn Kröger, Kevin Stevens, Adrian Immenhauser

## Abstract

Orthoceratoid cephalopods are common in the Palaeozoic rock record but became extinct in the Late Triassic. Many orthoceratoids contain cameral deposits, which are enigmatic calcareous structures within their chambered shell that presumably balanced their straight conchs in a horizontal position. Since the mid-19^th^ century, palaeontologists have attempted to understand the cameral deposit formation process. The various hypotheses include growth from cameral fluids, precipitation by a cameral mantle or even their dismissal as *post-mortem* structures. All of these previous interpretations have in common that they are complicated with contradictory evidence. Here, we present evidence from well-preserved *Trematoceras elegans* specimens from the Late Triassic St. Cassian Formation (Dolomites, northern Italy). We studied the specimens by using optical and electron beam microanalysis techniques and argue that the cameral deposits consist of primary aragonite and calcite fabrics. A fibrous microstructure, which is bilaterally symmetrically arranged with irregularities, is documented. Thin organic sheets originally delimited radial growth sectors. Based on these observations, we propose a new growth model that explicitly involves the cameral sheets. These sheets acted as an extension of the pellicle and held a thin film of supersaturated liquid in the otherwise emptied chambers by the cameral sheets via the capillary effect. Ions were supplied through the siphuncle, such as in living *Nautilus*, and enabled the precipitation of aragonite and calcite fabrics. This model goes beyond previous interpretations, resolves contradictory observations and has functional implications, suggesting that cameral sheets and deposits were an adaptation to increased growth rates.

---

Through geological time, morphological adaptations to a nektic life in the water column are some of the key drivers of cephalopod evolution (e.g., Teichert 1967; Crick 1988; Wade 1988; Kröger *et al*. 2011). This results, among other things, in a conflict between requirements arising from the maintenance of neutral buoyancy while accounting for constraints between manoeuvrability and stability (e.g., Crick 1988; Chamberlain 1993; Peterman *et al*. 2019; Peterman & Ritterbush 2022). Optimisation between these constraints was achieved independently in several lineages of ectocochleate (= externally shelled) cephalopods by evolving a coiled conch (e.g., Flower 1955a; Crick 1988; Chamberlain 1993; Klug & Korn 2004; Naglik *et al*. 2015). Similarly, the internalisation and reduction of the shell in coleoids allowed for greater morphological flexibility in the buoyancy apparatus (e.g., Chamberlain 1993; Kröger *et al*. 2011; Klug *et al*. 2016, 2019). Palaeozoic cephalopods furthermore displayed a variety of relatively short-lived adaptations, such as the periodical truncation of the shell in ascoceratids (Aubrechtová 2019), sphooceratids (Turek & Manda 2012) and potentially in oncoceratids (Stridsberg 1985).

An evolutionary trajectory without modern analogues related to active swimming occurred in orthoceratoid cephalopods (Ordovician to Triassic). Many orthoceratoid taxa contain calcareous structures – termed cameral deposits – within their chambers. These have generally been interpreted as balancing the shell in a horizontal position (e.g., Flower 1955b; Fischer & Teichert 1969; Crick 1988; Chamberlain 1993). This hypothesis has been challenged by more recent 3D hydrostatic models that suggest a vertical *syn vivo* orientation of orthoconic cephalopods, whereby cameral deposits would be insufficient to counterbalance the conch while maintaining neutral buoyancy (Peterman *et al*. 2019). Instead, cameral deposits would merely reduce hydrostatic stability and thus make instantaneous changes in orientation for horizontal movements more energy efficient (Peterman *et al*. 2019). Meanwhile, Mutvei (1956, 2002, 2018) dismissed cameral deposits as part of the living animal entirely and interpreted them as secondary, diagenetic products, potentially involving microbial activity.

Note that a variety of distinct calcareous structures also occurred within the siphuncle of many Palaeozoic cephalopods, called siphuncular deposits (e.g., Flower 1964; Teichert 1964; King & Evans 2019). It is not clear whether all types of siphuncular deposits have a common origin and if they are connected to cameral deposits (King & Evans 2019; Pohle *et al*. 2022a). In this study, we do not investigate siphuncular deposits but we mention them here for completeness, as similar processes might be involved in their formation. The epichoanitic deposits of lituitids are an exception, since their continuity across the siphuncle suggests that they represent a variant of cameral deposits that continued to grow into the siphuncle (e.g., Aubrechtová & Meidla 2019).

Despite repeated attempts over the past 165 years to explain the formation mechanisms of cameral deposits, a universally accepted solution has yet to be presented (e.g., Barrande 1859; Teichert 1933; Flower 1939, 1955b; Mutvei 1956; Fischer & Teichert 1969; Crick 1982; Blind 1991; Kolebaba 2002; Seuss *et al*. 2012a; Mutvei 2018). We make use of state-of-the-art optical and electron beam microscopy techniques to address this long-standing problem and investigate well-preserved specimens of *Trematoceras elegans* (Münster, 1841) from the Carnian (Late Triassic) St. Cassian Formation of northern Italy. We (1) compare our observations with the morphologies to be expected from the previous models of cameral deposit formation and (2) propose a new hypothesis for their growth that is in alignment with physiological and mineralogical constraints.

## Review of previous research

To allow for an in-depth comparison of our observations with previous hypotheses, we provide an overview of the current state of research, since more recent studies did not cover the literature comprehensively (e.g., Seuss *et al*. 2012a; Mutvei 2018). In the earliest descriptions of cameral deposits of orthoceratoid cephalopods, these seem to have been mistakenly interpreted as strongly curved or extremely thickened septa (e.g., Stokes 1840; McCoy 1844), or they were dismissed as *post-mortem* structures shortly thereafter (Woodward 1851). The first to recognise cameral deposits as distinct, organically precipitated parts of the shell was probably Barrande (1859), who regarded them as being formed by the mantle at the same time as the septum. Extensive summaries of the older literature at their time are given by Holm (1885), Teichert (1933), Flower (1955b) and Fischer & Teichert (1969). Historically, cameral deposits have overwhelmingly been studied on longitudinal sections, which show that they are concentrated on the ventral side of the chamber and gradually decrease in thickness and proportional volume towards the aperture (e.g., Flower 1955b, 1964; Teichert 1964; Fischer & Teichert 1969; Blind 1991). Less commonly, these longitudinal sections were supplemented by cross-sections, indicating their morphological complexity and bilaterally symmetric organisation (e.g., Flower 1939; Fischer & Teichert 1969). Rare cases of internal phragmocone moulds exposing the cameral deposits are the only evidence of their three-dimensional structure (Flower 1936; Kolebaba 1975; Evans 1994; Evans *et al*. 2012; Pohle & Klug 2018). These 3D-imprints are more commonly preserved in the Leurocycloceratidae (e.g., Flower 1941; Holland 1964) and Lamellorthoceratidae (e.g., Stanley & Teichert 1976; Bandel & Stanley 1989). Within cephalopods, cameral deposits appear to be restricted to the Orthoceratoidea, implying that these had a single evolutionary origin (King & Evans 2019; Pohle *et al*. 2022a). Their purported presence in other taxa, such as some Tarphyceratida (Ulrich *et al*. 1942), Discosorida (Flower & Teichert 1957) or the Belemnitida (Jeletzky 1966), needs to be reinvestigated.

The distribution of cameral deposits along the phragmocone indicates that they cannot possibly be directly mineralized by the mantle during chamber formation because this would render the animal negatively buoyant for most of its ontogeny. For this reason, Teichert (1933) proposed that tissues within the already-formed chambers were responsible for the formation of cameral deposits. Flower (1939) further developed this hypothesis and established the terms “cameral deposits” and “cameral mantle”. Vascular imprints on the septa of certain orthoceratoids (e.g., Leurocycloceratidae and Lituitidae) were seen as further support for this model (Flower 1941, 1955b, 1964; Holland 1964; Turek & Manda 2012; Aubrechtová & Meidla 2020; Aubrechtová & Korn 2022). However, there is no explanation as to how these living tissues could be supplied with blood through the connecting ring of the siphuncle, which is unperforated (Mutvei 2018). Some authors have therefore suggested that the connecting ring was resorbed during the lifetime of the animal and that the cameral mantle would grow into the chambers from the siphuncle to form cameral deposits later during ontogeny (Kolebaba 1974; Marek 1998; Kolebaba 1999a; Zhuravleva & Doguzhaeva 1999; Kolebaba 2002; Zhuravleva & Doguzhaeva 2004). This hypothesis relies largely on speculation and has therefore received substantial criticism (e.g., Kröger 2008; Turek & Manda 2012; Mutvei 2018). Furthermore, the proposed mechanism cannot explain all cameral deposits, as there are numerous documented examples where they are strongly developed despite the connecting ring being intact. Nevertheless, the epichoanitic deposits of the Lituitida are difficult to explain otherwise, as they represent cameral deposits that apparently continued to grow inside the siphuncle to cover the septal necks (e.g., Sweet 1958; Kröger 2008; Aubrechtová & Meidla 2020).

The full range of problems related to the interpretation of cameral deposit formation become apparent when considering that in one of the most detailed studies on this topic, the two authors came to opposite conclusions (Fischer & Teichert 1969). While Teichert (in Fischer & Teichert 1969) supported the cameral mantle hypothesis, an alternative hypothesis was discussed by Fischer in the same publication, who proposed that the deposits could be precipitated by cameral fluids within the chambers, extending earlier thoughts put forward by Joysey (1961). However, the cameral fluid hypothesis has problems explaining the regular symmetric pattern, for which Fischer acknowledged that there may be different types of cameral deposits, some of which could be formed by cameral tissues (Fischer & Teichert 1969). Nevertheless, other workers have favoured the cameral fluid hypothesis over the cameral mantle hypothesis as a general mechanism (Crick 1982; Crick & Ottensman 1983; Dzik 1984; Bandel & Stanley 1989; Blind 1991; Zhang 1998). Blind (1987, 1988, 1991) reaffirmed the *syn-vivo* deposition of cameral deposits and supported the notion that cameral deposits were precipitated by cameral liquid. However, he also argued that due to the proximity of the cameral deposits to the body chamber, the animal would be too heavy to be buoyant. Thus, in his growth model, the chamber never contained any gas, and the cameral deposits were directly mineralised from the liquid contained in the chambers (Blind 1991).

Scanning electron microscopy (SEM) has been applied repeatedly to cameral deposits, where in well preserved specimens, they have been reported to consist of mainly aragonite, partly diagenetically altered into calcite (Grégoire 1962, 1988; Grégoire & Teichert 1965; Fischer & Teichert 1969; Ristedt 1971; Dauphin 1981; Crick & Ottensman 1983; Bizzarini & Gnoli 1991; Zhuravleva & Doguzhaeva 1999). A few studies suggested that at least some of the calcite is primary (Stehli 1956; Dauphin 1989; Seuss *et al*. 2012a, *b*), while other SEM studies did not specifically address this question (Blind 1987, 1988, 1991; Zhang 1998; Zhuravleva & Doguzhaeva 2004; Doguzhaeva 2019). Using X-ray diffraction with a general area detection diffraction system (XRD-GADDS) in addition to energy dispersive X-ray spectroscopy (EDS), Seuss *et al*. (2012a, b) concluded that the primary calcite was enriched in Mg, i.e., high-Mg calcite, however, without providing quantitative data. The specimen in the latter study displayed evidence of sublethal shell damage, which disturbed the growth of its cameral deposits. Thus, the features seen in this specimen may not be universally transferable. Although Histon (1993) showed the potential of cathodoluminescence (CL) to distinguish between primary and secondary (i.e., inorganic *post-mortem*) deposits and to interpret their diagenesis, it remained the only study applying this technique to cameral deposits to date.

In contrast to microstructural studies, the number of studies applying geochemical proxy data to cameral deposits is limited and has for the most part been restricted to the measurement of elemental concentrations. These studies revealed similar levels of Mg/Ca and Mn/Ca ratios as in the shell wall but also pointed out that the cameral deposits yielded a significantly higher Sr/Ca ratio (Crick & Ottensman 1983; Brand 1987; Dauphin 1989). Likewise, only a single study dealt with the oxygen and carbon isotope values of cameral deposits (Seuss *et al*. 2012b). These authors concluded that the calcitic areas of the cameral deposits were – in part – diagenetically altered, as the isotope geochemical evidence deviated significantly from that of the shell of the same specimen.

Another aspect that merits attention is the fact that, due to the excellent preservation of fossil hard parts, the majority of the previous microanalytical and geochemical studies dealt with cephalopod sample material from the Pennsylvanian Buckhorn Asphalt Lagerstätte. However, several studies have shown that well-preserved cameral deposits are not restricted to this locality (Dauphin 1981, 1989; Bizzarini & Gnoli 1991; Zhang 1998; Zhuravleva & Doguzhaeva 1999, 2004; Doguzhaeva 2019). In the author’s view, it is crucial to expand the taxonomic and temporal scope of studies dealing with cameral deposits of cephalopods. Any model attempting to explain the formation of these structures should be generally applicable to Ordovician to Triassic cephalopods as opposed to one, albeit exceptional, specific Carboniferous Lagerstätte.

## Geological setting

The studied specimens (except one) come from marls of the lower Carnian St. Cassian Formation exposed at the Stuores Wiesen (Prati di Stuores, Stuores Meadows) (e.g., Kittl 1891; Urlichs 1994). It is one of the classical fossil locations of the St. Cassian Formation. The sediments exposed there belong to the *aon* and *aonoides* ammonoid biozones. The St. Cassian Formation consists of predominantly argillaceous sediments deposited in interplatform basins. The surrounding carbonate platforms hosted a rich marine invertebrate biota that thrived in tropical reef and lagoon environments. At the Stuores Wiesen, the basin sediments of the St. Cassian Formation yield autochthonous fossil assemblages of soft bottom dwellers as well as nekton and highly diverse assemblages of fossil material that was transported synsedimentarily from the platform into the basins (Fürsich & Wendt 1977; Hausmann & Nützel 2015; Roden *et al*. 2020). Cephalopods, including the orthoceratoid *Trematoceras*, ammonoids, aulacoceratids and rare nautilids (Leonardi & Polo 1952; Bizzarini & Gnoli 1991), are usually found in basinal deposits lacking transported shallow water benthic fossils or in autochthonous low-diversity assemblages of benthic and nektic fossils. It is not uncommon that autochthonous and allochthonous faunas are mixed.

The preservation of the St. Cassian fossils is variable but can commonly be excellent, including preservation of fine shell ornaments and original shell mineralogy including aragonite preservation. This exceptional preservation is due to the comparably mild degree of diagenetic alteration of the sedimentary rocks. Specifically, the incipient levels of lithification of these deposits are such that fossils can be easily extracted, especially small specimens (liberation Lagerstätte *sensu* Roden *et al*. 2020). Moreover, the argillaceous (low-permeability) nature of these units limited pore water circulation and carbonate-fluid exchange, a feature that contributed to the excellent preservation of fossil remains (Scherer 1977; Roden *et al*. 2020).

One of the studied specimens comes from the locality Staolin, east of Cortina d’Ampezzo (see Zardini 1978). It was found near the Rifugio Mga Lareto. The geological setting and depositional environment are similar to that of Stuores Wiesen but the material is younger (probably *austriacum* ammonoid biozone) and now belongs to the Heiligkreuz Formation (Keim *et al*. 2001). The basinal, marly facies of the Heiligkreuz Formation were commonly also assigned to the St. Cassian Formation (e. g., upper St. Cassian beds).

## Material and methods

### Terminology

The authors wish to point out that the term “deposit” (such as in cameral deposits) is misleading. This is because “deposition” (literally “laying down”) induces a passive process, such as sediment particles leaving their suspension and being gravitationally deposited from a water body onto the lake- or seafloor. As such, the terms “cameral fabrics” or “cameral fillings” would be more appropriate, as these terms are non-interpretative and might include passive, induced or active (biomineralization) secretion of these specific structures. That said, we also realise that the term “cameral deposits”, unsatisfactorily as it might be, is firmly established in the literature and hence, should not be changed. On a similar note, the term “growth” is here used in a descriptive way and does neither imply biogenic or abiogenic formation.

The general morphological terminology of the conch herein follows Teichert (1964) and Pohle *et al*. (2022b). We adopt the terminology of Fischer & Teichert (1969) for cameral deposits, with some modifications (Fig. 1; Table 1). These authors distinguished between planar (plano-) and lateral (latero-) deposits, the former growing parallel to the substrate, while the latter are approximately vertical to the substrate. As the two types transition into each other, lateral deposits effectively represent the rounded edges of the deposits. Crick (1982) argued that essentially all deposits are planar because they grow epitaxially over each other, while nucleating on the shell wall but not on the septa. Continued growth would then result in the rounded edges that cover the septa (Crick 1982). In our opinion, it is still useful to differentiate between the two types of deposits for descriptive purposes. Nevertheless, we replace the term “lateral” with the term “spherulitic”, as it is more intuitive with reference to its shape and also avoids misinterpretations with the orientation of the conch. Deposits may be present anywhere within the chamber, not only laterally.

**Fig. 1.**
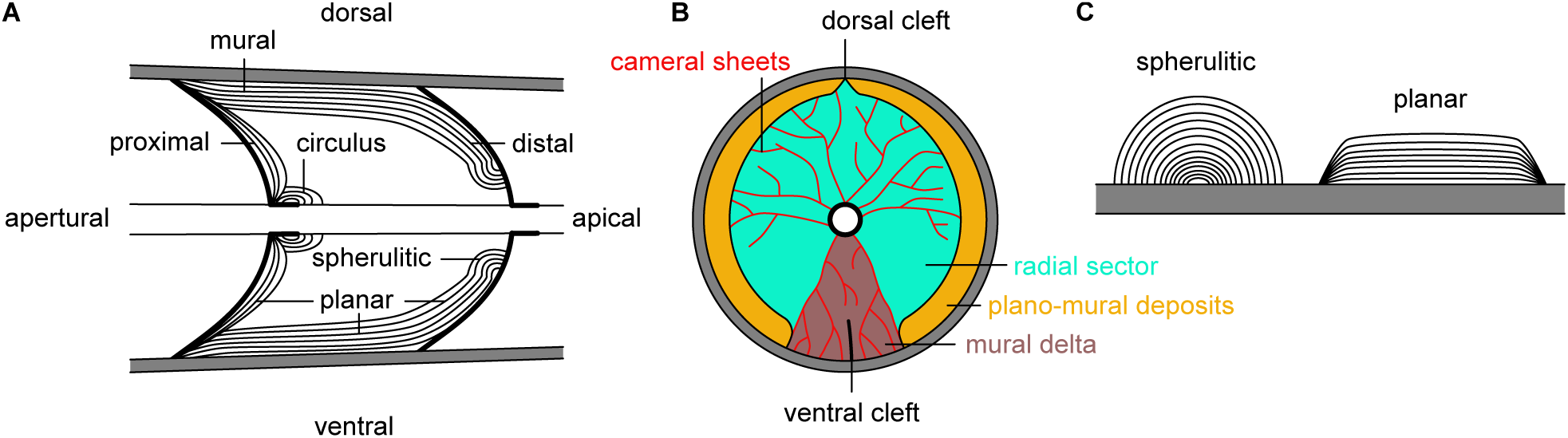
Terminology of cameral deposits in orthoceratoids (see Table 1), based on schematic drawings. A, longitudinal section of a single chamber, showing the proposed terminology. The terms “hyposeptal”, “episeptal” and “lateral” from previous studies are here replaced with “proximal”, “distal” and “spherulitic”. B, cross-section at apical part of a chamber, showing terminology of structurally differentiated areas of cameral deposits. C, schematic illustration of the difference between spherulitic and planar deposits.

**Table 1.** Terminology of cameral deposits and other structures within the chambers of ectocochleate cephalopods.

| Term | Explanation |
| --- | --- |
| Cameral sheets | Thin organic sheets within the chambers of ectocochleate cephalopods. |
| Circulus | Cameral deposits growing on the siphuncle close to the septal necks. |
| Distal | Location reference for cameral deposits. Growth initiation at the end of chamber that is further away (apical, on concave surface of septum) from the body chamber. Corresponds to “episeptal” in previous terminology. |
| Dorsal cleft | Small vertical discontinuity in cameral deposits on dorsal side. |
| Mural | Location reference for cameral deposits. Growth initiation on shell wall. |
| Mural delta | Triangular area at the ventral side of cameral deposits (in cross-section) |
| Pellicle | Thin organic lining on the internal surface of the chambers of phragmocone-bearing cephalopods. |
| Planar | Shape reference for cameral deposits. Grows in parallel, flat layers on top of each other. |
| Proximal | Location reference for cameral deposits. Growth initiation at the end of the chamber that is closer (apertural, on convex surface of septum) to the body chamber. Corresponds to “hyposeptal” in previous terminology. |
| Radial sector | Individual spherulite delimited by cameral sheets. |
| Spherulitic | Shape reference for cameral deposits. Grows from a single point into a spherical shape. Corresponds to “lateral” in Fischer & Teichert’s (1969) terminology. |
| Ventral cleft | Vertical discontinuity in cameral deposits on the ventral side, reportedly reaching up to half of the distance to the siphuncle. |

We furthermore replace the established terms “hyposeptal” and “episeptal” (see Teichert 1964) with the more intuitive terms “proximal” and “distal”, with the soft body (i.e., the body chamber) as reference point. This terminology avoids referring to the septum and defines “proximal” as the part of the chamber that is closer to the body chamber and “distal” as the part of the chamber that is closer to the apex. The traditional terminology implies that the deposits grow on top of (episeptal) or below (hyposeptal) the septum. This orientation corresponds to the orientation on illustrations with the apex downwards that has nothing to do with the *syn-vivo* orientation of the animal, which was either horizontal or apex-up (see Stridsberg 1990; Kröger *et al*. 2011; Pohle & Klug 2018). In addition, previous terminology declares the septum as its reference point, while “cameral deposit” refers to the chamber. Further confusion may arise when considering that the “episeptal” deposits are adapertural to the septum but in the adapical part of the chamber, while it is the other way around for the “hyposeptal” deposit. The use of terms such as “convex” or “concave” is similarly ambiguous because it remains unclear if it refers to the lower or upper growth surface of the cameral deposits. A further positional term is “mural”, which refers to the shell wall (= conotheca), without specifying ventral, dorsal or lateral (Fischer & Teichert 1969).

Following Fischer & Teichert (1969), the above terms can also be combined to refer to the position and shape of the deposits. In accordance with our adjustment in terminology, plano-mural deposits are parallel to the shell wall, and plano-distal and plano-proximal deposits are parallel to the respective septum. Likewise, spherulo-distal and spherulo-proximal deposits are spherulitic on the septum.

Further terms describe particular structural arrangements repeatedly found in cross-sections of cameral deposits. Perhaps the most conspicuous is the “mural delta”, a triangular area on the ventral side of the shell. According to Fischer & Teichert (1969), this area is first devoid of cameral deposits and filled later with spherulitic deposits. In the dorsolateral symmetry plane, there may be discontinuities in the cameral deposits from the shell wall inwards, marking the dorsal and ventral cleft, respectively (Fischer & Teichert 1969). Lastly, the “circulus” denotes cameral deposits growing outwards from the siphuncle (Fischer & Teichert 1969). Circulus, dorsal and ventral cleft play only a minor role in our material but are mentioned here for the sake of completeness.

The taxonomy of Triassic orthoceratoids was recently revised by Pohle & Klug (2024). They argued that given the current state of knowledge, *Trematoceras elegans* Münster, 1841 is the only valid orthoceratoid species in the alpine Triassic. Consequently, we assign all our specimens to this species, especially as our material was collected close to the type locality and horizon of this species.

### Material

We used thin sections of six specimens of *Trematoceras elegans*. The specimens (sections) are housed in the Naturmuseum Bozen under the repository numbers PZO 16535-16540. Except for PZO 16540, which was collected at Staolin near Cortina d’Ampezzo, all come from the Stuores Wiesen (see geological setting). All specimens are phragmocone fragments, and PZO 16539 includes the embryonic shell (Fig. S1). The embryonic shell was sectioned longitudinally. For each of the remaining five specimens, we produced one cross-section and one longitudinal section, respectively. Where multiple thin sections of the same specimens are available, cross- and longitudinal sections are labelled with the suffixes −01 and −02, respectively. We also produced thin sections of three unidentified ammonoid specimens in a similar size range as the *T. elegans* specimens from the Stuores Wiesen. Cross-sections were made of PZO 16557 and PZO 16559, and a longitudinal section of PZO 16558. If cameral deposits were precipitated *post mortem* as suggested by Mutvei (2018), the same structures should be visible in co-occurring ammonoids as well.

### Microscopy

Besides standard transmitted light microscopy (Olympus BX-51) using plane-polarized and cross-polarized light, the thin sections were investigated with electron beam microanalytic techniques. For SEM EBSD (electron backscatter diffraction), surfaces of thin sections were chemo-mechanically etched using colloidal silica (OP-S) for 30 minutes on a vibration polishing machine (manufactures: QATM, model: Saphir Vibro) to reduce surface irregularities and deformed regions on an atomic scale (Massonne & Neuser 2005). Afterwards, thin sections were carbon coated with a 7 nm thick conductive layer. All thin sections were gold coated with a 10 nm conductive layer before SEM-EDS (energy dispersive spectroscopy) and cathodoluminescence (CL) inspection. All thin sections were examined under a high-resolution field emission scanning electron microscope (HR-FESEM) type Zeiss Merlin Gemini 2 using an Everhart-Thornley-Detector (ETD) for secondary electrons imaging at low kV. Elemental concentration and distribution data were collected with 20 kV acceleration voltage and the Oxford X-Max^N^ 150 EDS detector. Crystallographic orientation information was collected with an Oxford Nordlys electron backscatter detector at 20 kV and 70° tilting of the thin section. No filtering or grain segmentation was applied. For mineral identification via Kikuchi bands, we used 10 bands and 80 for Hough space transform. Specimen were further investigated under a cathodoluminescence microscopy type HC8-LM by Lumic equipped with a hot cathode (Neuser *et al*. 1996) and a digital camera system DP73 by Olympus/Evident for recording digital images using the PreCiv software package by Evident. Analytical settings were set to 14 kV acceleration voltage and 0.1 mA probe current. Afterwards, the conductive layer was removed, and thin section surfaces were etched with Di-Na-EDTA (0.27 M) for 5 minutes to improve the contrast between organic-rich and organic-lean regions, dried and again gold coated during inspection under high vacuum in the HR-FESEM.

## Results

### Optical microscopy

An overview of the specimens, the applied microanalytical methods and the visible structures is given in Table 2. Out of the six orthoconic phragmocones (Fig. S1), cameral deposits are well visible in four specimens (PZO 16535, 16538, 16539, 16540; Fig. 2). For details on orthoceratoid specimens lacking cameral deposits, see Appendix S1 and Fig. S2. All three sectioned ammonoids were devoid of cameral deposits (PZO 16557–16559; Fig. S4).

**Fig. 2.**
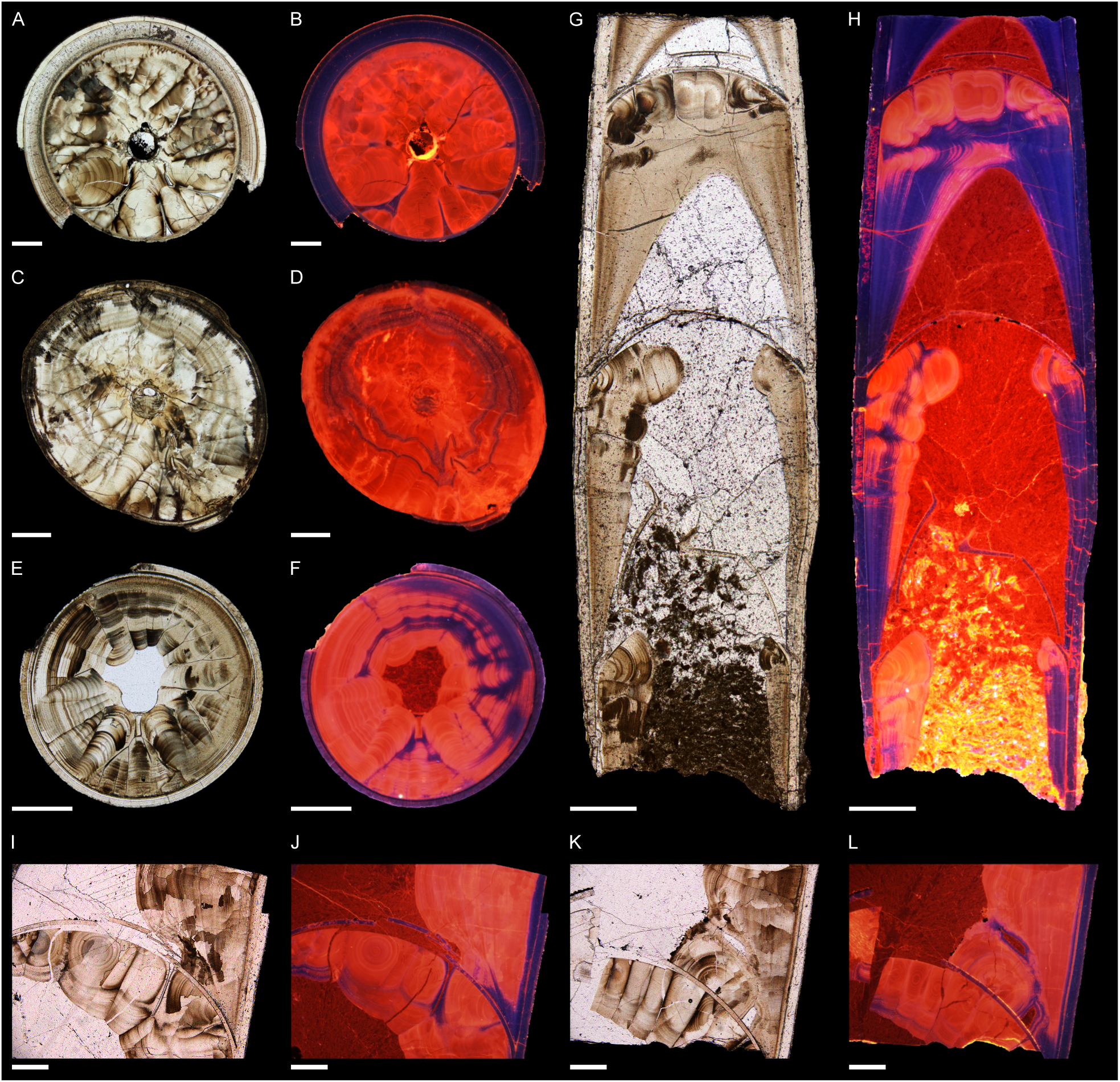
Transmitted light and cathodoluminescence microscopy images of *Trematoceras elegans* from the St. Cassian Formation, Carnian, Late Triassic. All cross-sections are oriented with assumed venter down, and longitudinal sections with apex up. A-B, PZO 16535-01, cross-section (mirrored for consistency). C-D, PZO 16540-01, cross-section. E-F, PZO 16538-01, cross-section. G-H, PZO 16538-01, longitudinal section (mirrored for consistency). I-L, PZO 16535-02, longitudinal section, successive chambers. Brownish areas in TL are regarded as organic rich, while brighter regions are carbonate rich. Bluish regions in CL images are regarded as not diagenetically altered, while reddish-orange to yellowish regions are regarded as diagenetically overprinted or primarily enriched in Mn^2+^ cations. All scale bars represent 0.5 mm.

**Table 2.** Overview of the studied specimens.

| Specimen | Taxon | Section | Size | Locality | Zone | Structures | Methods | Figures |
| --- | --- | --- | --- | --- | --- | --- | --- | --- |
| PZO 16535-01 | *T. elegans* | cross | 5 mm | Stuores | *aon/aonoides* | CS, PMD, SDD, CR | TL, CL, EBSD | 2A-B; 3E; 4E-J; S1C-D; S9; S10 |
| PZO 16535-02 |  | longitudinal | 7.5 mm |  |  | CS, PMD, SDD, MD, SPD | TL, CL, SEM | 2I-L; 8; S1C-D; S7; S8; S17; S18A |
| PZO 16536-01 | *T. elegans* | cross | 7 mm | Stuores | *aon/aonoides* | N/A | TL, CL | S1E, K; S2A, C, E |
| PZO 16536-02 |  | longitudinal | 11 mm |  |  | N/A | TL, CL | S1E, K; S2B, D, F |
| PZO 16537-01 | *T. elegans* (?) | cross | 2 mm | Stuores | *aon/aonoides* | N/A | TL, CL | S1I-J; S3F |
| PZO 16537-02 |  | longitudinal | 5.5 mm |  |  | N/A (?) | TL, CL | S1I-J; S3A-E |
| PZO 16538-01 | *T. elegans* | cross | 2 mm | Stuores | *aon/aonoides* | CS, PMD, SDD, MD, BS, CR | TL, CL, SEM, EDS | 2E-F; 3A-B, D; 6; S1F-G; S11B, D, F; S12; S13; S18B; S21 |
| PZO 16538-02 |  | longitudinal | 8 mm |  |  | CS, PMD, SDD, SN | TL, CL, SEM, EBSD, EDS | 2G-H; 3C; 4A-D; 5A-E; 7; S1F-G; S14; S18C; S19; S22 |
| PZO 16539 | *T. elegans* | longitudinal | 4.5 mm | Stuores | *aon/aonoides* | ES, PMD, SDD, SPD, SD (?), CR, SN | TL, CL | S1H; S5 |
| PZO 16540-01 | *T. elegans* | cross | 3.5 mm | Staolin | *austriacum* | CS, SDD, MD, DC, SID, CR | TL, CL, SEM, EDS | 2C-D; 5F; S1A-B; S6A, C, E; S11A, C, E; S15; S16; S20 |
| PZO 16540-02 |  | longitudinal | 9 mm |  |  | CS, PMD, SDD, SPD, SD, CR, SN | TL, CL | S1A-B; S6B, D, E |
| PZO 16557 | Ammonoida indet. | cross | 2.5 mm | Stuores | *aon/aonoides* | N/A | TL, CL | S4A, C, E |
| PZO 16558 | Ammonoida indet. | longitudinal | 7 mm | Stuores | *aon/aonoides* | N/A | TL, CL | S4B, D, E |
| PZO 16559 | Ammonoida indet. | cross | 7 mm | Stuores | *aon/aonoides* | N/A | TL, CL | - |

Size of cross-sections corresponds to the maximum diameter of the conch, and size of longitudinal sections corresponds to the length of the specimen, except for the ammonoid section, where it represents the diameter. Abbreviations of structures: CS = cameral sheets; PMD = plano-mural deposits; SDD = spherulo-distal deposits; SPD = spherulo-proximal deposits; MD = mural delta; DC = dorsal crest; BS = biconcave structure, SID = siphuncular deposits; CR = connecting ring; SN = septal necks; ES = embryonic shell; N/A = chambers devoid of cameral deposits. For abbreviations of methods see text.

In cross-sections, the deposits are organised in radially arranged, sharply delimited spherulitic sectors (Fig. 2A-F). As the position of the siphuncle is situated slightly off-centre, the bilateral symmetry plane can be identified by comparing the centre of the siphuncle to the centre of the conch in undeformed specimens (Fig. 3A). The thereby inferred symmetry of the conch aligns with the bilaterally symmetric pattern of the deposits (Fig. 3B, D, E). A triangular patch on the ventral side corresponds to the mural delta (Fig. 3D, E, S11B, D, E). Furthermore, equal-sized sectors are present on the left and right sides, decreasing in size towards the dorsum (Fig. 2A-F, 3D, E). Except for where the mural delta meets the shell wall, plano-mural deposits cover the entire inner surface of the shell wall and at least the marginal parts of the septum (Fig. 3D, E). These patterns are easily recognised in the cross-sections PZO 16535-01 and PZO 16538-01, the former representing a slightly more apical section that passes through the septum (Fig. 2A, B, E, F, 3D, E). The crosssection PZO 16540-01 is slightly crushed, and some of the structures are obscured, but the larger-sized spherulitic sectors (mural delta) on one side and plano-mural deposits on the other side are indicative of the specimen’s orientation (Fig. 2C, D). In contrast to the other specimens, the plano-mural deposits in PZO 16540-01 converge dorsally, which may indicate a dorsal cleft (Fig. 2C, D, S11A, C, E). Furthermore, the specimen is unique in that the deposits appear to grow inside the siphuncle, with the connecting ring apparently presenting no obstacle (Fig. S6). The symmetry of the cross-section can also be recognised in the plano-mural deposits, where the layers are thicker on one side and gradually thin towards the opposite side of the shell (Fig. 3E).

**Fig. 3.**
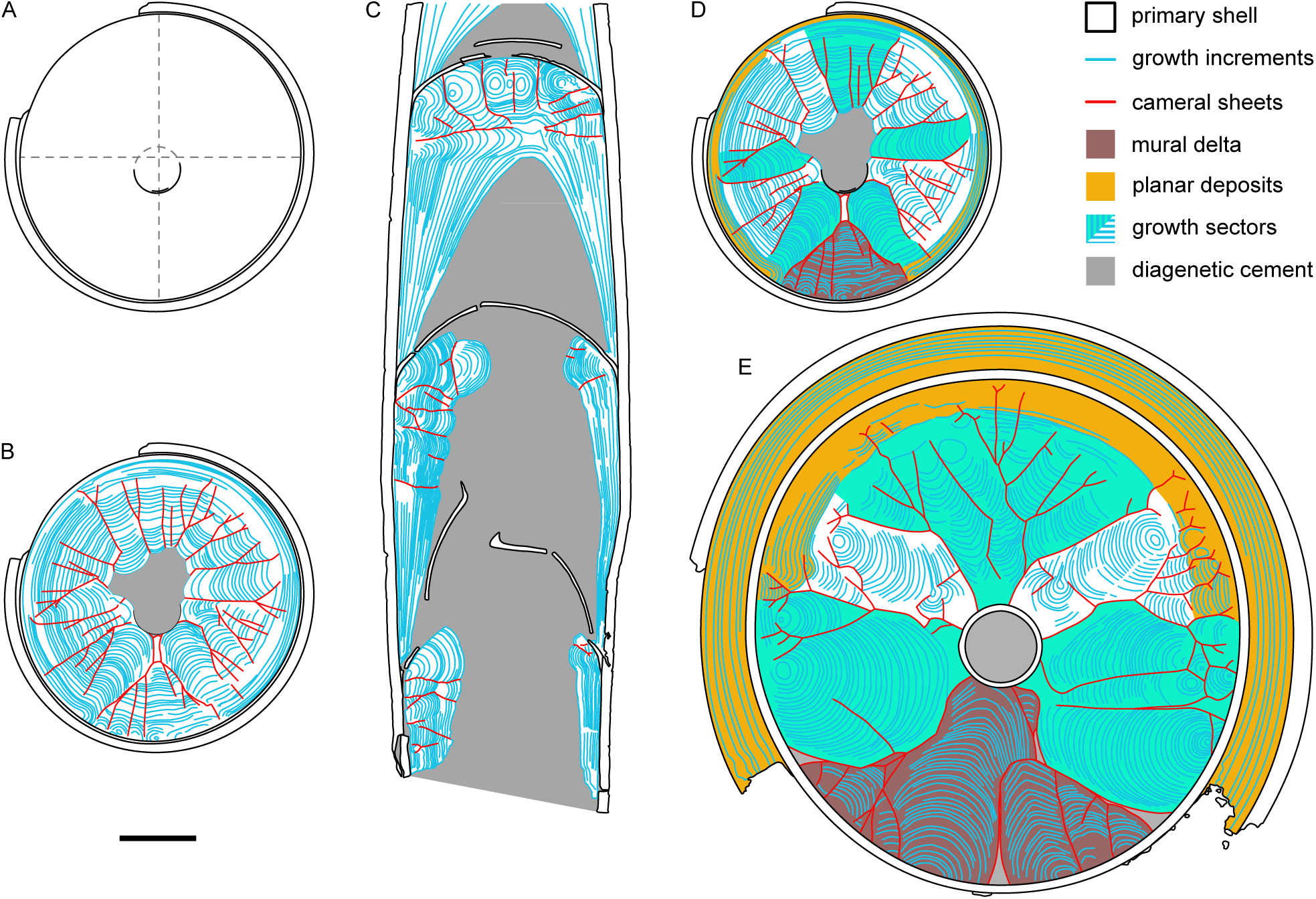
Interpretative drawings of structural organisation of cameral deposits and cameral sheets in *Trematoceras elegans*. A, PZO 16538-01, cross-section (see Fig. 2E-F). Primary shell and siphuncle only, with cross hair added to show offset of siphuncle from the centre, allowing the establishment of the dorsoventral symmetry plane. B, same specimen, cross-section, with cameral deposits and sheets added. C, same specimen, PZO16538-02, longitudinal section (see Fig. 2G-H; mirrored for consistency). D, same specimen, PZO 16538-01, cross-section with added colours to show differentiation of cameral deposits. E, PZO 16535, cross-section (see Fig. 2A-B; mirrored for consistency) with colours as comparison. Scale bar represents 0.5 mm.

As the cross-sections were made slightly adaperturally of the septum, they expose distal and mural deposits. Longitudinal sections show that these are the prevailing types of cameral deposits in *T. elegans* (Fig. 2G-L). Plano-mural deposits transition into spherulo-distal deposits at the septum and consist of progressively more growth increments in apical chambers (Fig. 2G, H, 3C). PZO 16535-02 possesses spherulo-proximal deposits on one side (ventral?) of a single chamber (Fig. 2K-L, S8B, D, F, 17). Further proximal deposits are present in the longitudinal sections PZO 16540-02 (Fig. S6B, D, F) and in the sectioned embryonic shell PZO 16539-02 (Fig. S5), although both specimens show evidence for diagenetic alteration in the orange luminescence of the shell wall. The growth lamellae of these deposits are also not well visible.

The septa and the outer shell of the specimens display a bluish cathodoluminescence colour (e.g., Fig. 2B, D, F, H, J, L). Within the cameral deposits, both blue and orange luminescence colours are found (Fig. 2). Growth increments are characterised by a range of orange or blue luminescence colours. The plano-mural deposits exhibit blue luminescence colour in most cases (Fig. S10A-D). The distal and proximal spherulitic deposits commonly display an orange luminescence (Fig. 2, S6C, D, S7C, S8C, D, S9C, D, S11D). This pattern is not ubiquitous, and some continuous blue layers are also present within the spherulitic growth zones (Fig. S6C, D, 7C, 8C, S11C, D). Blue luminescence is also prevalent in areas where adjacent cameral deposits left a small gap, e.g., where proximal and distal deposits are in near contact (Fig. S8D) or where there is a small gap between spherulitic sectors (Fig. S8C, S9C, D). The sector boundaries of the mural delta in PZO 16538-01 display predominantly bluish luminescence colours (Fig. 2F, S11D).

### Scanning electron microscopy

The EBSD (electron backscatter diffraction) phase maps point to the aragonitic mineralogy of the outer shell wall and the plano-mural deposits, while the spherulitic sectors consist mainly of calcite (Fig. 4A, E). However, the EBSD signal was relatively weak, with overall indexing rates of 73.1% and 52.9% for the cross section PZO16535-01 and the longitudinal section PZO16548-02, respectively. See Appendix S1 for additional details.

**Fig. 4.**
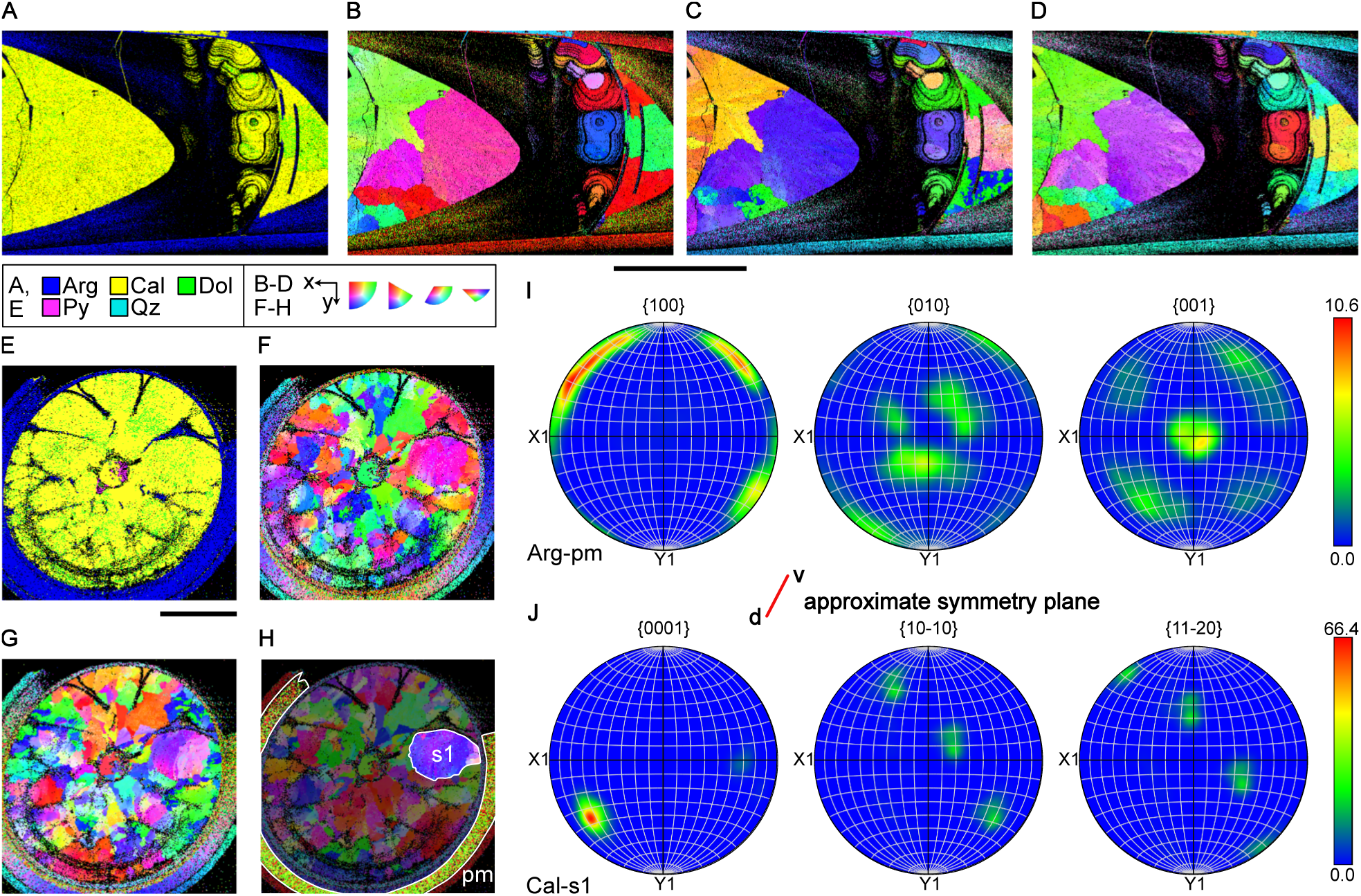
EBSD results of *Trematoceras elegans*. A-D, PZO 16538-02, longitudinal section. E-H, PZO 16535-01, cross-section. I-J, pole figures corresponding to E-H, with added approximate symmetry plane (d = dorsal, v = ventral). A, E, mineral phase maps, see colour legend for identified mineralogies. B-D, F-H, Inverse pole figure (= IPF) maps for the X, Y and Z axes for both samples. Similar colours indicate similar crystal orientations, see legend. I, aragonite pole figures for the pm subset in H, showing the distribution of orientations in the a-axis 100, b-axis 010 and c-axis 001, irrespective of location within the sample. Note strong co-orientation with shell wall (Fig. S23D). J, calcite pole figures for the ss subset in H, showing the distribution of orientations in the c-axis 0001, a_1_-axis 10-10 and a_2_-axis 11-20, irrespective of location within the sample. Note strong alignment within spherulite. Abbreviations: Arg =aragonite, Cal = calcite, Dol = dolomite, Py = pyrite, Qz = quartz, pm = proximal deposits, ss = spherulitic sector. See, e.g., Schwartz et al. (2009) or Cusack (2016) for further details on EBSD and interpretation of figures. All scale bars represent 1 mm.

In the longitudinal section PZO 16538-02, the a-axes of the aragonite crystals of the plano-mural deposits are aligned with those of the shell wall towards the centre of the conch, while the b- and c- axes each align in three preferential orientations, one of which is co-oriented with the shell wall in each axis (Fig. 4B-D, S23D, F). Thus, these fabrics are aligned with the shell wall but are partly rotated in their b- and c-axes. This pattern is consistent between the two chambers in the sample

The aragonite crystal orientation of the cross-section PZO 16535-01 is consistent with the pattern observed in the longitudinal section PZO 16538-02. In this specimen, the shell aragonite crystals are radially arranged and the a-axes directed towards the centre of the conch (Fig. 4F-H). This arrangement is also reflected in the aragonite pole figures (Fig. 4I), where the a-axes show strong co-orientation with the shell wall, while the b- and c-axes indicate rotation (Fig. S24D). Between the shell wall and the septum, the IPF (= inverse pole figure) maps reveal that there are two distinctly oriented aragonite phases, corresponding to the plano-mural (parallel to shell wall) and plano-proximal deposits (parallel to septum). The latter are only visible as a thin band on the dorsal side and differ in their orientation that is more similar to the septum but show a single central orientation maximum (i.e., longitudinal direction of the conch; Fig. S24F) instead a ring-like orientation distribution as in the septum (Fig. S24E). Note that the symmetry of the pole figures is impacted by the missing ventral shell and some parts of the specimen not fitting into the observation window.

The spherulitic portions of the cameral deposits that displayed orange luminescence colours were identified as calcite with a significantly increased hit rate (i.e., fewer “unidentified pixels”; Fig. 4A, E). In the longitudinal section PZO 16538-02, the IPF maps revealed a uniform c-axis orientation within the calcitic spherulite sectors but also showed a pronounced change in orientation compared to the adjacent sector (Fig. 4B-D). Furthermore, the orientations of the crystals within the dorsal and ventral spherulites are congruent with each other, i.e., their crystal axes are parallel. The pole figures of individual calcite spherulites indicate that the c-axes are aligned, while there are three major orientations for the a_1_- and a_2_-axes (Fig. S23G-J). The spherulites are more or less parallel to the septum (Fig. S23E), except for the spherulite close to the centre of the conch, of which the c-axes are aligned perpendicular to the sample plane (Fig. S23G). Thus, the latter spherulite is rotated by 90° compared to the other spherulites, which is consistent with the radial arrangement of the spherulitic sectors. Notably, the c-axes of the other spherulites (Fig. S23H-J) appear to be more or less co-oriented with the c-axes of the aragonitic plano-mural deposits (Fig. 23F) and the shell wall (Fig. 23D).

Reconstructing the calcite c-axis orientations is more complicated in the cross-section PZO 16535-01 (Fig. 4F-H). Individual spherulites display the same orientation pattern with c-axes parallel to the septum, directed towards the centre of the conch (Fig. 4J). This pattern is less obvious but still present in other spherulites (Fig. 24I). The smaller calcite spherulites on the dorsal side display several c-axis orientations that roughly align with the symmetry axis (Fig. S24H). Smaller patches of aragonite between the calcite spherulites have a-axes that are more or less perpendicular to the adjacent calcite spherulite (Fig. S24G).

When elemental compositions retrieved via EDS are compared, the calcitic deposits contain higher amounts of Mg (Fig. 5E), Mn (Fig. 5C) and Fe (Fig. 5D). In contrast, the aragonitic deposits are enriched in Sr (Fig. 5E) and contain slightly higher Ca concentrations (Fig. 5A).

**Fig. 5.**
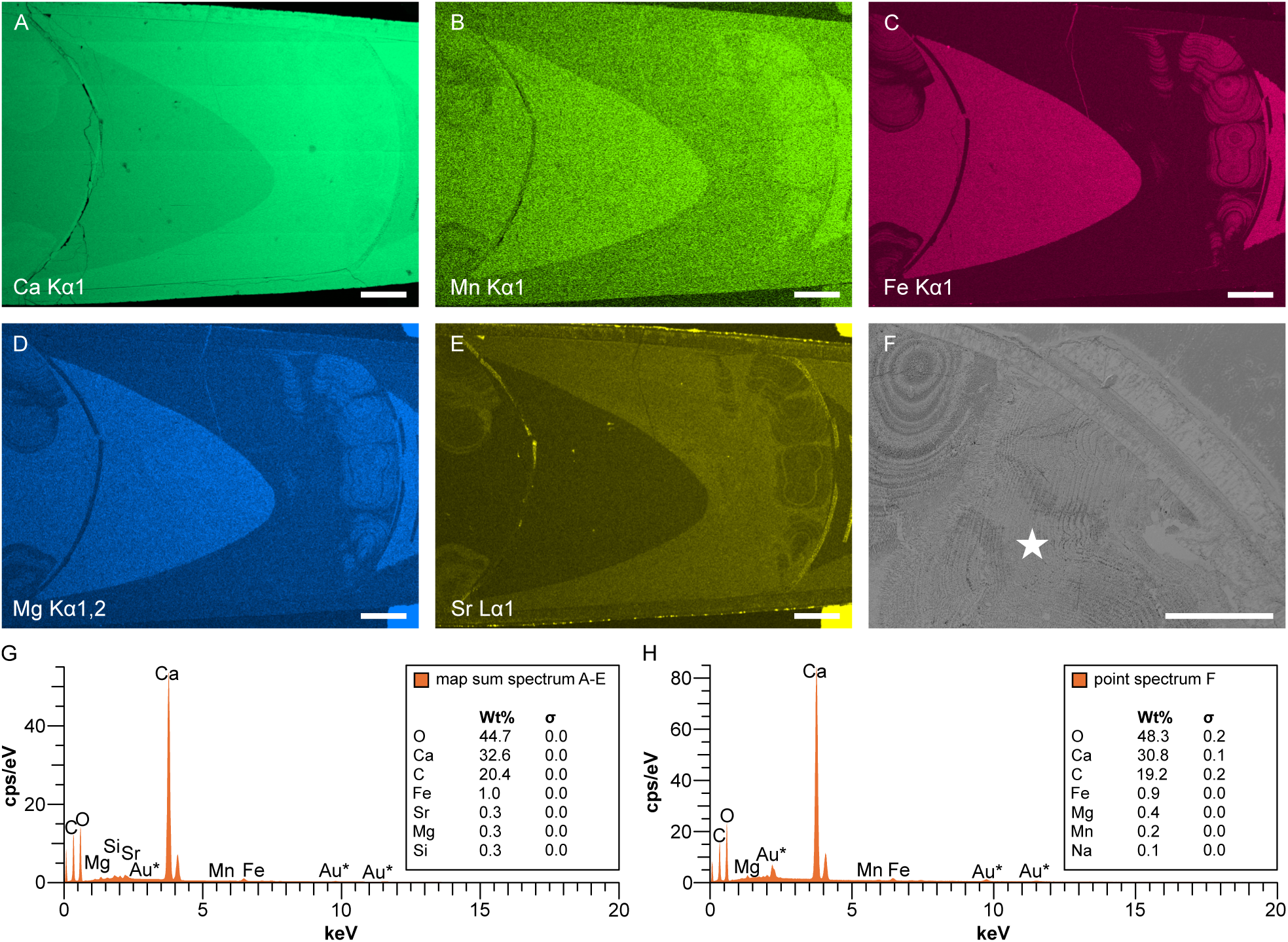
Element distribution in *Trematoceras elegans* based on EDS analysis. A-E, PZO 16538-02, longitudinal section. Element maps are modified by contrast and brightness enhancement to show differences better. A, calcium, note that horizontal stripes represent an artifact. B, manganese. C, iron. D, magnesium. E, strontium. F, PZO 16540-01, cross-section. SEM image, showing the position of point measurement in (H). G, map sum spectrum of PZO 16538-02 for (A-E) and Fig. S19. Corresponds to calcium carbonate with low amounts of iron, strontium, magnesium and silicium. All other elements have lower weight percentages (Wt%), see Fig. S19. H, point spectrum of specimen PZO 16540-01 (see F for position), showing essentially an identical composition to the map sum spectrum of PZO 16538-02. For elemental maps of PZO 16540-01, see Fig. S20. All scale bars represent 250 µm.

The elemental composition of the outer shell wall is almost indistinguishable from the aragonitic cameral deposits, except for a higher concentration of Sr in the latter (Fig. 5E). Conversely, the septa are enriched in Sr (Fig. 5E) but have an otherwise identical element distribution to the outer shell wall. The originally empty part of the chamber (now filled by blocky, diagenetic calcite) and the calcitic deposits have similar elemental concentrations, but the diagenetic calcite contains higher concentrations of Fe (Fig. 5D), Mn (Fig. 5C), and Mg (Fig. 5E).

Under the SEM, the excellent preservation of the shell material is exemplified by the characteristic cephalopod nacre tablets of the shell wall (Fig. 6I, S16A, B), which are typically rapidly altered during diagenesis. The microstructure of the aragonitic cameral deposits is fibrous, with crystal diameters between about 0.5-2.0 µm (Fig. 6B, C, 7B-D). Crystal dimensions correlate with the EBSD detection rates, i.e., the smallest aragonite crystals coincide with the lowest detection rates (Fig. 6A-D). The calcite crystals within the distal spherulites have a similar microstructure as the aragonite but tend to be less porous and, within a range of sizes, to be larger in diameter (Fig. 6D-E, 7E). Differences in crystal size and elongation direction are well visible at the boundaries between spherulitic sectors and growth increments (Fig. 6F, H, 7F-I). Lateral and vertical aragonite-calcite transitions occur as changes in microstructure, the former gradually (Fig. 6H, 7F, G) and the latter more abruptly. Locally, sudden changes in the direction of crystal elongation are noted (Fig. 6F, 7H, I, K, L, S13).

**Fig. 6.**
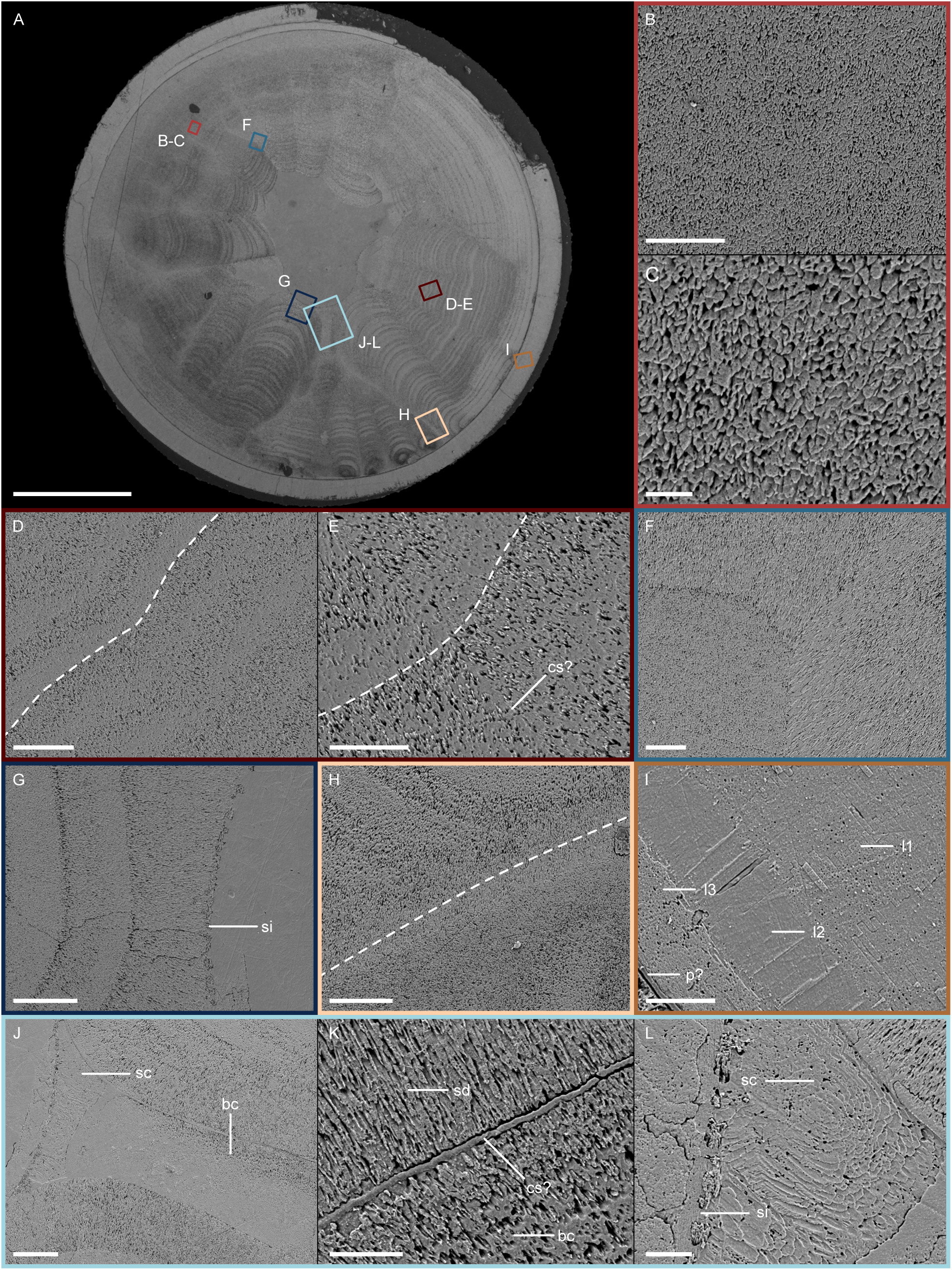
SEM image of *Trematoceras elegans*, PZO 16538-01, cross-section. A, entire specimen, with position of images (B-L) indicated by coloured rectangles. For a high-resolution version of this figure, see Figure S12. B-C, aragonitic cameral deposits, porous fibrous microstructure. D-E, calcitic cameral deposits, with alternating, less porous fibrous microstructure, example given by dashed line. Note the discontinuity perpendicular to the growth increments (potential cameral sheets = cs?). F, aragonite (bottom left) to calcite (top left, right) transition. Note the well-defined boundary between the different grain elongation directions, forming a Y-shaped pattern. G, contact area between cameral deposits and connecting ring, the latter blocking further growth. H, detail of mural delta. Note the mixture of different crystal sizes and porosities, in addition to the sharp transitions. I, detail of shell wall. Note the three layers of the shell: outermost nacre tablets (l1), middle vertical pillars (l2), and innermost small nacre tablets (l3 = septum?). The thin layer in the bottom left corner possibly represents the pellicle (p). J-L, Biconcave structure (bc) ventrally of siphuncle, with dorsal area displaying scale-like, concentric microstructure (sc). K, Detail of boundary (potential cameral sheets = cs?) between the inner side of biconcave structure (bc) and spherulo-distal deposits (sd). Note the apparent continuation of growth for a short distance and subsequent transition into lower porosity. L, detail of dorsal area of biconcave structure, directly adjacent to siphuncle (si). Note the scale-like, concentric structure. Scale bars represent: 2 µm (C); 5 µm (K, L); 10 µm (B, E, F); 20 µm (D, G-J); 500 µm (A).

**Fig. 7.**
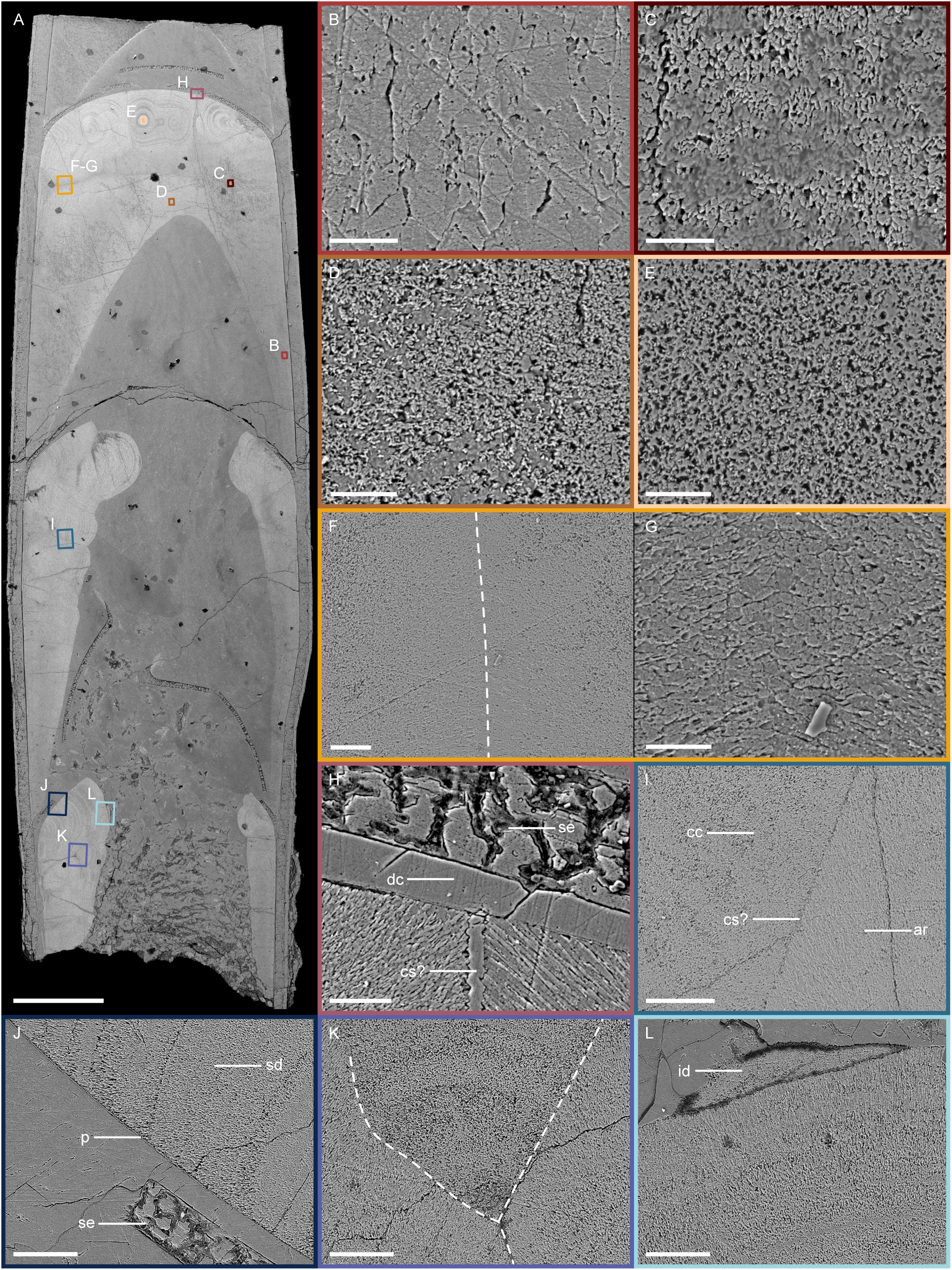
SEM image of *Trematoceras elegans*, PZO 16538-02, longitudinal section. A, entire specimen, with the position of images (B-L) indicated by coloured rectangles. For a high-resolution version of this figure, see Figure S14. B-D, aragonitic cameral deposits, porous fibrous microstructure, with variable crystal size. E, calcite cameral deposits, less porous fibrous microstructure. F, lateral aragonite (middle) to calcite (left, right) transitions close to cameral sheet. Note the well-defined boundary between the different direction of grain elongation in the middle (dashed line), and gradual transitions between different crystal sizes. G, close up of boundary between aragonitic crystals in the opposite grain elongation direction (dashed line in F). H, detail of septum-cameral deposit contact. Note that the cameral deposits display distinct directions of the elongations of grains and a well-defined boundary (cs?), likely slightly enlarged through a crack. The septum is slightly detached from the cameral deposits, with diagenetic calcite in between (dc). I, detail of calcite (cc) to aragonite (ar) transition. Note the well-defined boundary between differing grain elongation directions (cs?). J, spherulo-distal deposits (sd) detached from septum (se). Note the thin structure lining cameral deposits on the lower side, possibly representing the pellicle (p). K, the boundary between three separate growth sectors of different grain elongation directions (dashed lines). L, overgrowth of spherulite by isolated, irregularly shaped deposit (id), possibly secondary. Scale bars represent: 5 µm (B-E, G); 10 µm (F, H); 20 µm (I-L); 500 µm (A).

The growth increments of the spherulo-distal deposits are visible as wavy, somewhat irregular alterations between layers of lower and higher porosity in the aragonite or calcite (Fig. 6D, E, 8C). Specifically, zones of lowest porosity border what seem to be zones of highest porosity (examples are given as dashed lines in Fig. 6D, E). The growth layers are well-correlated across the boundaries between the spherulites, a feature that suggests that they formed *in tandem* (Fig. 8A, B). The boundaries, as such, are clearly defined, even at high magnifications (Fig. 8). There appears to be no material difference between the deposits and the boundaries, apart from a small discontinuity of approximately 1-2 µm thickness (Fig. 8C, D). In addition, the crystals immediately next to the boundary appear to consist of an admixture of both calcite and aragonite, while layers further away are devoid of aragonite fibres (Fig. 8C, D). Distinct structures separating adjacent deposits are only visible in a few cases. Besides the biconcave area below the siphuncle (Fig. 6K) and between spherulites (Fig. 7H), such sheet-like structures are also visible on the septal surface of the cameral deposits. Possibly, the latter sheets represent the pellicle (Fig. 7J).

**Fig. 8.**
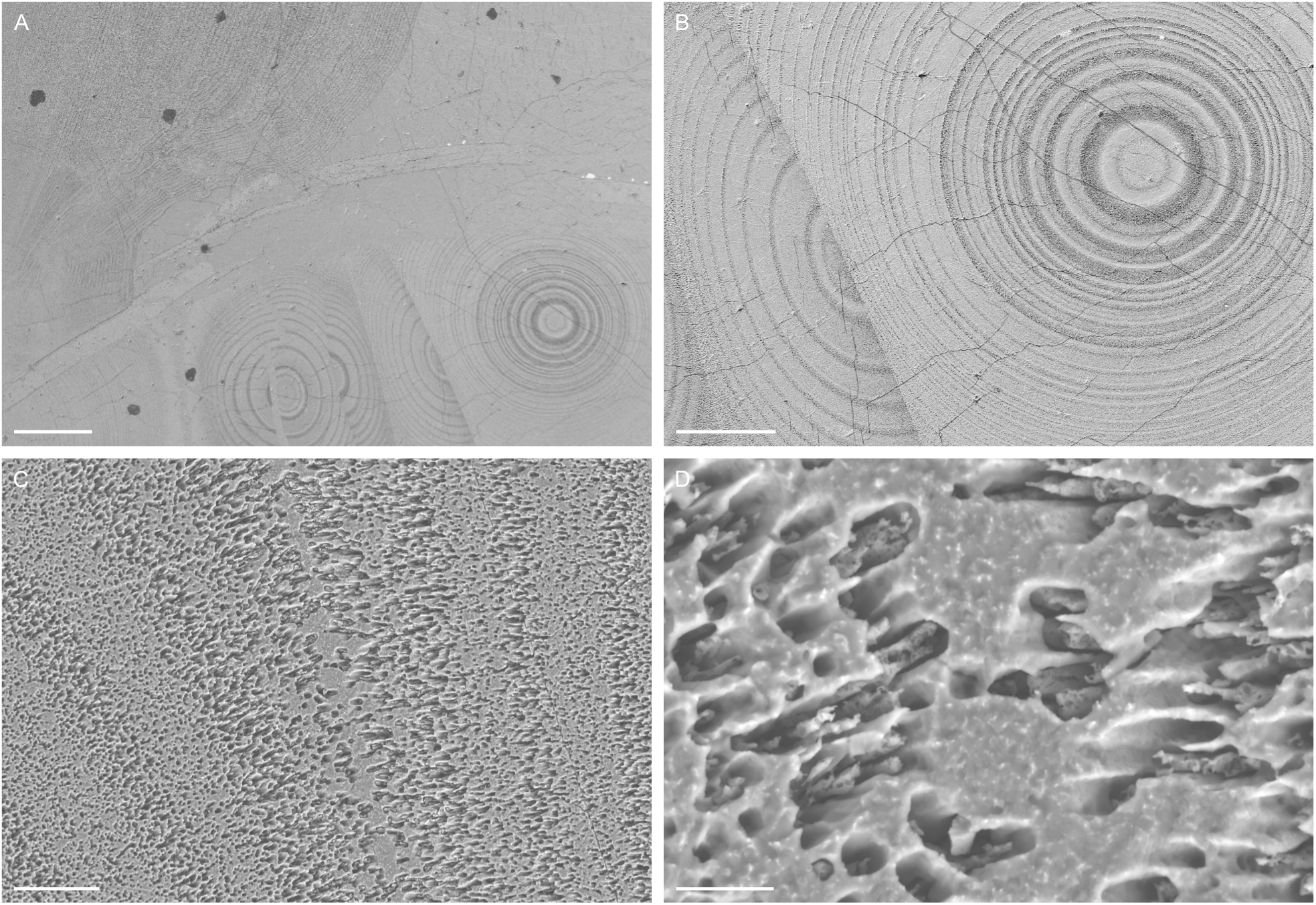
SEM images of potential cameral sheets in *Trematoceras elegans*, PZO 16535-02, longitudinal section. A, overview of four distal spherulitic sectors with well-defined boundaries (bottom), a partly broken septum (middle) and plano-mural deposits of the previous chamber (top left). B, a single spherulitic sector, with a well-defined boundary to a neighbouring sector (interpreted as a cameral sheet). Note that the growth layers can be precisely correlated across the cameral sheet. C, close up of growth increments close to cameral sheet. Note the changes in porosity between growth increments and close to cameral sheet. D, close up of cameral sheet. No distinct structure is discernible, but aragonite fibres terminate adjacent to cameral sheet. Scale bars represent: 200 µm (A); 100 µm (B); 10 µm (C); 2 µm (D).

## Discussion

### Interpretation of structures

The cameral deposits reported here show a bilaterally symmetric organisation in alignment with the symmetry plane of the conch. Bilateral symmetry of cameral deposits has been shown previously in various groups of orthoceratoids from the Ordovician (Evans *et al*. 2012; Aubrechtová *et al*. 2020), Silurian (Kolebaba 1974, 1975, 1999a, *b*, 2002), Devonian (Flower 1936, 1939; Stanley & Teichert 1976; Niko 1991; Pohle & Klug 2018) and Carboniferous (Fischer & Teichert 1969; Crick 1982; Blind 1991; Seuss *et al*. 2012a). Note that while bilateral symmetry is common to all these examples and, therefore, likely a general feature of cameral deposits, there are clear morphological differences that likely reflect taxonomy. Whether the morphology is informative on species level or more relevant for the comparison between higher taxa is currently unknown, although Fischer & Teichert (1969) reported the morphology to be consistent within the same genus.

In *Trematoceras elegans*, the distal deposits are spherulitic and transition into plano-mural deposits that thin out towards the apertural end of the chamber. Proximal deposits appear to be less common in *Trematoceras*, although this may be an ontogenetic effect. The specimens in our sample with proximal deposits were later ontogenetic stages. One of these specimens had more than 150 growth increments of plano-mural deposits (PZO 16535-02, Fig. S18A). In contrast, an ontogenetically earlier stage had only about 35 growth increments in the same position but lacked proximal deposits entirely (PZO 16538-02, Fig. S18C). Thus, spherulo-proximal deposits are likely ontogenetically delayed, as shown by Crick (1982) in Carboniferous taxa.

Aragonite has long been proposed to form the original mineralogy of cameral deposits (e.g., Flower 1955b). Stehli (1956) was the first to suggest that cephalopod cameral deposits from the upper Carboniferous Buckhorn Asphalt deposit (Boggy Formation) consisted partially of primary calcite, lacking evidence for diagenetic alteration. Conversely, several authors rejected the possibility of primary bimineralic cameral deposits based on the loss of microstructure in the calcitic areas (Fischer & Teichert 1969; Crick & Ottensman 1983). A more recent study reported the presence of primary high-Mg calcite areas in the cameral deposits of Buckhorn Asphalt orthoceratoids (Seuss *et al*. 2012a, *b*).

Some parts of our specimens show evidence of diagenetic alteration, e.g., orange luminescence in patches of the outer shell (e.g., Richter *et al*. 2003) or non-aligned crystal orientation in the calcite spherulites (e.g., Pederson *et al*. 2020). However, the microstructures are generally well preserved, in particular the nacre tablets in the shell wall (Fig. 6I, S16B), which together with their blue luminescence (Fig. 2) indicates little diagenetic alteration. The calcite crystal orientations revealed by EBSD maps show consistent patterns within the same spherulite, between neighbouring spherulites and between successive chambers. The consistency of these patterns supports the primary calcitic mineralogy of the spherulites. The calcite in our specimens displays orange luminescence colours, which is commonly taken as evidence for diagenetic alteration of biominerals, caused by the presence of Mn (e.g., Richter *et al*. 2003). However, this notion is complicated by the fact that some recent organisms also incorporate Mn^2+^ cations into their biominerals, causing orange luminescence (e.g. Barbin 2013; Stevens *et al*. 2017) and the fact that kinetic effects such as growth rates induce differences in cation incorporation (Ten Have & Heijnen 1985). The spatial distribution of calcite and aragonite is consistent across chambers. This implies that the spherulo-distal deposits are primarily calcitic, while the plano-mural deposits are mainly aragonitic in mineralogy. A far more irregular fabric is expected for the case of aragonite-to-calcite neomorphism. That said, fabric-retentive neomorphism has been reported (see Mueller *et al*. 2024 and references cited therein). Further support for the primary calcitic composition of the spherulites is the fact that they are repeatedly overgrown by primary aragonite (blue luminescence); if the calcite were a diagenetic feature, this spatio-mineralogical relation would be difficult to explain when considering the different solution kinetics of these minerals. The preservation of biogenic aragonite in the outer shell provides further support to this model.

Earlier studies apparently assumed that cameral deposits would be homogeneous structures with uniform mineralogy (Fischer & Teichert 1969; Crick & Ottensman 1983). The combined EBSD and EDS data shown here document that the mineralogy and ultrastructure are far more variable and complex (Fig. 4, 5). As the spherulitic layers show changes in Mn, Fe, Mg and Sr concentrations, it is conceivable that the changes in porosity seen under the SEM (Fig. 6D, E, 8B, S13C, D, S16C, D) represent transitions between aragonite and low-Mg calcite. These findings also have consequences for the reported elemental concentrations of Crick & Ottensman (1983). The values documented in their study reflect the aragonitic parts of the cameral deposits but are not representative of the entire cameral deposits (Fig. 5).

In spite of these general patterns, irregularities exist both in the symmetry of the deposits and their composition. Some growth layers deviate from the predominant aragonite-calcite distribution pattern, with lateral (i.e., within the same growth layer) and vertical (i.e., between growth layers) mineral transitions. These transitions are reminiscent of speleothems, where differences in the fluid Mg concentration cause similar changes. When threshold limits are crossed, i.e., abundant calcite precipitation shifts the Mg-to-Ca ratio of the fluid, the thermodynamically preferred mineralogy is aragonite (e.g., Wassenburg *et al*. 2012, 2016; Hashim & Kaczmarek 2021).

The perhaps most significant observation made here is that the delimitations between the radial spherulitic sectors are well-defined. The boundaries have a dark colour under transmitted light and their arrangement follows a conspicuous, more or less bilaterally symmetric pattern (Fig. 2). Some of these dark lines delimit the outer surface of the spherulites, in contact with diagenetic cement (Fig. 2A, 3E, S9). This feature is taken as strong circumstantial evidence of radially branching cameral sheets that were present prior to the deposition of the cameral deposits. The original, arguably organic structures are not preserved, as there is no clear microstructure visible apart from a small discontinuity between the spherulites. Furthermore, where some sheet-like structures are seemingly preserved (Fig. 6K, 7H, J), their elemental compositions match those of the diagenetic blocky calcite within the chambers, containing higher Fe and Mg concentrations.

Two main arguments are in favour of the presence of cameral sheets during the lifetime of the animal, prior to the formation of the cameral deposits: (1) the growth layers can be precisely correlated across the cameral sheet, a feature that points to contemporaneous precipitation (e.g., Fig. 8A, B); and (2) the well-defined boundaries indicate that a (arguably thin) planar sheet must have been present that delimitated crystal growth. In the absence of such compartmentalisation, i.e., the cameral sheets, the boundary between sectors is expected to display a more irregular shape, including an undefined transition zone where the direction of grain elongation changes (compare, e.g., microbially induced spherulites, Chafetz *et al*. 2018; spindles in ooids, Mono *et al*. 2025). An example of how a natural spherulite unconstrained by cameral sheets might look is shown in Figure S7B, D, E. There, post-secretion-alteration obliterated the original structure, resulting in an irregular fabric. Co-occurring stromatolites or corals from the St. Cassian Formation display less regular growth without these narrow delimitations between spherulites and different microstructures (e.g., Fürsich & Wendt 1977; Wendt 1993; Sánchez-Beristain & Reitner 2012).

No evidence of organic remains between the cameral deposits (e.g., in the form of phosphorus or sulphur, see Fig. S19C, S20G, H) is preserved. This implies that the organic sheets themselves decayed *post-mortem* and diagenetic cement filled the narrow interspaces left after their decay. This is visible as the narrow discontinuities between sectors mentioned before. The sheets were likely very thin, akin to the pellicle, a thin (1-5 µm) organic lining inside the chambers of modern *Nautilus* (Ward 1987). In the orthoceratoids from the Buckhorn Asphalt, organic remnants were reported and interpreted as parts of the pellicle that broke off and were trapped between the cameral deposits (Grégoire & Teichert 1965; Grégoire 1988). In our view, this interpretation seems scientifically unsatisfactory, as it is unclear how the fragments would detach from the septum and end up in between the growth layers of the cameral deposits. In conclusion, the combined evidence shown here points to the presence of very thin, organic cameral sheets with a poor preservation potential prior to the formation of the cameral deposits.

Potentially homologous structures are known from well-preserved ammonoids. These structures have been referred to as “conchiolin lamellae” (e.g., Schindewolf 1967), “cameral membranes” (e.g., Landman *et al*. 2006; Polizzotto *et al*. 2015; Mironenko & Smurova 2024), “intracameral conchiolin layers” (Weitschat 1986; Weitschat & Bandel 1991) or “intracameral sheets” (e.g., Checa 1996). Here, the term “cameral sheets” is preferred. This is because “membrane” carries a functional interpretation, and the prefix “intra” is unnecessary because “extracameral” sheets do not exist. Given that ammonoids originate from orthoceratoids (Engeser 1996; Kröger *et al*. 2011; Klug *et al*. 2015; Hoffmann *et al*. 2022), we anticipate that cameral sheets are more widespread within ectocochleate cephalopods. Thus, cameral sheets may represent an autapomorphy of the clade combining ammonoids and orthoceratoids with the Coleoidea. Similar structures in orthoceratoids are currently only known in the Lamellorthoceratidae, a short-lived (Early-Middle Devonian) group of specialised orthoceratoids (e.g., Stanley & Teichert 1976; Bandel & Stanley 1989; Niko 1991; Cichowolski & Rustán 2020). We propose that the lamellae within their chambers were homologous to the cameral sheets described here and reported in ammonoids.

The fact that cameral sheets appear to be much more regularly preserved in lamellorthoceratids may be explained by their heavier calcification. The lamellae also appear to be more numerous than cameral deposits in other orthoceratoids. Further evidence for cameral sheets in orthoceratoids is represented by alleged “vascular imprints” on the apertural and apical surfaces of septa in Ordovician lituitids (Sweet 1958; Aubrechtová & Meidla 2020; Aubrechtová & Korn 2022) and the Silurian cameral deposit-bearing taxa *Leurocycloceras* and *Sphooceras* (Flower 1941; Holland 1964; Turek & Manda 2012). We argue that these structures on the apical septal surfaces are not vascular imprints of a “cameral mantle” but rather traces of the attachments of the organic cameral sheets. The presence and form of cameral sheets in orthoceratoid cephalopods may also be reflected by “pseudosepta” (e.g., Holm 1885; Schröder 1888; Remelé 1890; Flower 1939; Teichert 1964); these are visible on longitudinal sections as diagonal, crenulated structures developed along the plane of contact of thick proximal and mural deposits. In some orthoceratoid specimens, a thin layer of dark matter was reported along the “pseudosepta”, as well as on the surface of the cameral deposits and within them, between individual growth increments (e.g., Aubrechtová & Meidla 2020). All the above structures within orthoceratoid phragmocones are consistent with an interpretation as cameral sheets, though further studies are needed to confirm this hypothesis. Note that orthoceratoid “pseudosepta” must not be confused with pseudosepta and pseudosutures known from ammonoids (Polizzotto *et al*. 2015). If the orthoceratoid “pseudosepta” are tentatively considered as cameral sheets, this confusion can conveniently be avoided. To summarise the evolutionary perspective, cameral deposits were, in the author’s view, a general feature of the Orthoceratoidea. They originated in the Dissidoceratida and are present in all orthoceratoids except for the Rioceratida, which is considered ancestral to all other orthoceratoids (King & Evans 2019; Pohle *et al*. 2022a). The origin of cameral sheets is difficult to pinpoint because of their limited preservation potential and, hence, the fossil record, requiring either excellent preservation (as in ammonoids), calcification of the sheets (as in lamellorthoceratids) or circumstantial evidence, as in the cameral deposits studied here.

Despite this patchy record, the currently known taxonomic distribution of cameral deposits is in line with the hypothesis presented here (Fig. 9). It appears that cameral deposits and sheets were lost in the Coleoidea. However, a connection may be drawn to the horizontal and vertical membranes within the chambers of *Sepia* (Checa *et al*. 2015). The microstructure of the pillars crossing these sheets is remarkably similar to the cameral deposits (Checa *et al*. 2015). Likewise, the so-called adapical ridge in *Spirula* (Lemanis *et al*. 2020; Checa *et al*. 2022; Griesshaber *et al*. 2023) or the infillings between the septum and shell wall of *Nautilus* (Grégoire 1962, 1987; Mitchell & Phakey 1995) bear some resemblance to the cameral deposits in their microstructure. It is, therefore, possible that remnants of cameral deposits and sheets are more widespread within cephalopods than currently thought. Upcoming work will need to clarify whether all these structures are homologous or analogous.

**Fig. 9.**
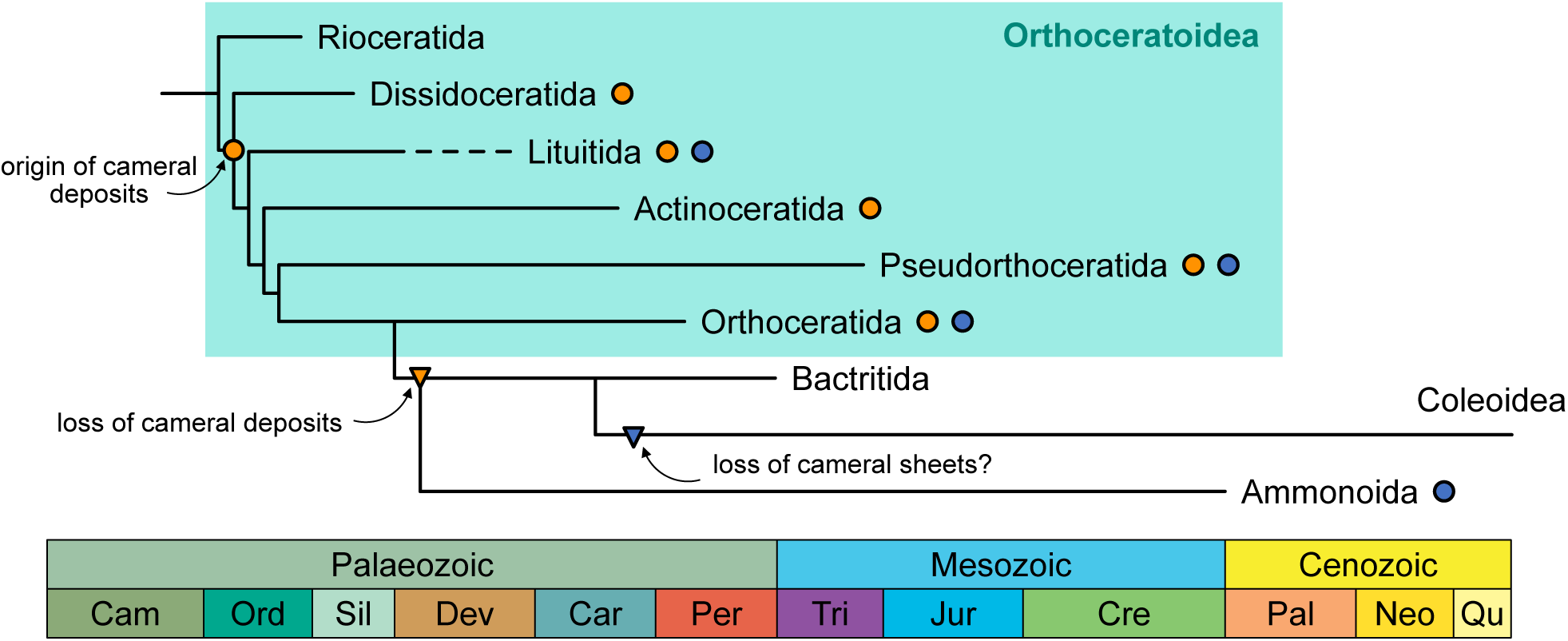
Phylogenetic distribution of cameral deposits and cameral sheets. Orange circles indicate presence of cameral deposits within the group; blue circles correspond to known cases of cameral sheets. Note that the exact timing of origin and loss of cameral sheets is difficult to assess due to the low preservation potential. Phylogeny after Kröger et al. (2011), Pohle et al. (2022a) and Hoffmann et al. (2022).

### Previous growth models

For any growth model to be accepted, it has to be able to predict the observed morphological structures of the cameral deposits, including the variability seen within a single chamber, during ontogeny and between taxa – not only within a single study, but also considering previously published observations. The consistency in the basic patterns of cameral deposits and their variability (e.g., Flower 1955; Fischer & Teichert 1969; Dzik 1984) suggests a common formation mechanism, and likely a single evolutionary origin. We discuss previously proposed hypotheses and compare the predicted morphologies to the observed patterns. Due to the similarities in the bauplan of the recent *Nautilus* and all extinct ectocochleate cephalopods, it is undisputed that the function of the phragmocone is identical, i.e., to provide buoyancy (e.g., Denton 1974; Westermann 1975; Crick 1988; Peterman *et al*. 2019; Peterman & Ritterbush 2022). Therefore, we further constrain these models in that the shell must be able to maintain neutral buoyancy throughout its growth. This assumption is supported by the frequent occurrence of cameral deposit-bearing orthoceratoids within distal black shale facies (Kröger *et al*. 2009). If a growth model does not meet both requirements, it is therefore rejected.

Under the *post-mortem* growth model (Mutvei 1956, 2002, 2018), the buoyancy requirement is met, as the living animal would never contain any cameral deposits and, therefore, would rely entirely on cameral liquid as a counterbalance to the gas-filled chambers, as in *Nautilus* (e.g., Ward 1979; Greenwald *et al*. 1980; Ward *et al*. 1980). However, as has been pointed out repeatedly (e.g., Flower 1964; Fischer & Teichert 1969; Crick 1982; Seuss *et al*. 2012a; Pohle & Klug 2018), the complex symmetrical structure of the cameral deposits cannot be explained by *post-mortem* processes. Although Mutvei (2018) accepted the biogenic microstructure of the cameral deposits, he insisted on their *post-mortem* formation, proposing calcifying bacteria as the culprits. This growth model still cannot explain the observed structures beyond their laminated appearance. Any *post-mortem* deposits would grow independently of the symmetry plane of the conch and would not show a regular progression from thicker deposits at the apex towards thinner deposits in apertural chambers, with the first few chambers adjacent to the body chamber being devoid of cameral deposits (Crick 1982). Instead, this model would produce more randomly distributed and oriented deposits (Fig. 10A). Another argument is the complete absence of cameral deposits in co-occurring ammonoids (Fig. S4), and corresponding structures have never been reported in other molluscs, such as gastropods, in which similar *post-mortem* conditions must have been present. Indeed, cameral deposits appear to be entirely restricted to orthoceratoids, and the few exceptions either need to be reevaluated taxonomically (King & Evans 2019) or possibly reflect actual *post-mortem* structures, as they have not yet been shown to possess the same laminated, bilaterally symmetric structure (e.g., Flower & Teichert 1957; Jeletzky 1966). Therefore, this model can be firmly refuted. We point out that this assessment does not exclude the existence of secondary *post-mortem* deposits – these are real taphonomic processes, and care must be taken not to confuse secondary with primary cameral deposits. Guidelines to distinguish the two have been already established by Teichert (1933, 1964). As shown here, true cameral deposits are further characterised by their complex, symmetrical and regular ontogenetic patterns, in addition to the sharp delimitation between growth sectors by the cameral sheets.

**Fig. 10.**
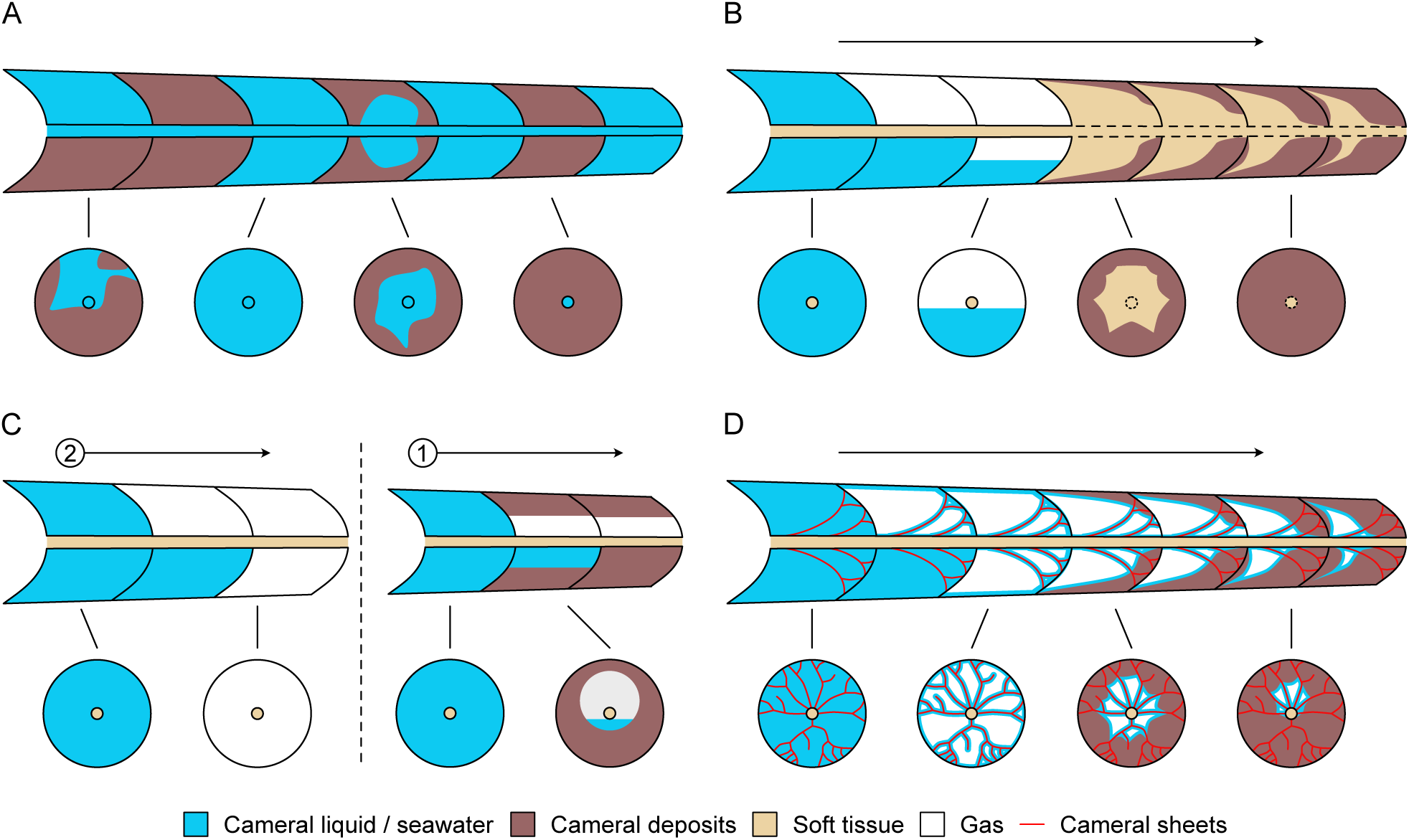
Growth cycle models of cameral deposits in orthoceratoids. Ontogenetic progression of cameral deposits is shown by tracing the state of the chambers in adapical direction (arrow, numbers represent different ontogenetic stages). (A-C) represent previous hypotheses, and (D) is the hypothesis proposed herein. Arrows represent ontogenetic sequence of cameral deposit formation. A, post mortem (e.g., Mutvei 1956, 2018): Cameral deposits are secondary types of cements, deposited inorganically or microbially mediated and grow independently of the ontogeny of the shell and orientation. B, cameral mantle (e.g., Flower 1955b; Teichert in Fischer & Teichert 1969): Chambers are first emptied, subsequent syn-vivo destruction of siphuncle and growth of cameral deposits into chambers to directly control precipitation of cameral deposits. C, cameral liquid (e.g., Fischer in Fischer & Teichert 1969; Blind 1991): Precipitation from liquid retained in the chambers. Note that this model requires a change in the precipitation activity (phases 1 and 2) and is not buoyant in early ontogenetic stages. D, cameral membranes (this study): Organic sheets are formed when the soft body vacates the chamber, increase the functional surface area of the siphuncle and guide the precipitation of cameral deposits.

The cameral mantle hypothesis is appealing as it can explain the symmetrical arrangement of cameral deposits without difficulties (Flower 1955b; Fischer & Teichert 1969; Fig. 10B). In principle, this model could result in any morphology, as a soft tissue provides precise control on the shape of the biominerals (Immenhauser *et al*. 2016). However, it appears that if the animal were able to control the cameral deposits directly by a cameral mantle, they would likely display microstructures typical for biologically controlled biominerals of cephalopods (e.g., nacre). Our specimens, as well as countless previously investigated sections, unequivocally show that the deposits grew inwards from the outer shell wall and the septum, starting at distinct nucleation sites with subsequent spherulitic growth. If this model were correct, the surface of the growth increments must have been completely covered by the secreting mantle tissue. This would mean that the actively biomineralizing surface would first cover only isolated patches of the septum but be continuous along the outer shell wall and would later increase and decrease in a non-trivial pattern. The cameral mantle would eventually need to shrink to be replaced by cameral deposits. Crick (1982) argued that there would always be a small gap between the deposits and the siphuncle caused by the remnants of the cameral mantle, which is in disagreement with the observation that chambers that are completely filled by cameral deposits exist. The entire mechanism appears highly inefficient, and it seems more likely that if it were possible, the cameral mantle would precipitate cameral deposits uniformly along the shell wall and septa (i.e., planar deposits only); yet this kind of morphology is never observed during the long geological record of orthoceratoids. A good example of such morphology is provided by the belemnite rostrum, which was deposited directly by the mantle and consists of comparatively simple growth increments without major predominant irregularities and spherulitic areas (Hoffmann & Stevens 2020). Vice versa, biologically controlled cephalopod shell material never exhibits the micro- and macrostructure observed in cameral deposits. The complex anatomy of the cameral deposits is also not necessary to fulfil the presumed function of balancing the shell.

A further caveat of the cameral mantle hypothesis is that this model requires the supply of cameral tissues with blood. As pointed out by Mutvei (2018), the siphuncle is closed off towards the chambers by the siphuncular epithelium, without perforations allowing the passage of blood vessels. Thus, the transport of entire blood cells through the siphuncle is impossible, unless a partially open circulatory system is postulated, which involves numerous additional assumptions. Any exchange of molecules between siphuncle and chamber must happen through cellular transport proteins. Considering that the molluscan hemocyanin, which is responsible for oxygen transport in all living cephalopods, is among the largest known proteins (Kato *et al*. 2018), extracellular transport would be highly unlikely, if at all possible. In addition, if blood could be supplied through the connecting ring as implied by Flower (1955b, 1964) and Teichert (Fischer & Teichert 1969), it would defy the osmotic function of the siphuncle. The only solution to this problem would be partial resorption of the siphuncle during the life of the animal (see, e.g., Kolebaba 1974, 1999a, 2002; Marek 1998; Zhuravleva & Doguzhaeva 2004).

However, in most orthoceratoids, including the specimens documented here, the connecting ring is completely preserved in the same chamber as the cameral deposits, meaning that this hypothesis could only explain a few exceptional cases, if at all. In fact, it is difficult to prove whether the connecting ring was destroyed *syn vivo* or whether it was simply breaking in those places not supported by cameral deposits *post mortem*. All arguments suggest that the presence of a cameral mantle is highly implausible. As a final nail in the coffin for the cameral mantle hypothesis, it would probably be difficult to maintain neutral buoyancy because of the added weight of the tissues, allowing only a smaller fraction of the chambers being filled by tissues and cameral deposits (compare Peterman *et al*. 2019).

The cameral fluid hypothesis solves some of the problems associated with the cameral mantle hypothesis (Fig. 10C). For example, there is no need for blood supply through the siphuncle. However, the precipitation from cameral liquid cannot explain the regular symmetric arrangement of these deposits. Previous authors suggested that the chamber has to be filled with a fluid phase in order to precipitate deposits on the entire inner surface, i.e., also on the dorsal side (Fischer & Teichert 1969; Blind 1991). In order to precipitate calcium carbonate in this configuration, the cameral liquid must be supersaturated with respect to CaCO_3_. This requires ion pumps to modify a large volume of cameral liquid, an energy-intensive process. More significantly, this model would prevent the conch from being buoyant, as the small juvenile chambers would never contain sufficient amounts of gas.

Conversely, adult chambers would need to be emptied without forming deposits. This mechanism would imply that there is a change in the siphuncle function during life and that juveniles were not buoyant at all. We think that this is unlikely because the structure of the siphuncle is consistent throughout ontogeny, and all modern cephalopods with internal and external shells can swim in the water column directly after hatching. Furthermore, if this model were correct, the deposits would be predicted to be oriented parallel to the gravitational surface (i.e., the growth axis in case of a horizontal orientation of the conch; but see Peterman *et al*. 2019). The deposits in *Trematoceras elegans* are tilted towards the outer shell wall both ventrally and dorsally, which is difficult to explain with the cameral liquid model.

Modifications of the cameral liquid model have been proposed, which may account for some of these inconsistencies. Fischer (in Fischer & Teichert 1969, p. 25) proposed that the inner shell surface “was not uniform in terms of its electrochemical configuration, in that it possessed areas receptive to polymerization of conchiolin which served as base for aragonite crystallization as well as areas not receptive to such overgrowth, i.e., that it was epitaxially differentiated”. The complexity and regularity of the cameral deposits make it difficult to conceive that the symmetry could be achieved consistently using the initial surface as the only control. It also does not explain the well-defined boundaries between the spherulitic sectors.

Another modification of the cameral liquid model was proposed by Crick (1982), which was apparently largely overlooked by later authors. He proposed that the chambers were not filled with cameral liquid, but the latter only covered the inner surface due to capillary forces. In Crick’s (1982) hypothesis, the siphuncle repeatedly secreted a carrier protein and mineralisation matrices to initiate the precipitation of aragonite. This carrier protein would also terminate the growth of the previous aragonite layer (Crick 1982). There are two main problems with this model: (1) it would require the transport of proteins through siphuncular epithelium, which has, to the author’s knowledge, not been documented in any living phragmocone-bearing cephalopod; and (2) it remains unclear how the proteins could be evenly distributed over the surface of the cameral deposits in order to maintain the symmetric growth.

For any modification of the cameral fluid hypothesis, the amount of fluid needed to precipitate a certain volume of cameral deposits presents a significant problem. For inorganic pore cements, the volume of pore fluid needed to provide enough CaCO_3_ to occlude a pore is not a fixed value. It depends on the pore size, the concentration of CaCO_3_ in the fluid, and the desired level of occlusion. Generally, the volume of fluid would need to contain sufficient CaCO_3_ to completely fill the pore volume and form a precipitate. As a rule of thumb, and taking average marine pore water as the fluid, approximately 1000 pore volumes of marine porewater must circulate through the open pore to provide sufficient CaCO_3_ to occlude it with cement (compare, e.g., Swart 2015; Immenhauser 2022). Consequently, even if fluid exchange between siphuncle and chamber was much higher in orthoceratoids than in *Nautilus* (e.g., Dzik 1984; Mutvei 2002), the rate at which chambers could be filled would be too low to be effective. An additional mechanism is therefore necessary to explain the high rate of fluid circulation.

### A new growth model

Because none of the above models can explain the observed features, we propose a new mechanism for the growth of cameral deposits (Fig. 10D). Our hypothesis is similar to the cameral fluid hypothesis but differs in two important aspects: (1) the involvement of cameral sheets; and (2) the precipitation from a thin film of cameral liquid only. The new hypothesis argues that cameral sheets laid down the basic structure of the cameral deposits during chamber formation. The cameral sheets were therefore secreted when the soft body detached from the last formed septum. The cameral sheets potentially represent a modification of the pellicle, an organic layer lining the inner surface of the chamber in *Nautilus* and ammonites (Denton & Gilpin-Brown 1966; Ward 1987). Because these sheets were primarily organic and hence subject to oxidation and microbial disintegration, they are difficult to observe in fossil specimens. That said, as laid out before, several arguments speak for their primary presence in the phragmocone of *Trematoceras* and other orthoceratoids.

The here-presented hypothesis suggests that the cameral sheets extended the functional surface of the pellicle and thus optimised bodily fluid transport within the chamber to the siphuncle. This led not only to higher liquid emptying rates but also to a greater amount of liquid being held back on the pellicle and surface of the sheets due to the capillary effect. In *Nautilus*, this effect enables the emptying of chambers despite being “decoupled” from the siphuncle (Ward 1979; Ward *et al*. 1980). The liquid trapped between the cameral sheets would then precipitate the deposits, possibly in combination with a chemical differentiation of the cameral surfaces as suggested by Fischer (in Fischer & Teichert 1969), controlling the nucleation sites on the shell wall and septum. The precipitation of the spherulites was initiated between the sheets and along the outer shell wall and blocked at the sheets. The ontogenetically oldest growth increments are accordingly located within the centre of the spherulites. Ions could be supplied through the siphuncle, as is also possible in *Nautilus* (Mangum & Towle 1982), probably at an increased rate. In comparison with the cameral liquid hypothesis, a much lower number of ions would be required to supersaturate the liquid due to its greatly reduced volume. There is no need for the transport of carrier proteins into the chambers, as suggested by Crick (1982).

The osmotic gradient that causes emptying of the chambers in *Nautilus* is mainly induced by Na^+^/Ka^+^-ATPase within the siphuncular epithelium (Greenwald *et al*. 1984). Accordingly, the concentration of Na^+^ is much higher within the siphuncle than within cameral liquid, and about 50 times higher than the concentration of Ca^2+^ in either of them (Mangum & Towle 1982). The solubility of NaCl (359 g/L) is significantly higher than the solubility of Ca(HCO_3_)_2_ (166 g/L). This implies that the cameral liquid can be supersaturated without necessarily causing a reversal of the osmotic gradient. That said, we do not exclude that periodical refilling of the chambers (e.g., due to changes in hydrostatic pressure during vertical migrations) occurred.

The epitaxial growth of the deposits was apparently restricted to areas where the liquid film was already in contact with previously formed carbonate but not on the cameral sheets. This could be explained by the contact of the liquid with the cameral deposits, producing a local solubility equilibrium. Thus, the number of transported ions was probably close to the threshold needed to induce precipitation. Notably, the proposed mechanisms also correspond to ion-by-ion, i.e., “classical” crystal mineralisation rather than a “non-classical” one, which appears to be otherwise typical for biomineral secretion (e.g., Cuif *et al*. 2010).

The bimineralic composition of the cameral deposits represents no obstacle to our hypothesis but, indeed, further supports it. Neither the cameral mantle nor the cameral fluid hypothesis would predict the co-precipitation of aragonite and calcite. Inorganic precipitation of aragonite and calcite is controlled by the concentration of Mg^2+^ within the bodily fluid (Eichinger *et al*. 2023 and references therein). Since our model involves very low volumes of liquid, concentrations may change rapidly during the precipitation of the cameral deposits. This is because ions are also continuously incorporated and, therefore, involve cycles of fluctuating ion concentrations, assuming that ions are continuously supplied through the siphuncle. It is further conceivable that the liquid volume was not constant, e.g., due to changes in osmotic pressure at different water depths. This process can explain vertical aragonite-calcite transitions.

In cephalopods, bimineralic shells have only been reported in some Palaeozoic orthoceratoids (De Baets & Munnecke 2018). Generally speaking, shell hardparts that include both aragonite and calcite, include, for example, the common blue mussel *Mytilus edulis*, among many other marine organisms (see discussion and critique Hahn *et al*. 2014). Lateral aragonite-calcite transitions are equally not uncommon in nature. With reference to the fabrics discussed here, this is likely because Mg^2+^ concentrations displayed gradients within the chamber fluid. In speleothems, to apply this non-biological reference, lateral aragonite-calcite transitions within growth increments occur if dripwater Mg^2+^ concentrations are close to the threshold between aragonite and calcite precipitation (Wassenburg *et al*. 2012, 2016). If ions are transported along the cameral sheets, fluid Mg^2+^ concentrations are expected to be higher within the fluid along the cameral sheets compared to the bulk fluid of the chamber, resulting in the precipitation of aragonite. Cathodoluminescence and SEM imaging confirm this notion, showing that aragonite is precipitated close to the cameral sheets (Fig. 2B, F, J, 8C, D; see also gaps between calcite spherulites in EBSD, Fig. 4A, E). The spatial distribution of calcite and aragonite is complex and likely related to the 3D-architecture of the cameral sheets and the liquid flow paths.

Although we did not investigate the geochemistry of the cameral deposits in detail, our growth model has implications for the interpretation of element concentrations and isotope signals from cameral deposits. The presumably rapid changes in ion concentrations would result in a complex pattern that is currently difficult to predict. Additionally, the interpretation of unaltered *versus* altered proxy data is challenging, as cameral deposits are most likely not the product of direct biological control and do not underlie the same processes as other typical biominerals. This implies that biological, kinetic and crystallographic parameters interact in a complex manner. Clearly, more work is needed to discriminate between primary and secondary geochemical data of cameral deposits, in particular, the geochemistry of the primary calcite phases is of interest.

In chronological order, our growth model for a single chamber is summarised as follows (see also Fig. 10D):

1. The mantle detaches from the last-formed septum and moves towards the aperture.
2. During this relocation of the mantle, organic cameral sheets are inserted at the apical end of the mantle, attached to the previous septum.
3. A new septum is mineralised, sealing off the chamber that is filled with cameral liquid. Note that further steps can occur in parallel with the formation of new chambers and additional cameral deposits, i.e., cameral deposits are precipitated simultaneously across multiple chambers.
4. The osmotic pumping activity of the siphuncle initiates to empty the chamber. The cameral sheets hereby increase the functional surface of the siphuncle and increase the rate of this process.
5. Due to capillary forces, a thin film of fluid remains on the cameral sheets and the inner surface of the chamber. Due to the decrease in volume, this fluid may already contain a higher concentration of Ca^2+^ ions from which precipitation of cameral deposits can begin.
6. Ca^2+^ and other ions needed for calcium carbonate precipitation are transported into the chamber via the siphuncle. Due to the relatively small volume of the fluid, supersaturation can be achieved quickly to induce precipitation of cameral deposits.
7. Growth increments are added while ions are further supplied through the siphuncle, either continuously or periodically. Precipitation continues until an obstacle is met, e.g., the siphuncle or cameral sheets.

The growth model presented here provides an internally consistent explanation of the bilaterally symmetric organisation of the cameral deposits, their regular ontogenetic progression, mineralogy and microstructure while allowing taxonomic variability in the shape of cameral deposits. Different primary arrangements of the cameral sheets arguably cause the latter. This could also explain the concentration of cameral deposits on the ventral side, although cyclic (partial) refilling of the chamber may have played a role as well, provided the shell was oriented in a horizontal position. The reason for the lack of cameral deposits in ammonoids despite the presence of cameral sheets is arguably related to differences in the transport activity of the siphuncle – if no ions are supplied to the cameral liquid, then no deposits are formed. Finally, our model does not exclude the possibility of forming cameral deposits when cameral sheets are absent – in this case, precipitation is predicted to occur at a much lower rate, resulting in strongly reduced cameral deposits. This prediction fits with the potential cameral deposit homologues in *Spirula* (Lemanis *et al*. 2020; Checa *et al*. 2022; Griesshaber *et al*. 2023) or *Nautilus* (Grégoire 1962, 1987; Mitchell & Phakey 1995). Coincidentally, the latter structures are formed where capillary forces are expected to be the highest, i.e., in the narrow interspace at the junction between the septum and the shell wall, leading to cameral liquid becoming ‘trapped’.

If our hypothesis holds, this provides further arguments to consider the Astroviida Zhuravleva & Doguzhaeva, 2004 [*emend*. King & Evans, 2019] and Pallioceratina Marek, 1998 [*emend*. King & Evans, 2019] as invalid (see discussion in King & Evans 2019). Both taxa are based on the presumed presence of a cameral mantle and not on directly observable diagnostic characters. Members of these groups need to be reinvestigated to ascertain their taxonomic position.

### Implications for functional morphology

Previous authors have exclusively attributed a counterbalancing function to cameral deposits (Teichert 1933, Flower 1955b; Fischer & Teichert 1969; Chamberlain 1993) or even considered them to be non-functional (Blind 1991). Only recent 3D models investigating buoyancy, orientation and stability have shown that this view may be too simplistic (Peterman *et al*. 2019). Additionally, *in-situ* observations of *Spirula spirula* in a hydrostatically unexpected apex-down position indicate that hydrostatic orientations may be more flexible than currently thought (Lindsay *et al*. 2020). The growth model presented here implies that cameral deposits had important functions beyond counterbalancing, thus providing a more reasonable trajectory for their evolutionary origin. If counterbalance was their only function, their appearance would be difficult to explain, as they would only be functional when fully formed while needing precise ontogenetic control as the soft body also increases continuously in weight. Under these conditions, a hypothetical evolutionary precursor stage of restricted, “non-functional” cameral deposits would probably not be fixed in a population, as they would present higher metabolic costs through added weight and a higher proportion of chambers to be emptied in order to maintain neutral buoyancy. In that sense, the virtual models of Peterman *et al*. (2019) suggest a vertical orientation with a stability-decreasing and, thus, mobility-increasing function of cameral deposits that can explain their evolutionary origin more elegantly. Still, the question remains why cameral deposits were concentrated on the ventral side of the conch. Thus, some taxa may have indeed had body chambers short enough to allow for a horizontal orientation of the conch. Alternatively, this asymmetry may have affected hydrostatic stability or was simply an artifact of morphogenesis.

Our proposed growth model implies further significant functional advantages of cameral deposits. First, due to the increased surface area of the cameral sheets, the functional area of the connecting ring and the pellicle is greatly increased. This enables improved bodily fluid transport rates, which are a limiting factor for the growth rates of phragmocone-bearing cephalopods (Ward 1982). This relationship results from the fact that the animals add weight during growth (i.e., shell and soft body), which needs to be compensated by emptying chambers. If growth proceeds faster than the chambers can be emptied, the animal becomes negatively buoyant and sinks. As an additional effect for increased growth rates, fully formed cameral deposits would completely seal the chambers, conveniently preventing passive liquid backflow into the chambers without expending pumping energy as in *Nautilus* (Greenwald *et al*. 1982).

Furthermore, the formation of cameral deposits effectively results in liquid ballast being replaced with solid ballast of higher density. Combined, this means that the animal can concentrate its pumping energy on a smaller total volume of liquid, thus further increasing chamber emptying rates and, by extension, growth rates. We, therefore, propose that the primary and probably plesiomorphic function of the combined cameral sheets and deposits was to increase growth rates. This also provides the possibility of a two-step evolutionary scenario in which cameral sheets evolved first, and cameral deposits came later, further increasing the adaptational value of the cameral sheets.

Another potential advantage of cameral deposits that has not been discussed before is that they reinforce the inner walls of the phragmocone, thus making them more mechanically resistant against crushing from, e.g., predator bite forces or hydrostatic pressure. Seuss *et al*. (2012a) documented a sublethally injured specimen, showing that the deposits were sufficient to withstand attacks, at least from smaller predators. The increased mechanical strength is also indicated by a tendency of chambers without deposits being crushed (e.g., Fig. 2G-H, S1F) and septa in these chambers often being destroyed (Mutvei 2018), although we did not statistically test this. This may be only a minor side effect, but testing different arrangements of cameral deposits by finite element analysis could provide clues as to whether their arrangement was also advantageous against crushing (compare, e.g., Lemanis *et al*. 2020; Lemanis & Zlotnikov 2023). As the apical portions of the shell are thinner and thus fundamentally weaker, cameral deposits, therefore, may have reinforced the mechanically weaker parts of the shell. Alternatively, the sealing of the apical chambers by the cameral deposits could have enabled orthoceratoids to lose their apical end without a major effect on their buoyancy and, hence, survival. This may have resulted in some extreme forms, where the process of decollating the apex was apparently induced deliberately (e.g., in *Sphooceras*, see Turek & Manda 2012).

## Conclusions

We provided a detailed description of cameral deposits of *Trematoceras elegans* from the Late Triassic St. Cassian Formation, leading to an updated terminology and the following observations:

- The cameral deposits are divided into regions of distal spherulitic sectors, plano-mural deposits and, in more advanced stages, spherulo-proximal deposits.
- The growth increments in the sectors were deposited simultaneously.
- The sectors are divided by radially arranged, originally organic cameral sheets.
- Cameral deposits display bilateral symmetry, although with common local irregularities.
- The original composition of the deposits is bimineralic, i.e., mostly primary aragonite in the plano-mural deposits and mostly primary calcite in the proximal and distal spherulitic deposits.
- The microstructure is fibrous in both the calcitic and aragonitic areas, with variable crystal size.

Based on these findings and detailed consideration of previous growth models, a new model for the formation of cameral deposits is proposed that is intrinsically stable when compared with the observations made. In our model, the cameral sheets play a crucial role, as they provide the means to distribute cameral liquid and ions effectively throughout the chamber while enabling control of the shape of the cameral deposits. If these arguments hold, then a 165-year-old palaeontological riddle has been solved. Moreover, the model allows for an improved understanding of orthoceratoid palaeobiology, as previous models implied widely different physiologies. We show that cameral deposits were functional beyond hydrostatics and allowed for higher growth rates. If true, then orthoceratoids also likely also had higher metabolic rates than *Nautilus*.

Cross-sections of cameral deposits are rarely reported, and we here call for an increased effort to study their taxonomic variability. Many taxonomic studies deem longitudinal sections sufficient, apparently assuming that cameral deposits would be more or less radially symmetric, apart from a decrease in thickness towards the dorsum. Our results and previous cross-sections show that this assumption is incorrect. Even better would be 3D reconstructions using CT-scanning or serial grinding tomography to avoid misinterpretations due to the plane of section (see Pohle *et al*. 2024; Turek & Aubrechtová 2024).

In addition, the geochemistry of cameral deposits needs further study to better understand their unique formation process from rapidly changing fluid compositions and how to differentiate between primary and diagenetically altered signals. Further investigations are also needed to confirm whether the bimineralic composition seen in *Trematoceras* was also prevalent in other orthoceratoids or whether some taxa built their deposits exclusively relying on aragonite or even calcite. If calcite is also prevalent in Palaeozoic orthoceratoids, cameral deposits may be used as a climate archive for proxy data, representing an analogue to belemnite rostra in the Mesozoic.

## Acknowledgements

We dedicate this paper to the late Harry Mutvei for his major contributions to cephalopod research, even if his interpretation of cameral deposits differed from ours. This study was funded by the Deutsche Forschungsgemeinschaft (DFG, German Research Foundation), project no. 507867999. AN acknowledges funding from the Deutsche Forschungsgemeinschaft (Project NU 96/11-1). MA was supported by the Research Plan of the Institute of Geology of the Czech Academy of Sciences (RVO67985831). Evelyn Kutatscher (Naturmuseum Bozen) and Andrzej Kaim (Institute of Palaeobiology, Warsaw) supported fieldwork in the St. Cassian Formation. Matthias Born (Ruhr University Bochum) is thanked for the preparation of thin sections. We are grateful to Sabine Weisel (Ruhr University Bochum) for polishing the samples prior to EBSD analysis. The constructive reviews by David Peterman (Penn State University) and two anonymous reviewers improved the final version of the manuscript.

## Author contributions

Conceptualisation: AP, RH, KS, AI; Data Curation: AP, RH; Formal Analysis: AP, RH; Funding Acquisition: AN, KS, AI; Investigation: AP, RH; Methodology: AP, RH, AI; Project Administration: AP; Resources: RH, AN; Supervision: AI; Validation: AP, RH, AN, BS, MA, BK, KS, AI; Visualisation: AP, RH; Writing – Original Draft Preparation: AP; Writing – Review & Editing: AP, RH, AN, BS, MA, BK, KS, AI.

## Supporting information

**Appendix S1.** Extended results and additional figures, cited in the article (Fig. S1-S24).

## Data archiving statement

Data for this study are available in the Dryad Digital Repository: XX

