## Supporting Information for "Microstructural and geochemical evidence offers a solution to the cephalopod cameral deposits riddle"

Alexander Pohle[1], René Hoffmann[1], Alexander Nützel[2], Barbara Seuss[3], Martina Aubrechtová[4], Björn Kröger[5], Kevin Stevens[1], and Adrian Immenhauser [1,6]

[1]Institute of Geology, Mineralogy and Geophysics, Ruhr University Bochum, Universitätsstraße 150, 44801 Bochum, Germany;

[2]SNSB-Bayerische Staatssammlung für Paläontologie und Geologie, Richard-Wagner-Straße 10, 80333 Munich, Germany;

[3]GeoZentrum Nordbayern, Friedrich-Alexander University Erlangen-Nürnberg, Loewenichstraße 28, 91054 Erlangen, Germany;

[4]Institute of Geology, Czech Academy of Sciences, Rozvojová 269, 16500 Prague 6, Czech Republic;

[5]Finnish Museum of Natural History, University of Helsinki, P.O. Box 44, Fi-00014 Helsinki, Finland;

[6]Fraunhofer Institution for Energy Infrastructures and Geothermal Systems, Am Hochschulcampus 1, 44801 Bochum, Germany

### Extended results

All references are contained in the main article.

**Absence of cameral deposits:** The largest *Trematoceras elegans* specimen PZO 16536, with a diameter of 7 mm, shows no traces of cameral deposits (Fig. S2). Another specimen with a diameter of 2 mm (PZO 16537) is largely pyritised (Fig. S3), and the position of the siphuncle could not be identified. Therefore, it cannot be excluded that this phragmocone actually belongs to an unidentified aulacoceratid or "mojsisovicsteuthid" coleoid (see Pohle & Klug 2024). Under CL microscopy, orange-luminescent, layered structures that are reminiscent of cameral deposits are visible on the mural part of this specimen but a diagenetic origin cannot be excluded (Fig. S3C-E).

All ammonite contain only pyrite and secondary, non-laminated cement (translucent under TL) within their chambers. The shell is well-preserved as indicated by cathodoluminescence (see below). Thus, there is no evidence for cameral deposits in these specimens.

**Additional description of structures:** The number of growth increments in the cameral deposits varies between specimens. In PZO 16538-02, which has a conch diameter of 2 mm, about 35 growth increments with a thickness of about 15 µm are present, whereas the 5-mm-wide PZO 16535-02 accumulated about 155 layers of approximately 10 µm thickness (Fig. S18). Growth increments alternate between light and dark brown layers, the latter possibly reflecting organic-rich layers (Fig. 2, S5-S11) with the organic inclusions resulting in a less translucent appearance. Individual growth increments can be traced across sector boundaries (Fig. 2, S6-S11).

Some areas of the deposits, especially within the planar deposits, are fibrous, with the elongation of the crystal fibres directed perpendicular to the growth increments (Fig. S10A, B, S11A). When studied under transmitted light, these growth increments appear to have more gradual boundaries compared to others with clear incremental boundaries.

**Cathodoluminescence:** The shell wall and septa of the ammonoids display bluish luminescence, which indicates that they are well-preserved (Fig. S4C, D). The same applies to the large specimen *Trematoceras elegans* PZO 16536 that is devoid of cameral deposits (Fig. S2C, D). Thus, it appears unlikely that cameral deposits are missing due to diagenesis. In contrast, the shell and septa of the pyritised orthocone (Fig. S3C-E) and the embryonic shell (Fig. S5C, D) display a yellowish-orange luminescence colour, a feature that might point to a secondary fabric and diagenetic alteration. Nevertheless, the fabric of the cameral deposits in the embryonic shell is still clearly visible under TL, pointing to a fabric-retentive alteration.

Features of local alteration of cameral deposits in otherwise well-preserved fabrics are common. For example, the primary fabric of the cameral deposits is locally obliterated in PZO 16535-02 and replaced by more irregularly spaced growth layers or blocky calcite, while the immediately adjacent septum appears to be unaltered (Fig. S7B, D, F).

Under cathodoluminescence, the growth increments are generally well visible even in areas where the fibrous structure of the fabrics seems blurred under transmitted light (Fig. 2, S6C, D, S7C, D, S8C, D, S9C, D, S10C, D, S11C, D). The latter areas also tend to exhibit blue luminescence. In PZO 16535-02, the plano-mural deposits display an orange luminescence except for the outermost layers and the area where the septum meets the outer shell wall, and only on the side where the deposits are thicker (i.e., the presumed ventral side; Fig. 7C, D, S8C, D).

**Electron backscatter diffraction:** The low indexing rate may be partly caused by the specimens not filling the entire observation window and is likely also related to the small size of aragonite crystals (0.5 µm in diameter; Fig. 4B-H).

The diagenetic cement (i.e., areas of the chambers devoid of cameral deposits) was identified as calcite showing large grains with no preferential orientation, consistent with secondary blocky calcite. Minor constituents of dolomite (probably an identification error), pyrite and quartz were also detected.

**Microstructures:** Under the SEM, the septa differ from the shell wall in having smaller nacre tablets (Fig. S16A), although in PZO 16538-02, they appear to be partly destroyed, probably during preparation (Fig. 7H, J).

The growth of the cameral deposits comes to a halt at the siphuncle (Fig. 6G). In the case of more advanced growth stages, the deposits appear to have continued to grow through the connecting ring (Fig. S16E). The microstructure of the deposits within the siphuncle is similar to the rest of the cameral deposits, with elongated aragonite fibres (Fig. S16F). In cross-section, the connecting ring is only preserved where the cameral deposits touch the siphuncle (Fig. 2A-F, 6G, J, S16E).

In PZO 16538-01, there is a vertical, biconcave structure ventrally adjacent to the siphuncle, apparently part of the mural delta (Fig. 6A, J). This structure reaches about one-third towards the shell wall and is sharply delimited towards the cameral deposits by a thin sheet (Fig. 6K). The deposits appear to grow through this sheet, though they change from the fibrous aragonite to the less porous calcite morphology (Fig. 6K). Within the biconcave area, growth zones are only in part visible (Fig. 6J). The dorsal end of the biconcave area immediately adjacent to the siphuncle is sharply delimited and differs in its microstructure from all other cameral deposits, with the appearance of sheet-like scales that are oriented concentrically (Fig. 6L). No comparable structure is visible in other specimens, and it is not clear how far this feature extends in the longitudinal direction.

**Energy dispersive spectroscopy:** As would be expected for (low Mg) calcitic and aragonitic mineralogies, EDS mapping reveals that Ca, C and O are the main elements in the longitudinal section PZO 16535-02 (Fig. 5G). Point-measurements in the calcite spherulites of the cross-section PZO 16540-01 (Fig. 5F, H) display the same elemental composition. Minor elements include Fe, Mg and Sr. The differential incorporation of these elements into aragonite and calcite, respectively, is controlled by the different ionic radii of these elements, which corroborates the mineralogies identified via EBSD.

EDS point measurements of the sheet-like structures close to the biconcave area (Fig. 6K) and between the spherulo-distal deposits (Fig. 7H) indicate elevated levels of Fe and Mg (Fig. S22B). The elemental compositions of these sheets are similar to the secondary blocky calcite within the chamber (Fig. S22C). The structure that we identify as likely pellicle (Fig. 7J) displays the same elemental composition (Fig. S22E). Neither of these structures appears to contain detectable amounts of S and P, indicative of preserved organic material. The dorsal end of the biconcave area adjacent to the siphuncle in PZO 16538-01 (Fig. 6L) is the only place where noticeable levels of S and Na, as well as higher levels of Mg (1.8 wt% compared to 0.4% in the calcitic cameral deposits) were detected (Fig. S21).

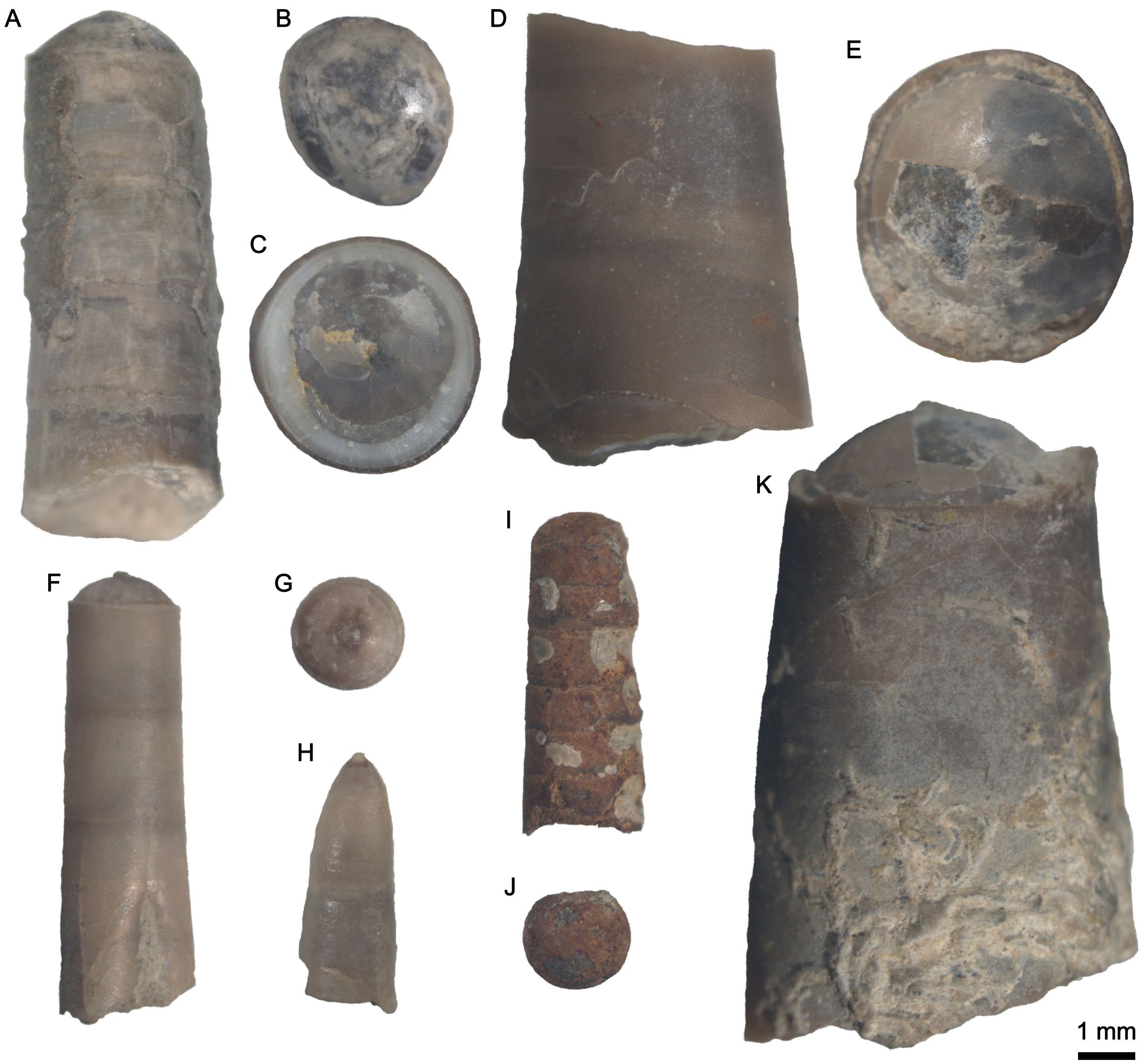


**Fig. S1.** *Trematoceras elegans* from the St. Cassian Formation, used in this study, prior to thin sectioning. A, PZO 16540, longitudinal view (orientation uncertain). B, PZO 16540, apical view, dorsal side up. C, PZO 16535, apical view, presumable dorsal side up. D, PZO 16535, longitudinal view (orientation uncertain). E, PZO 16536, apical view (orientation uncertain). F, PZO 16538, longitudinal view (orientation uncertain). G, PZO 16538, apical view, dorsal side up. H, PZO 16539, embryonic shell, longitudinal view (orientation uncertain). I, PZO 16537, longitudinal view (orientation uncertain). J, PZO 16537, apical view (orientation uncertain). K, PZO 16536, longitudinal view (orientation uncertain).

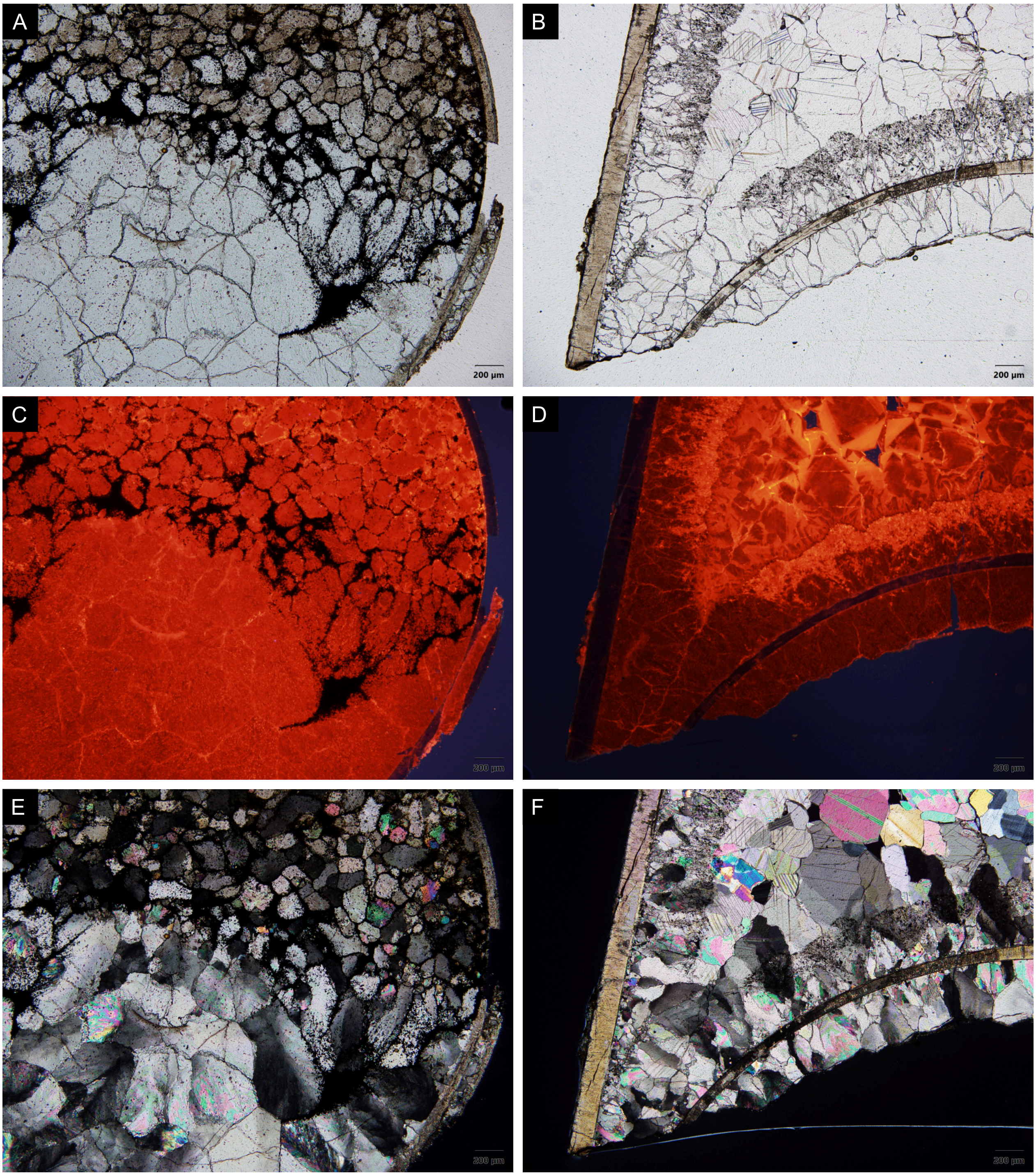


**Fig. S2.** Thin sections of *Trematoceras elegans*, PZO 16536, specimen without cameral deposits. A, PZO 16536-01, cross-section, transmitted light. B, PZO 16536-02, longitudinal section, transmitted light. C, PZO 16536-01, cross-section, cathodoluminescence. D, PZO 16536-02, longitudinal section, cathodoluminescence. E, PZO 16536-01, cross-section, polarised light. F, PZO 16536-02, longitudinal section, polarised light.

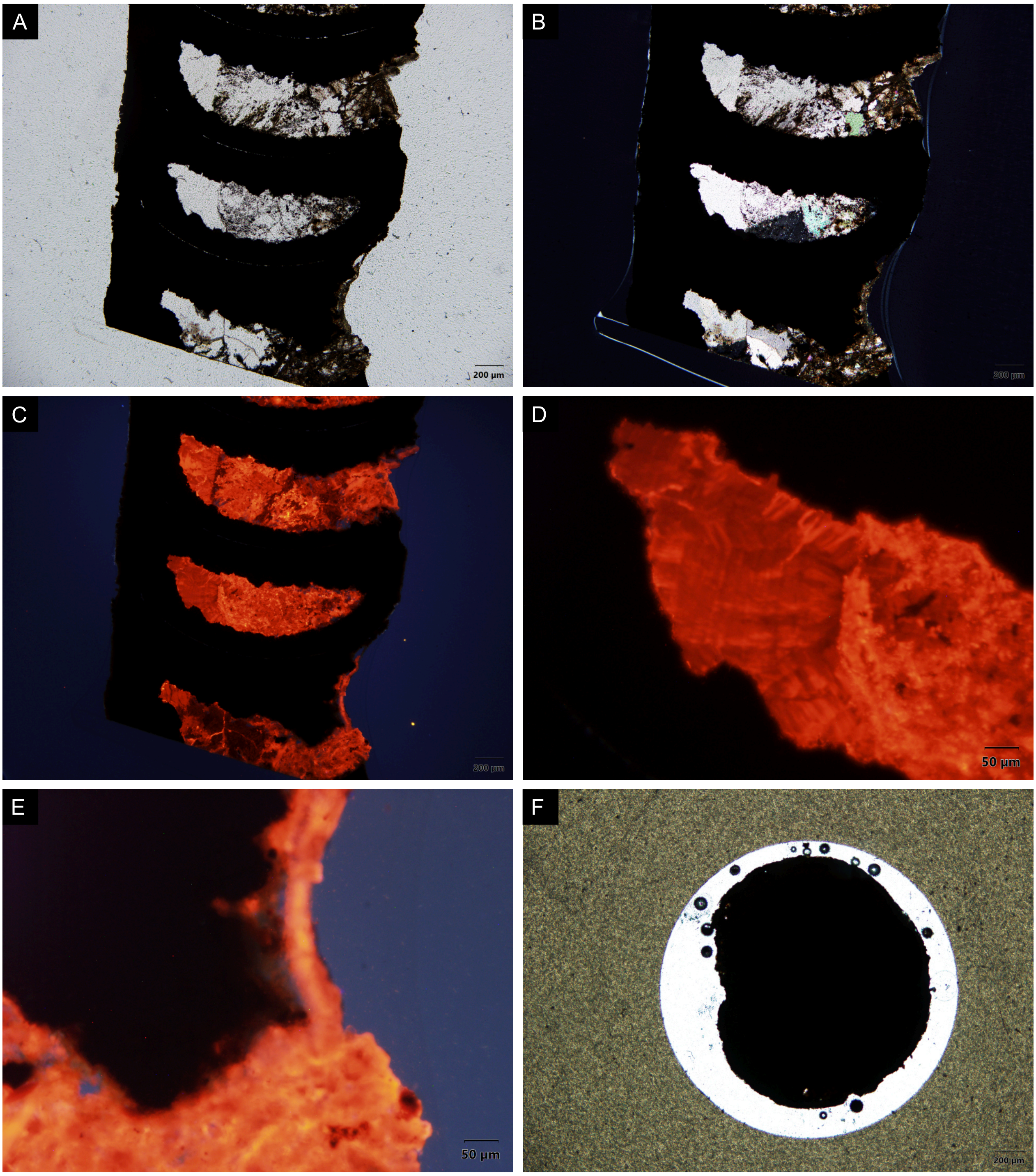


**Fig. S3.** Thin sections of *Trematoceras elegans*, PZO 16537, strongly weathered specimen probably without cameral deposits. May alternatively represent an aulacoceratid phragmocone, because siphuncle is not visible. A, PZO 16537-02, longitudinal section, transmitted light. B, PZO 16537-02, longitudinal section, polarised light. C, PZO 16537-02, longitudinal section, cathodoluminescence. D, detail of PZO 16537-02, longitudinal section, cathodoluminescence, potential cameral deposits. E, detail of PZO 16537-02, longitudinal section, cathodoluminescence, potential weathered remnant cameral deposits. F, PZO 16537-01, cross-section, transmitted light.

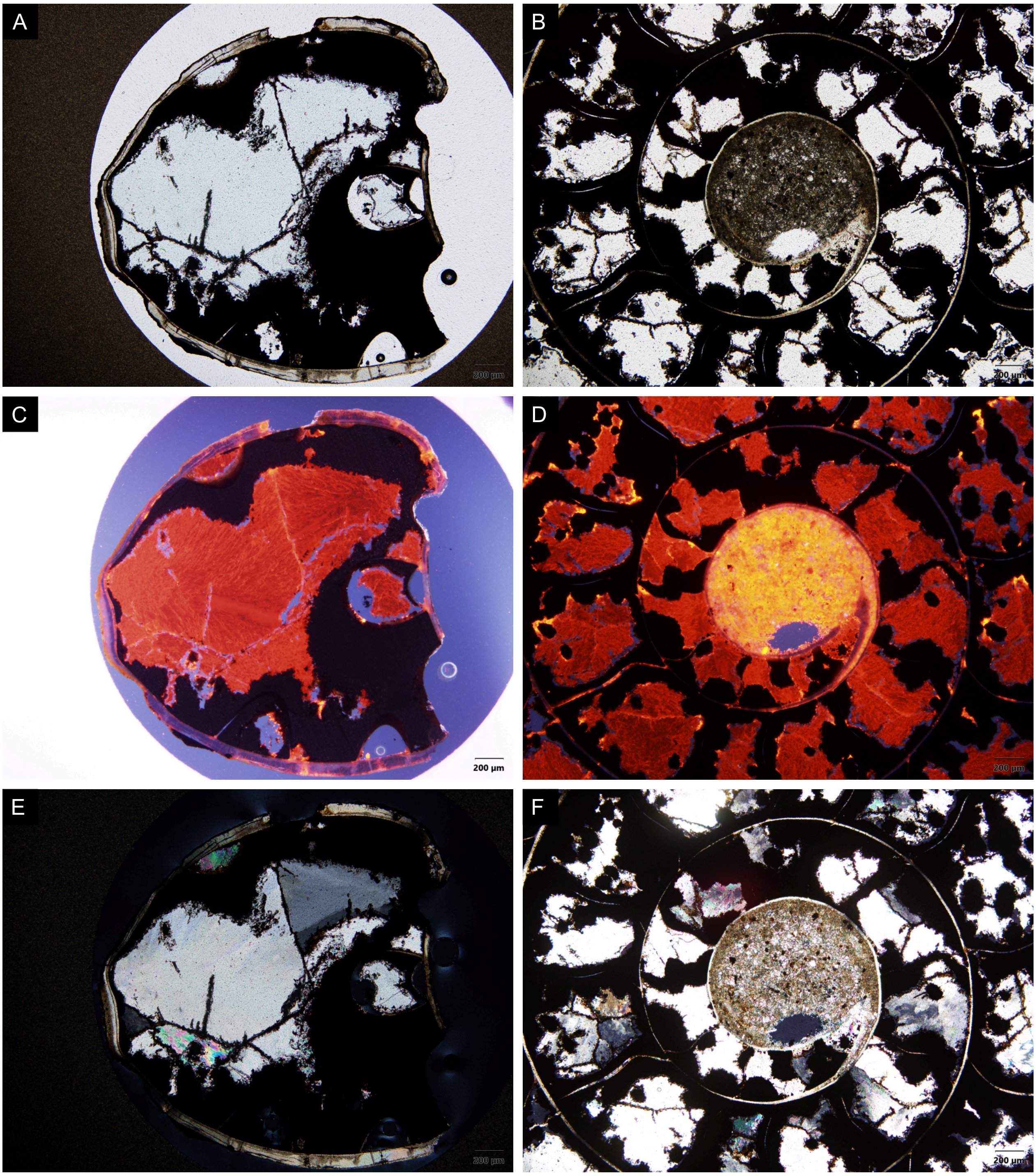


**Fig. S4.** Thin sections of co-occurring ammonoids. A, PZO 16557, cross-section, transmitted light. B, PZO 16558, longitudinal section, transmitted light. C, PZO 16557, cross-section, cathodoluminescence. D, PZO 16558, longitudinal section, cathodoluminescence. E, PZO 16557, cross-section, polarised light. F, PZO 16558, longitudinal section, polarised light.

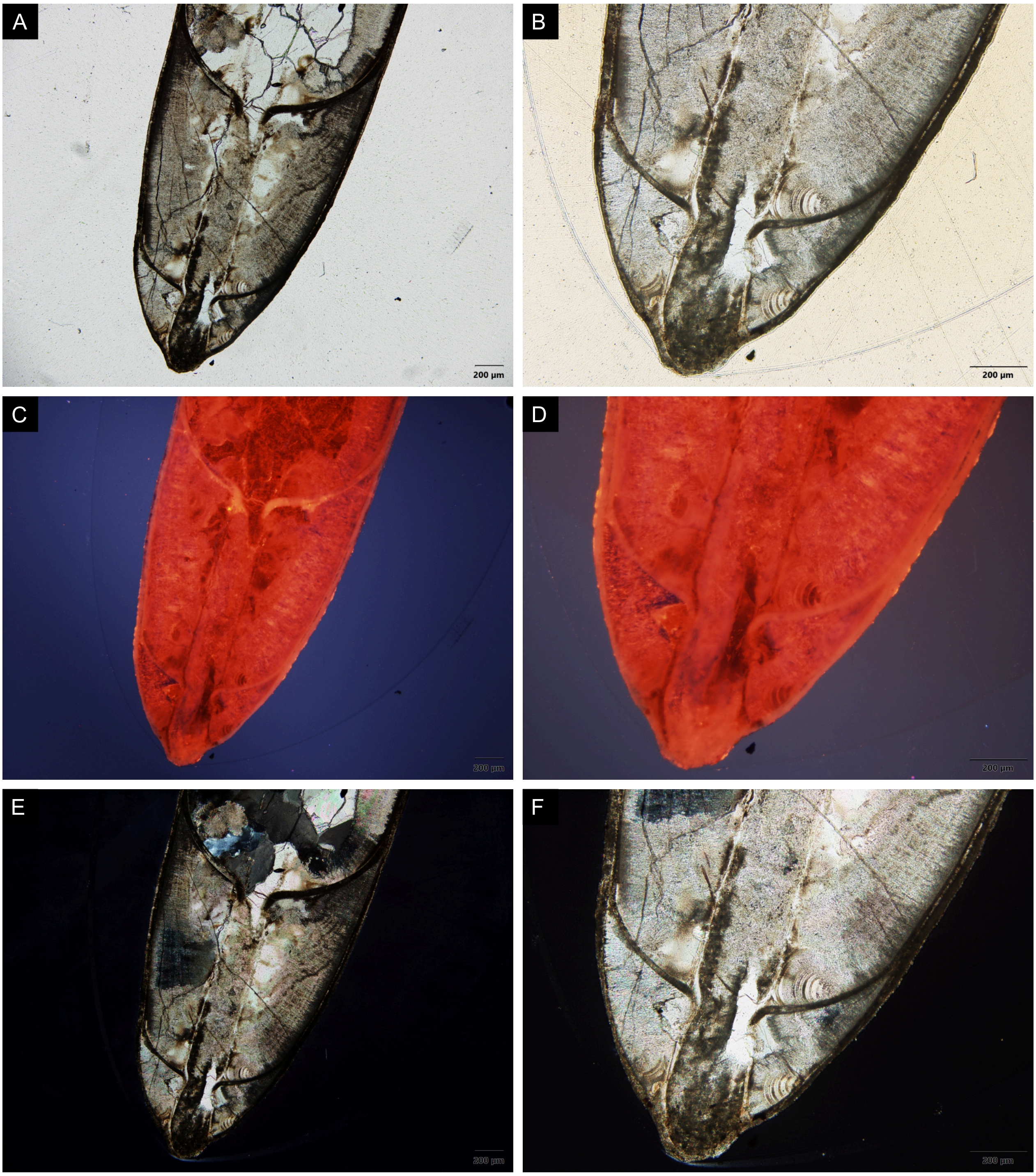


**Fig. S5.** Longitudinal thin section of the embryonic shell of *Trematoceras elegans*, PZO 16539. A-B, transmitted light. C-D, cathodoluminescence. E-F, polarised light.

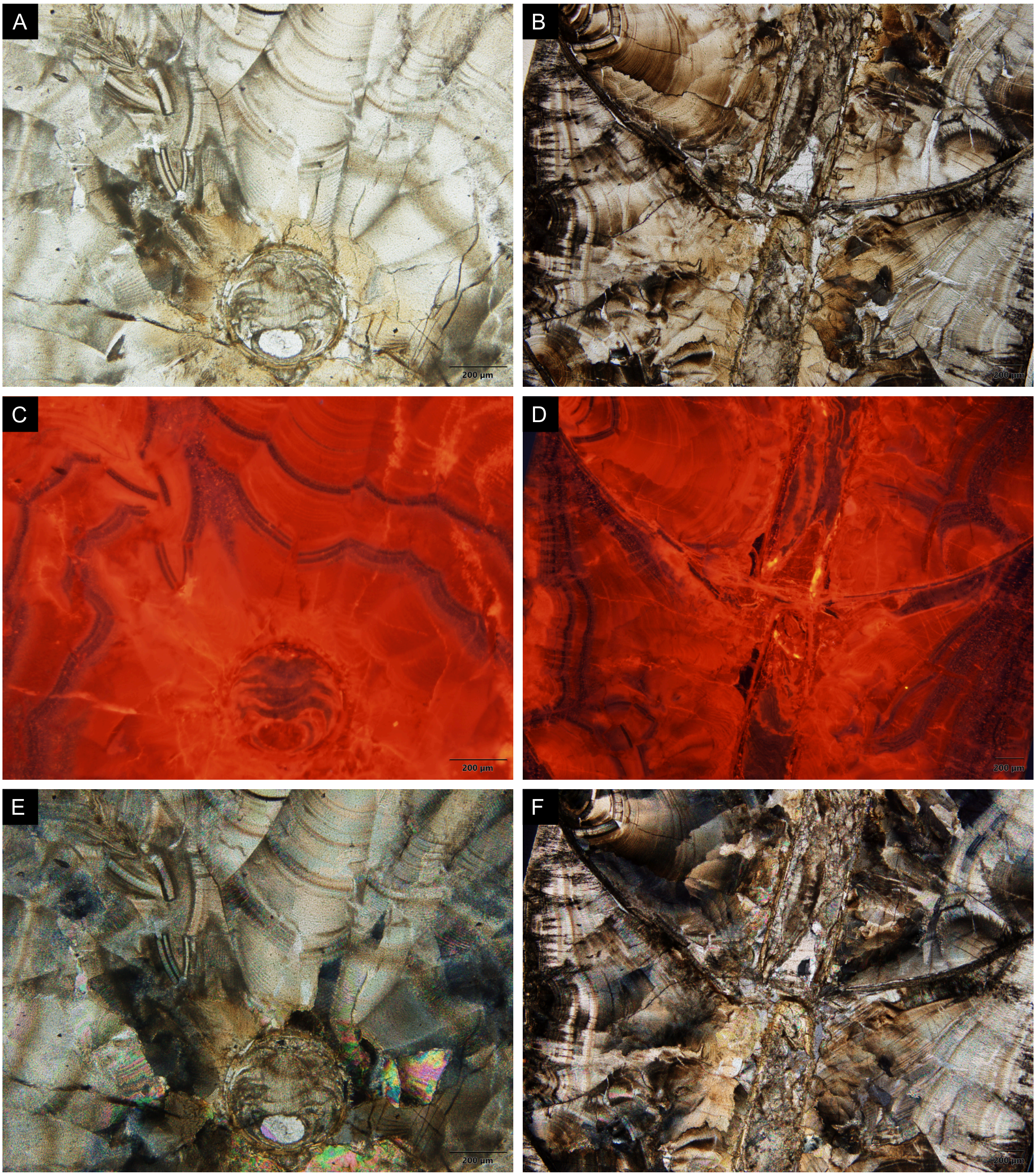


**Fig. S6.** Thin sections of *Trematoceras elegans*, PZO 16540, details of siphuncle. A, PZO 16540-01, cross-section, transmitted light. B, PZO 16540-02, longitudinal section, transmitted light. C, PZO 16540-01, cross-section, cathodoluminescence. D, PZO 16540-02, longitudinal section, cathodoluminescence. E, PZO 16540-01, cross-section, polarised light. F, PZO 16540-02, longitudinal section, polarised light.

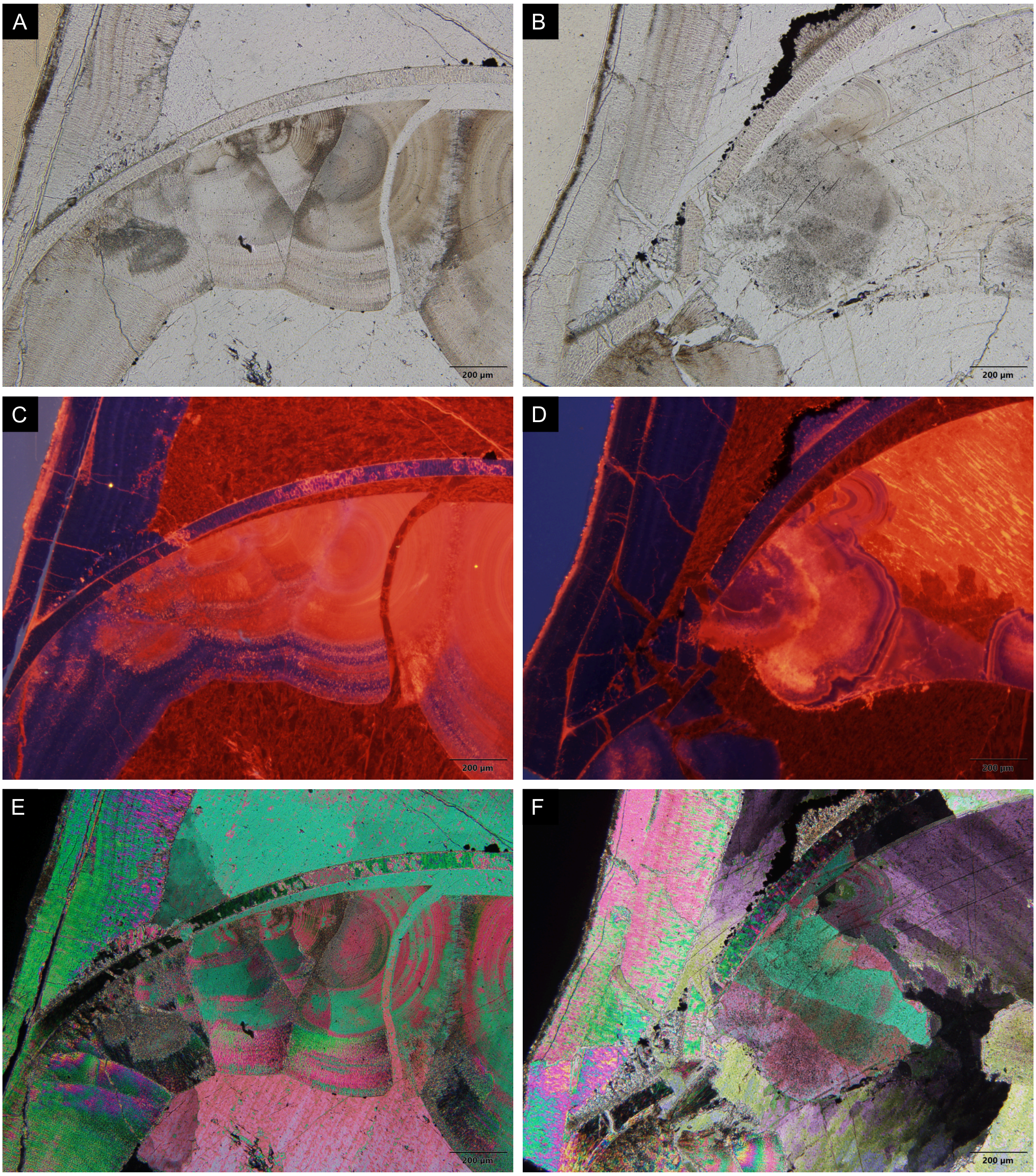


**Fig. S7.** Longitudinal thin section of *Trematoceras elegans*, PZO 16535-02, distal cameral deposits and diagenesis. A-B, transmitted light. C-D, cathodoluminescence. E-F, polarised light.

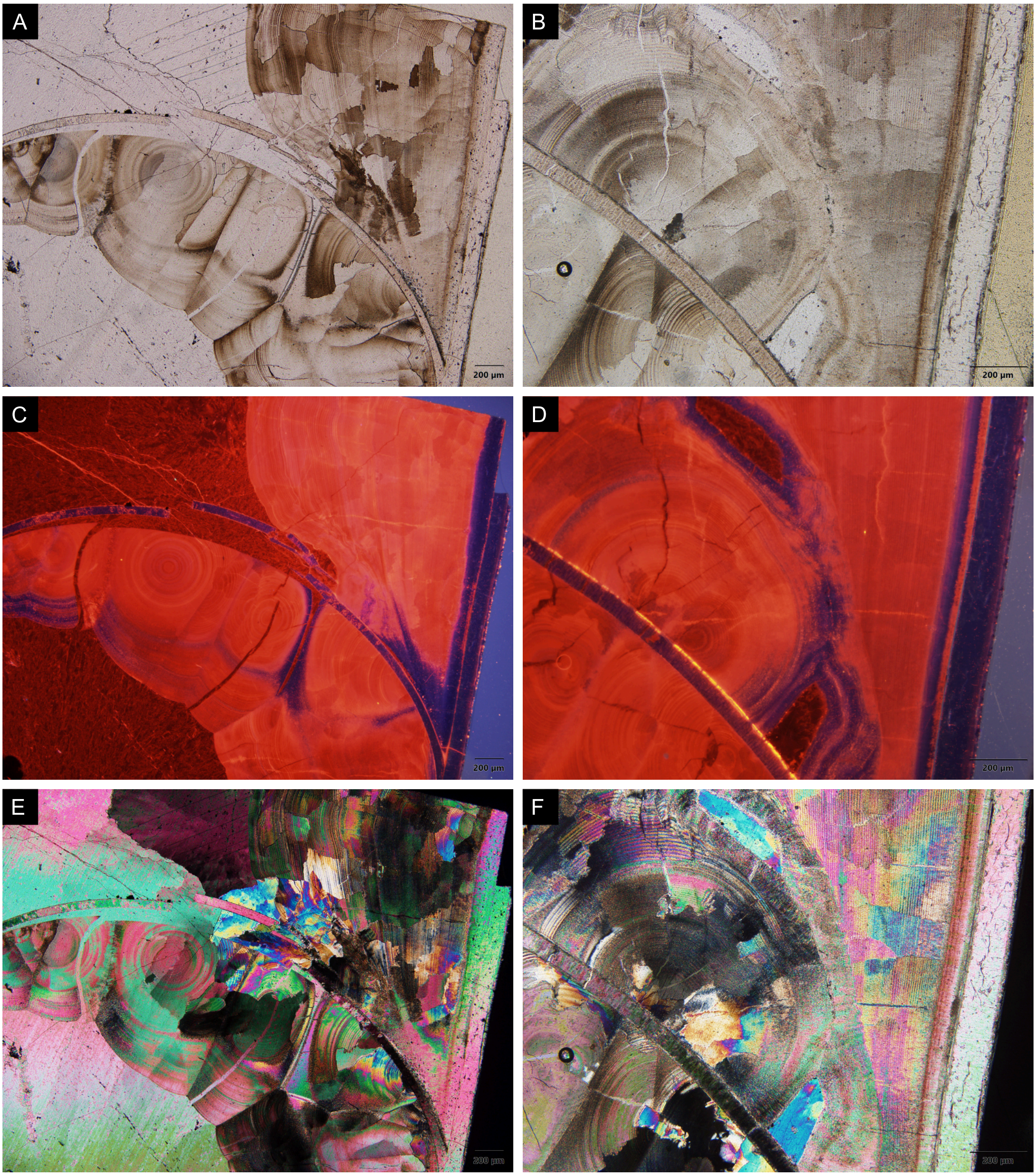


**Fig. S8.** Longitudinal thin section of *Trematoceras elegans*, PZO 16535-02, distal and proximal cameral deposits. A-B, transmitted light. C-D, cathodoluminescence. E-F, polarised light.

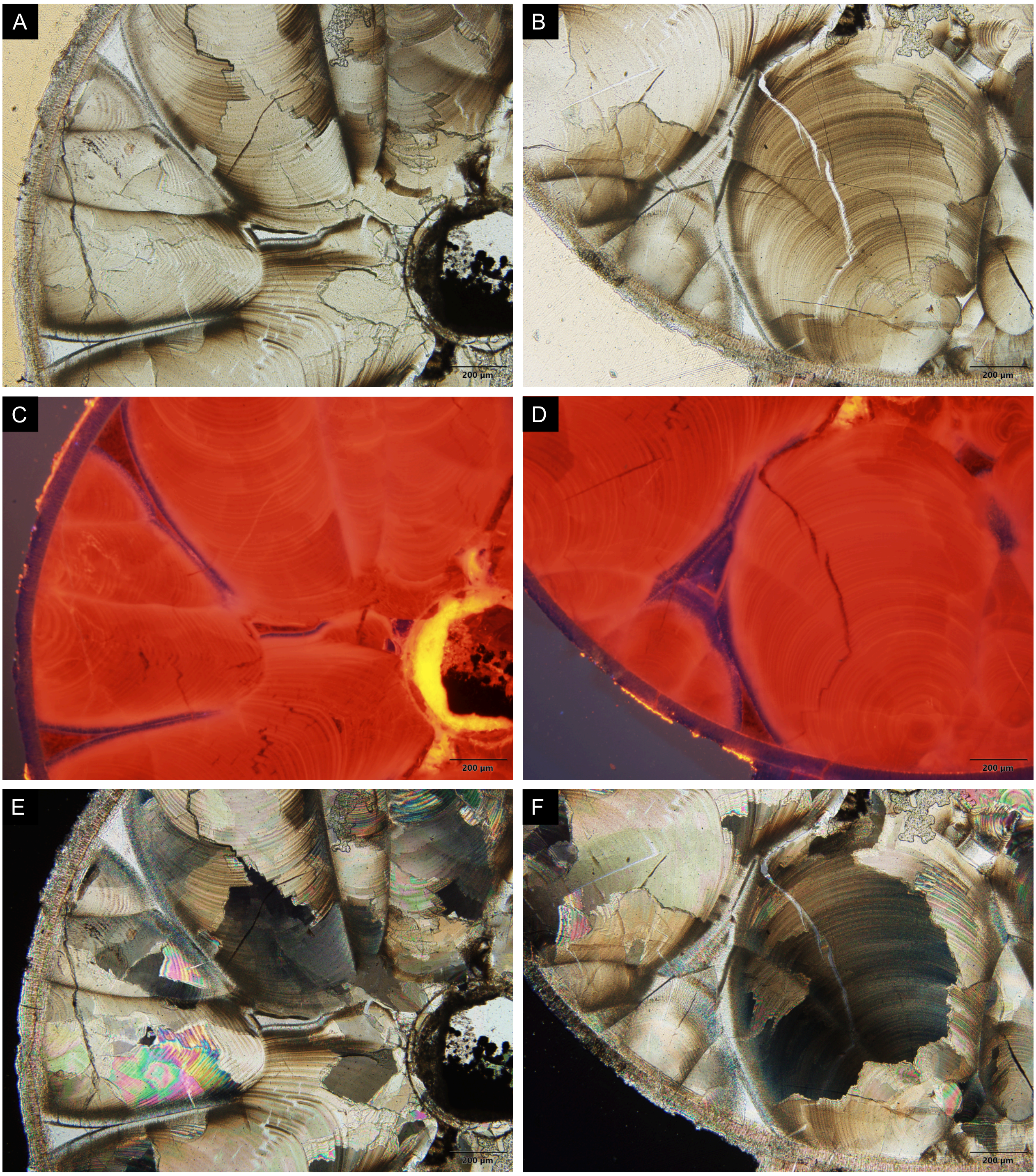


**Fig. S9.** Cross thin section of *Trematoceras elegans*, PZO 16535-01, close-up view of distal cameral deposits on the ventral side. A-B, transmitted light. C-D, cathodoluminescence. E-F, polarised light.

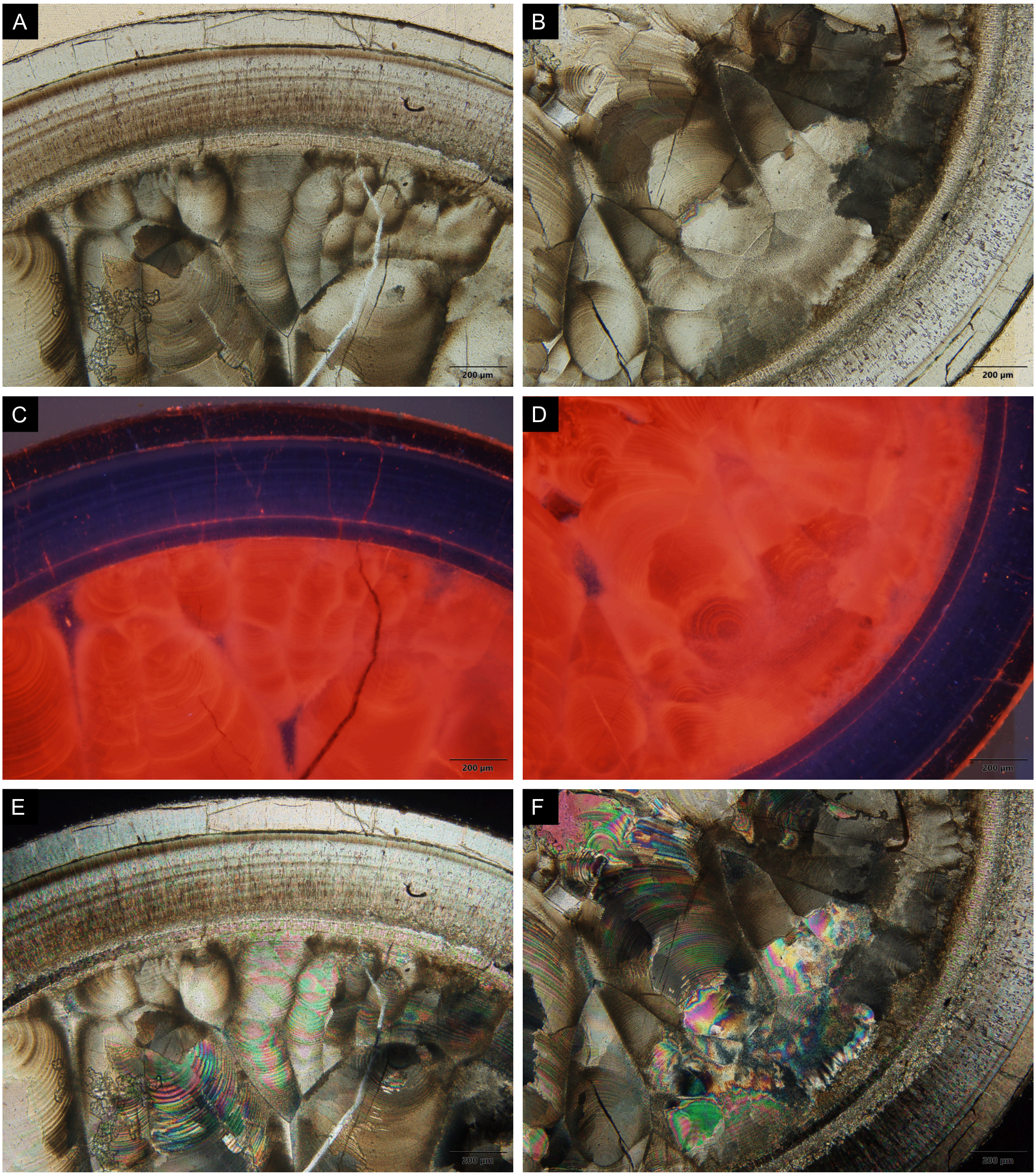


**Fig. S10.** Cross-section of *Trematoceras elegans*, PZO 16535-01, close-up view of lateral and dorsal cameral deposits. Note the septum dividing the cameral deposits in proximal (outer ring) and distal (inner) cameral deposits. A-B, transmitted light. C-D, cathodoluminescence. E-F, polarised light.

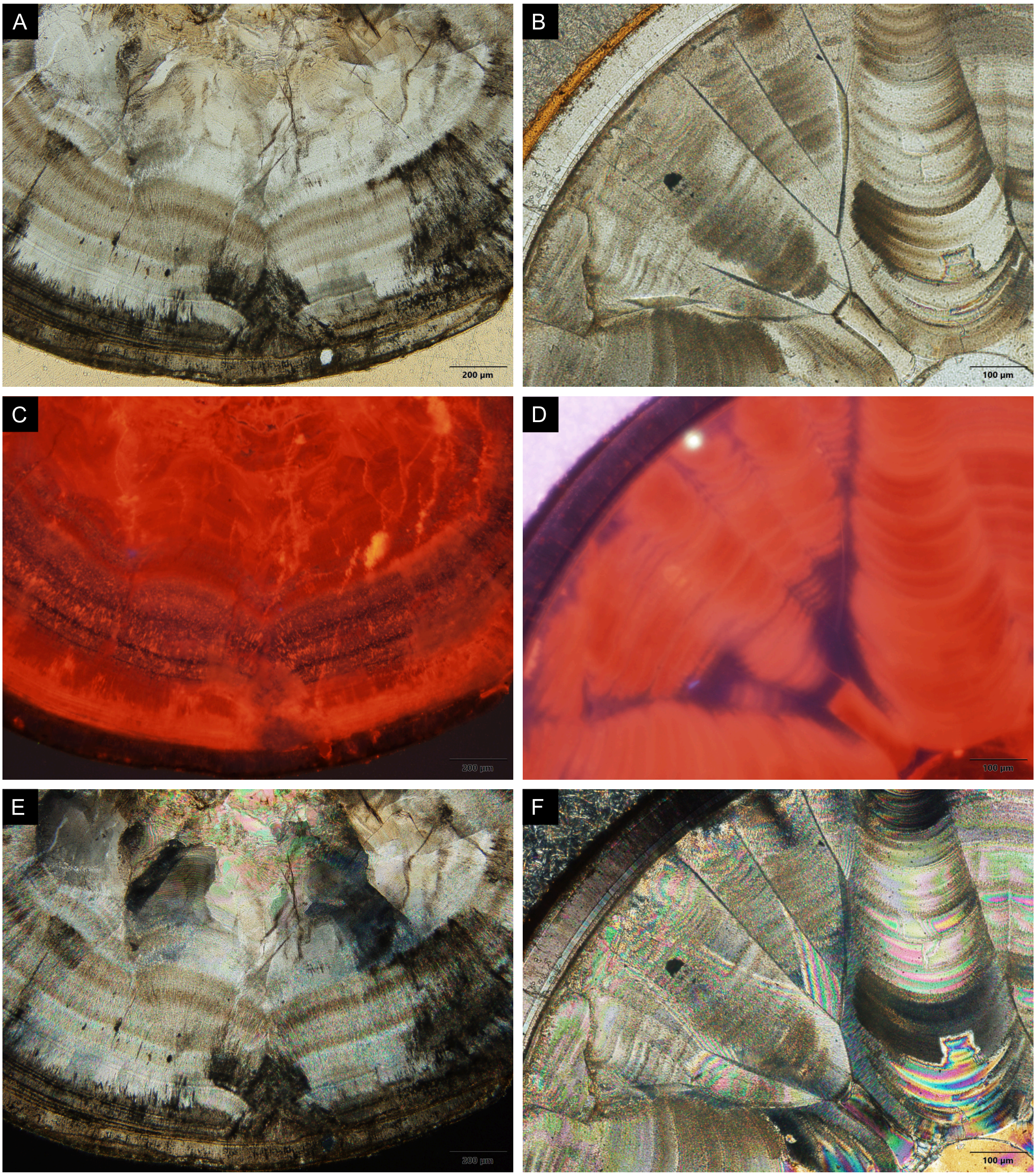


**Fig. S11.** Cross thin sections of *Trematoceras elegans*, details cameral deposit structures. A, C, E, PZO 16540-01, note potential dorsal cleft. B, D, F, PZO 16538-01, note mural delta. A-B, transmitted light. C-D, cathodoluminescence. E-F, polarised light.

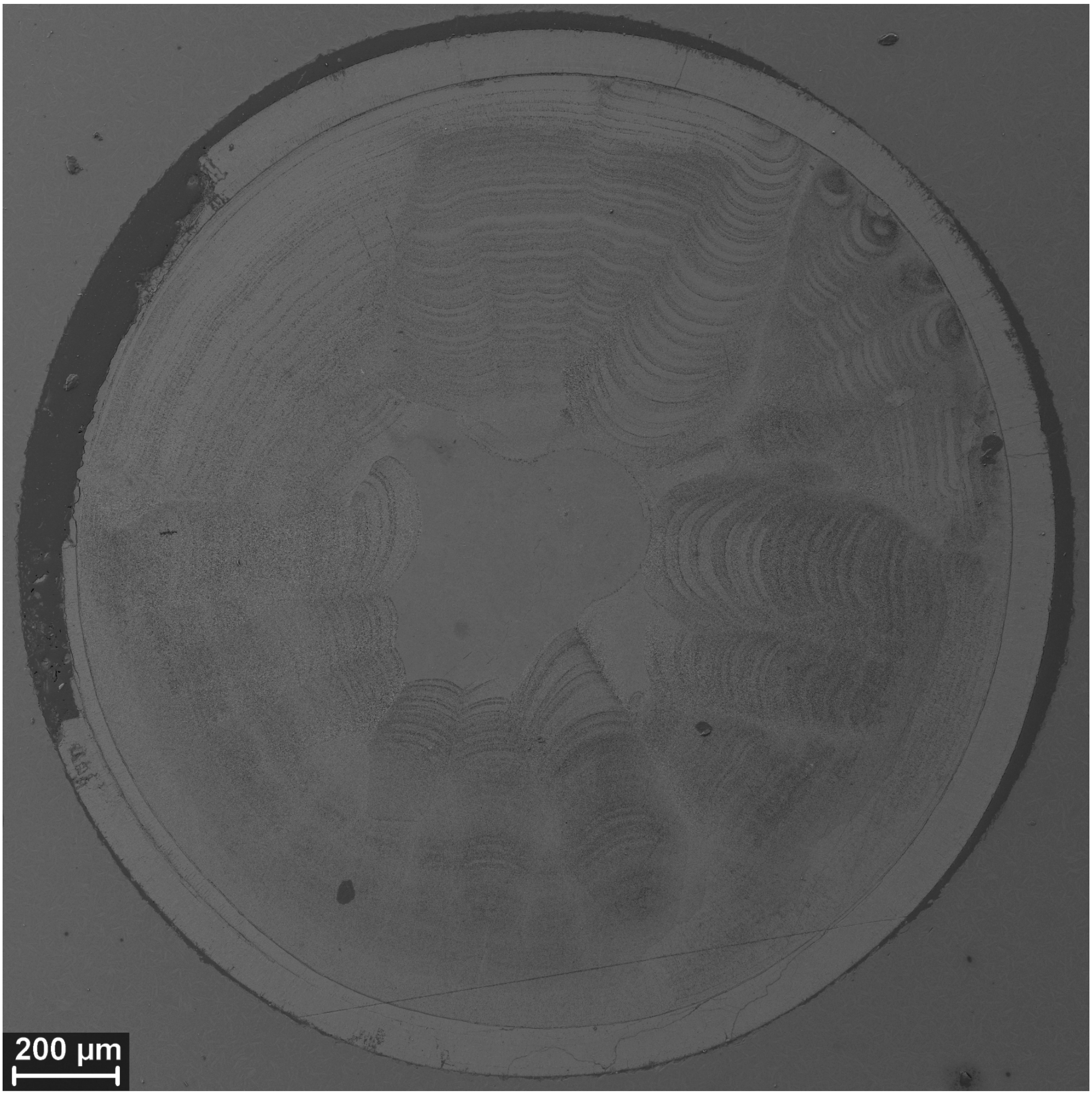


**Fig. S12.** SEM image of *Trematoceras elegans*, PZO 16538-01, cross-section. Specimen not oriented according to presumable symmetry plane, presumable ventral side approximately to the upper right (note mural delta).

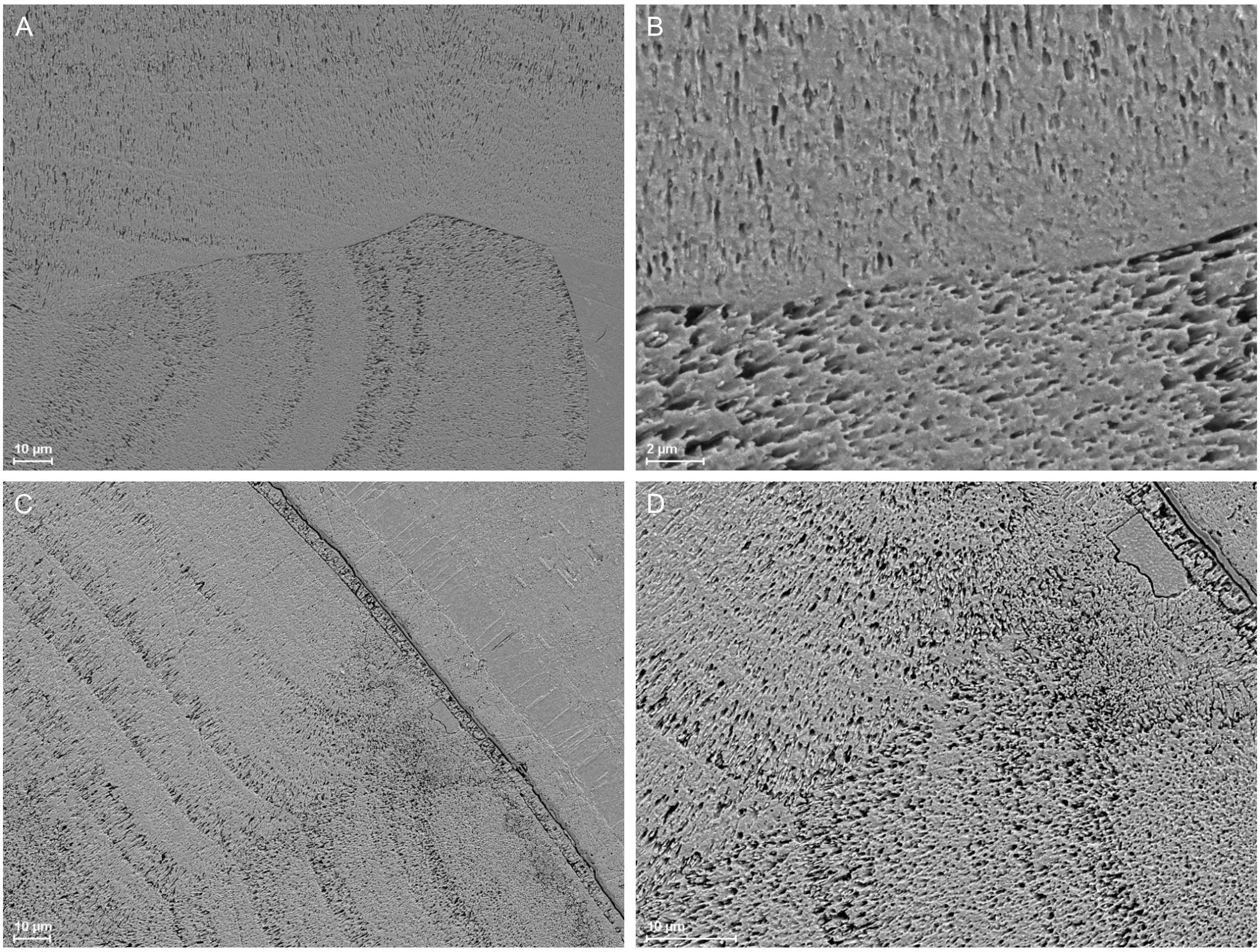


**Fig. S13.** SEM images of *Trematoceras elegans*, PZO 16538-01, cross-section, cameral deposits and cameral sheets. A-B, boundary between adjacent, differently oriented sectors. C-D, boundary between adjacent, similarly oriented sectors close to shell wall.

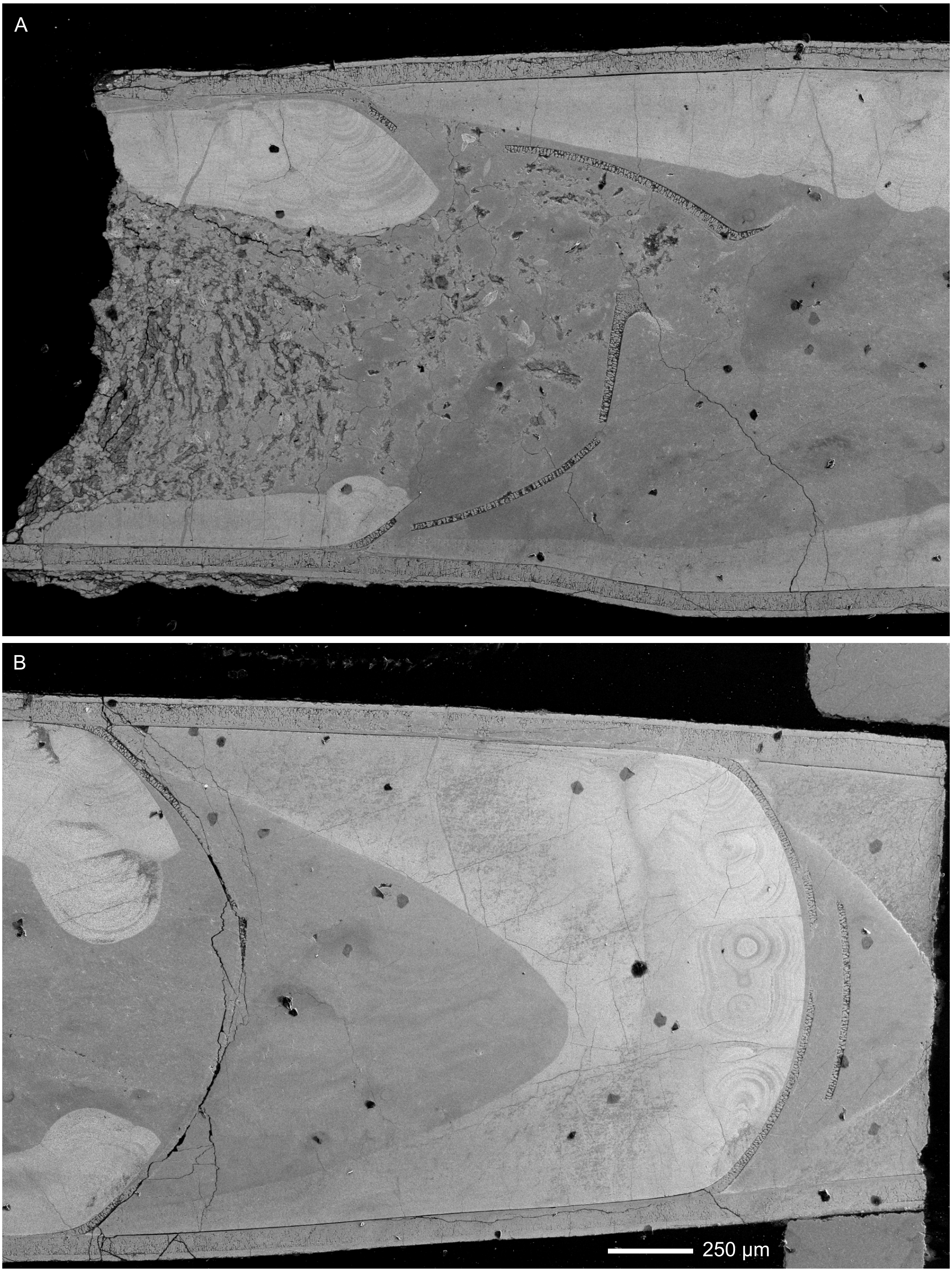


**Fig. S14.** SEM image of *Trematoceras elegans*, PZO 16538-02, longitudinal section. Dorsal side down. A, apertural part. B, apical half.

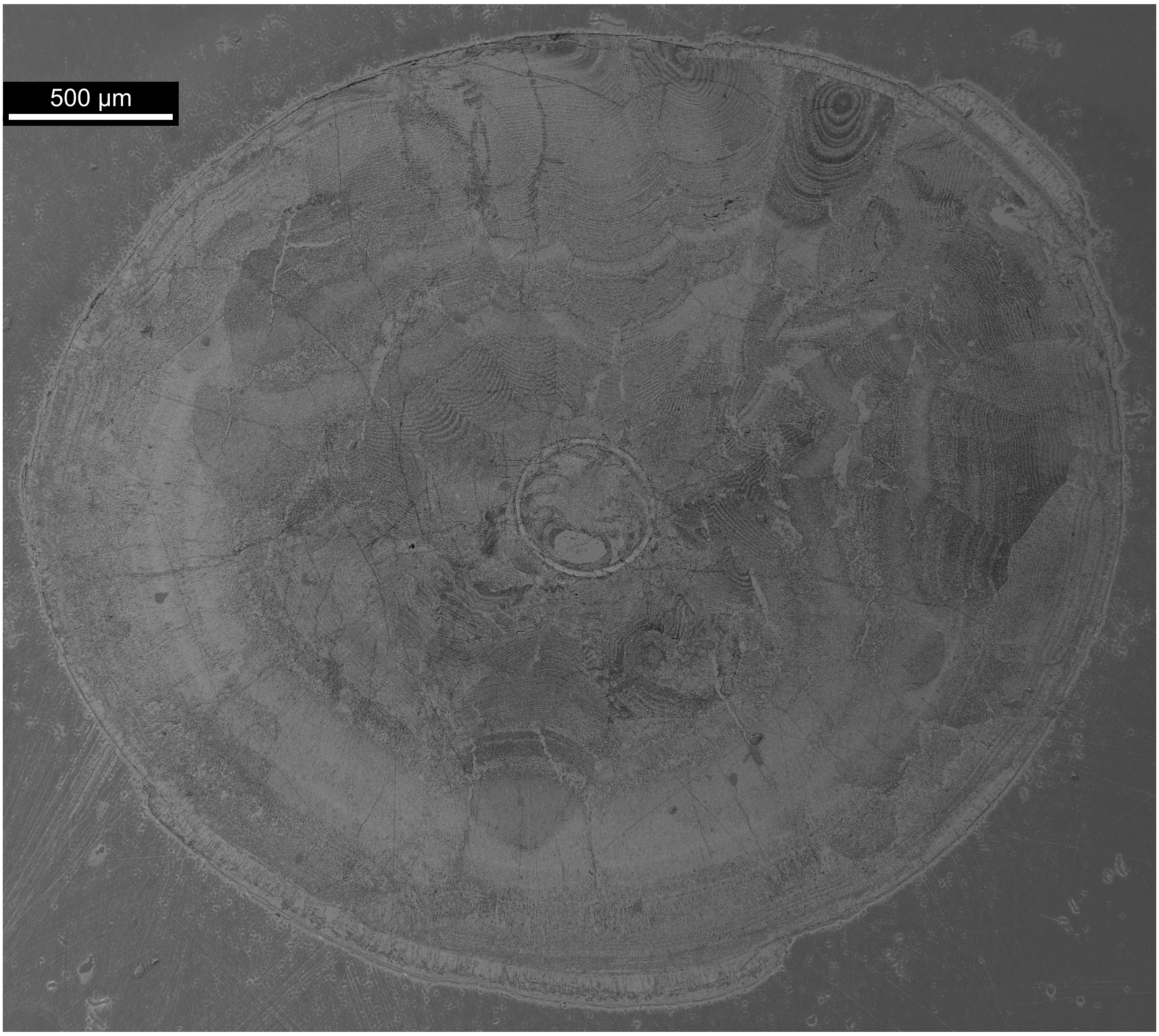


**Fig. S15.** SEM image of *Trematoceras elegans*, PZO 16540-01, cross-section. Specimen not oriented according to presumable symmetry plane, presumable dorsal side down.

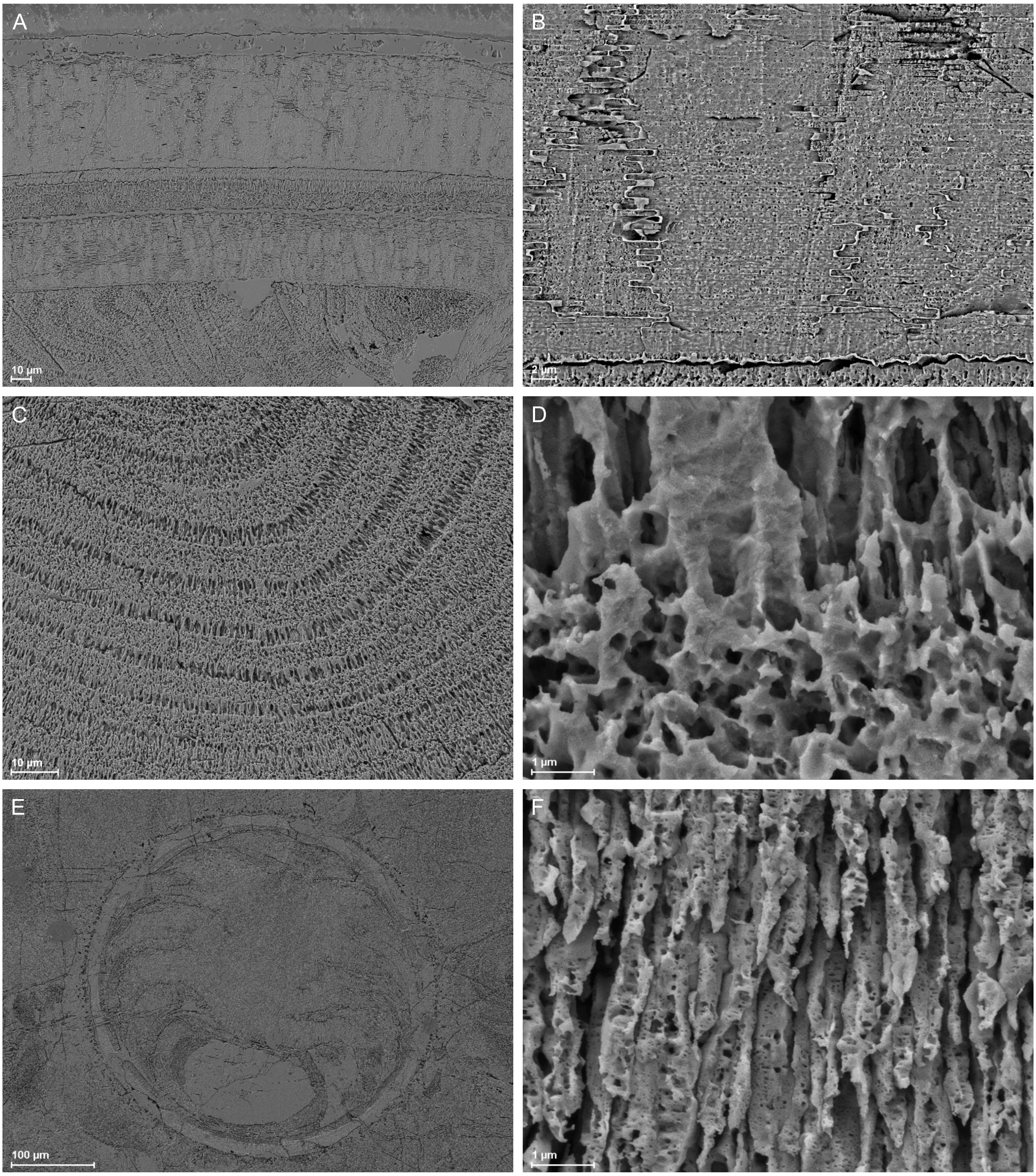


**Fig. S16.** SEM images of *Trematoceras elegans*, PZO 16540-01, cross-section. A, top to bottom: outer shell wall, proximal cameral deposits, septum, distal cameral deposits. B, nacre tablets of outer shell wall. C, single cameral deposit growth sector. D, microstructure of cameral deposits at the transition between growth increments. E, siphuncle with connecting ring and endosiphuncular deposits. F, microstructure of siphuncular deposits, consisting of aragonite needles

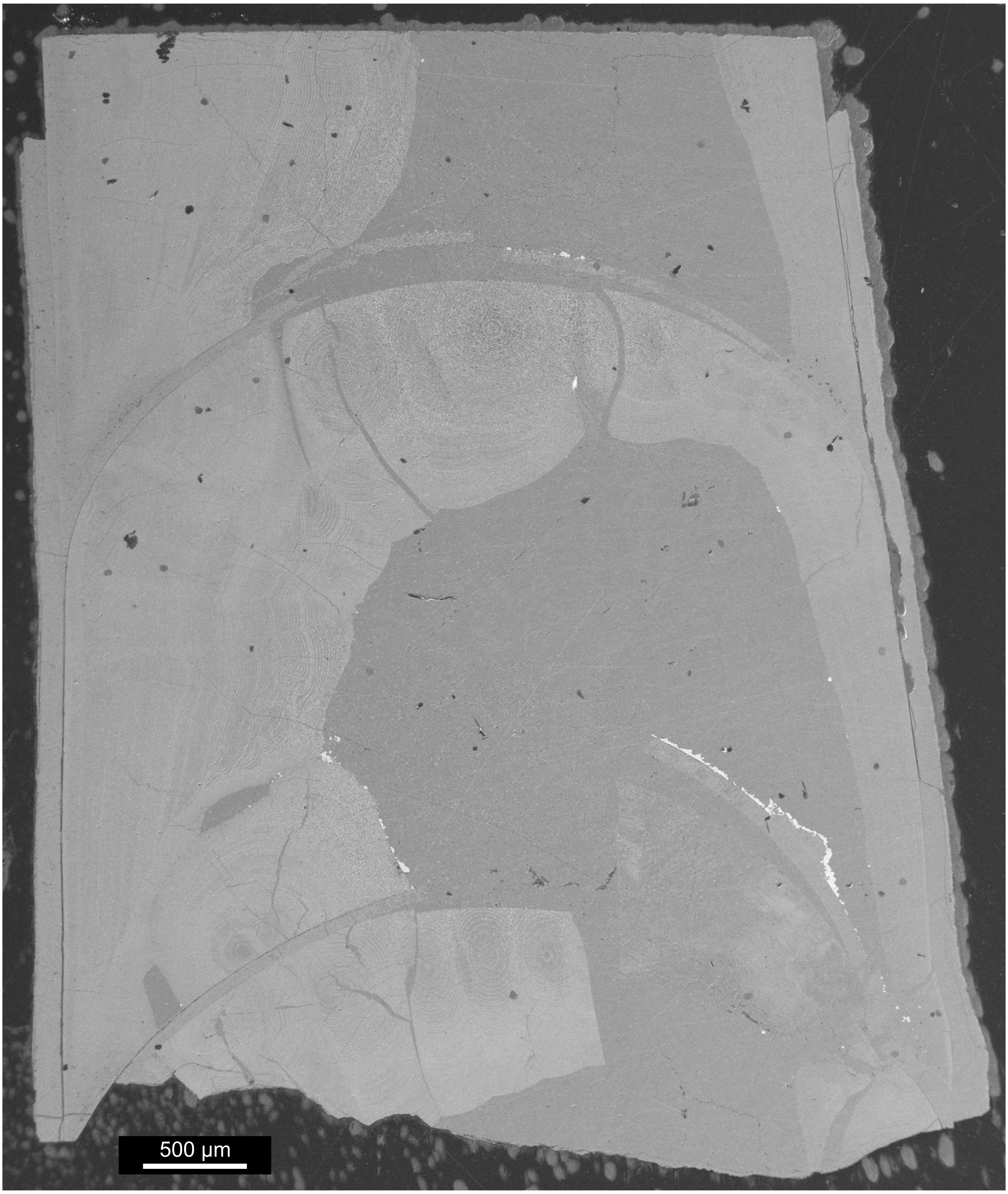


**Fig. S17.** SEM image of *Trematoceras elegans*, PZO 16535-01, cross-section. Presumable dorsal side to the right.

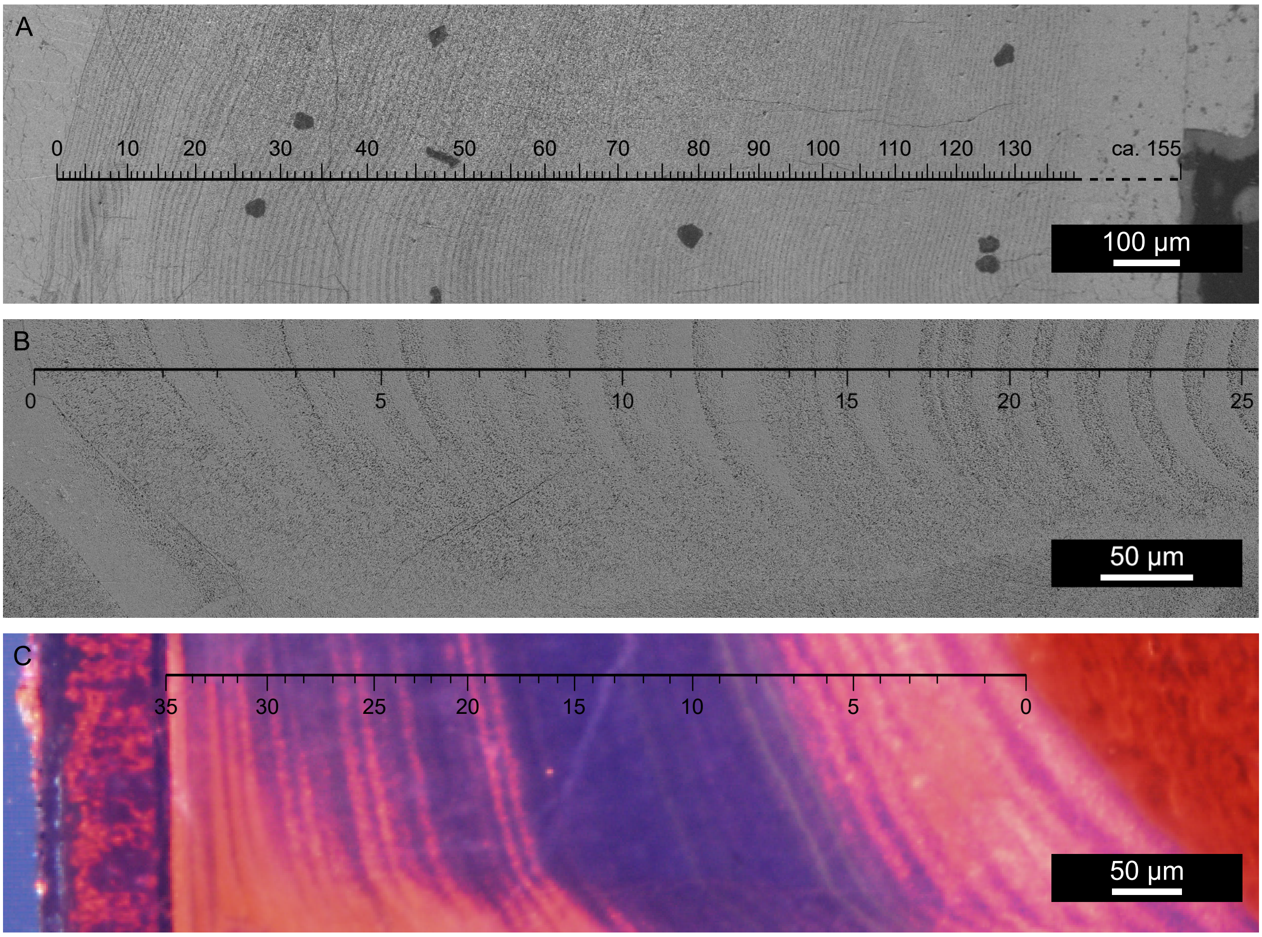


**Fig. S18.** Number of growth increments in cameral deposits of *Trematoceras elegans*. A, SEM image of longitudinal section, PZO 16535-02, plano-mural deposits. Shell wall right, layers closest to shell wall not discernable, thus estimated. B, SEM image of cross-section, PZO 16538-01, spherulo-distal deposits. Ventral shell wall right, siphuncle left. C, CL image of longitudinal section, PZO 16538-02, plano-mural deposits.

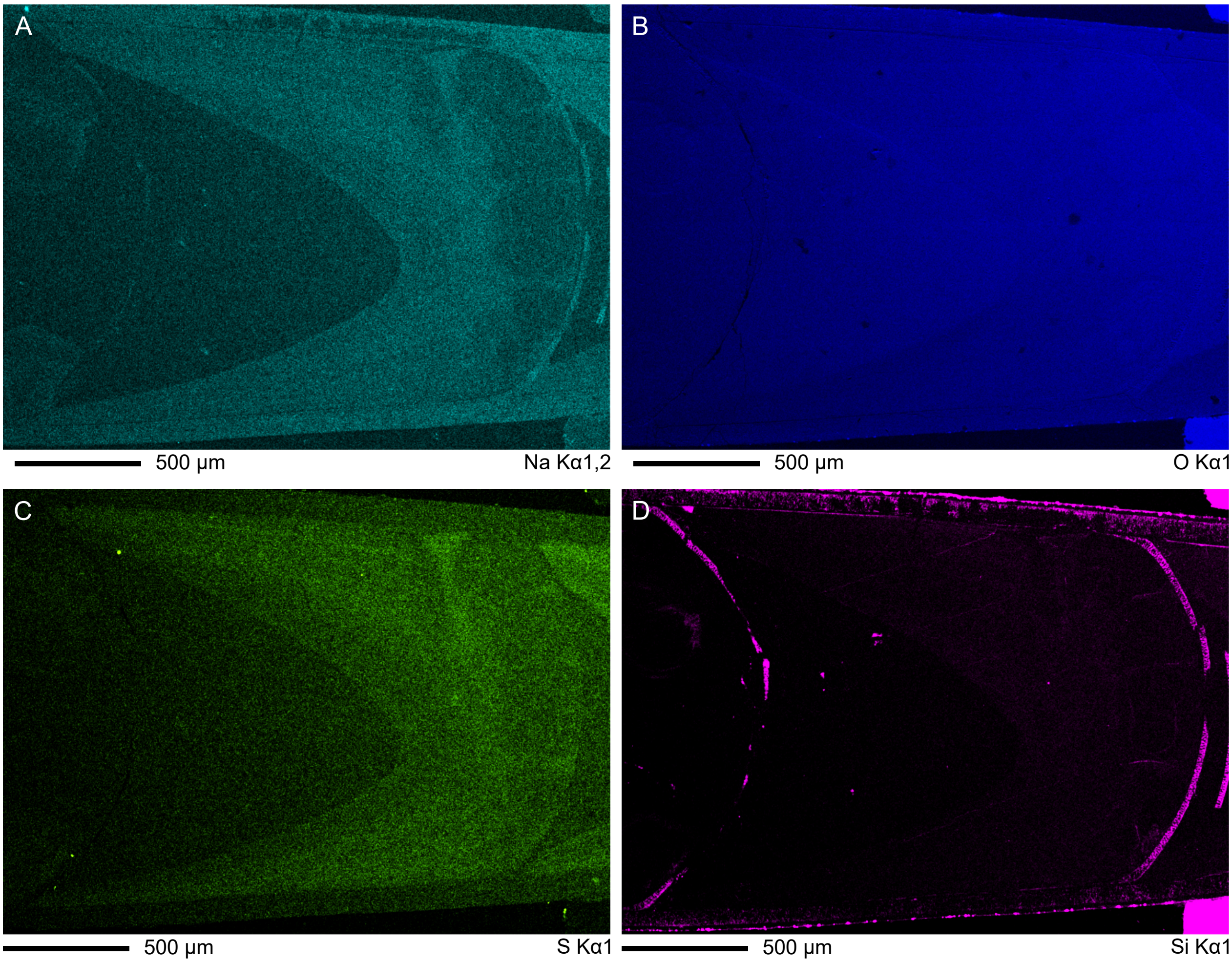


**Fig. S19.** Additional EDS element mappings of *Trematoceras elegans*, PZO 16538-02, longitudinal section. For further mappings of the same specimen, see Figure 5 in the main article. A, sodium. B, oxygen. C, sulphur. D, silicon.

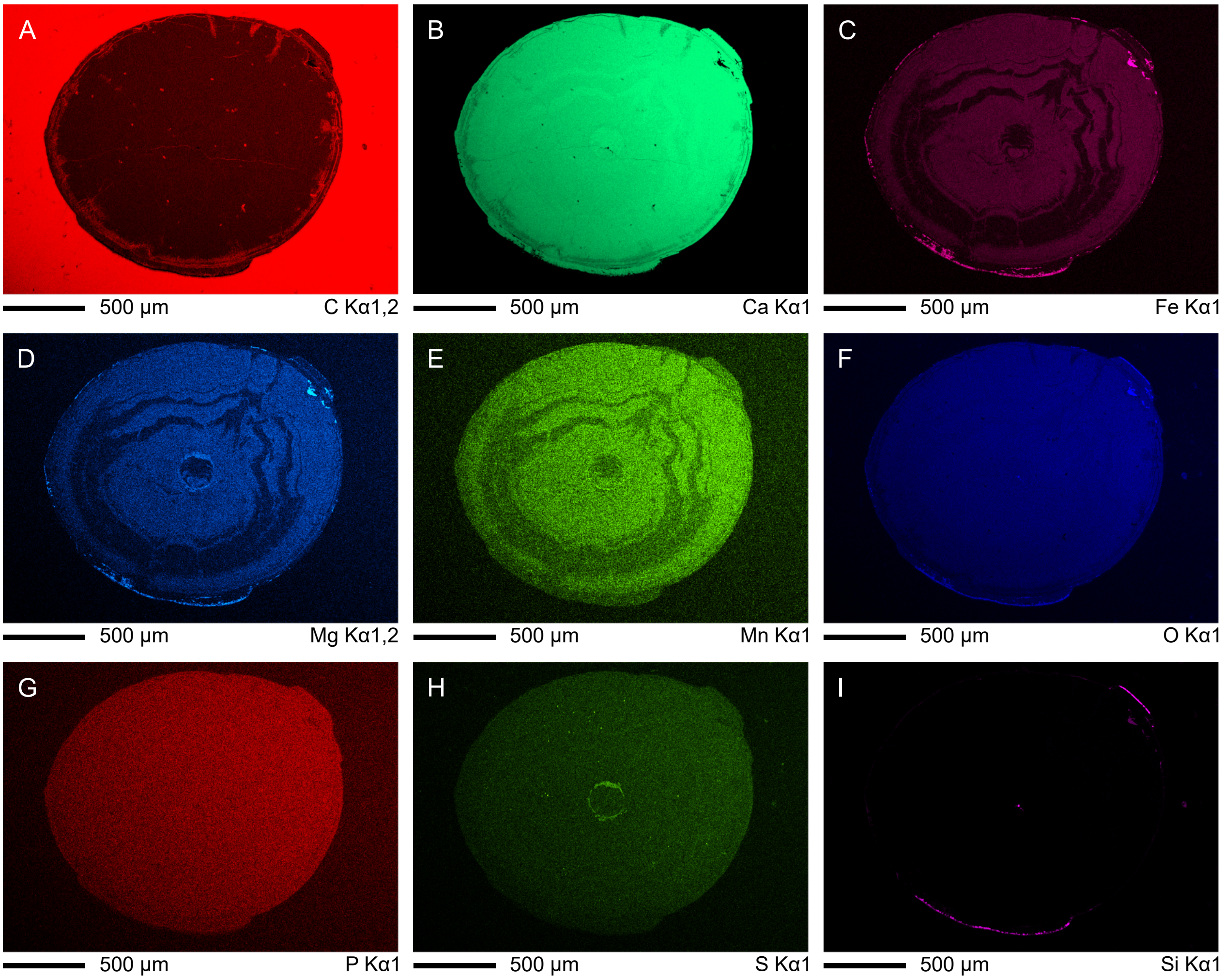


**Fig. S20.** EDS element mappings of *Trematoceras elegans*, PZO 16540-01, cross-section. For a point measurement of this specimen, see Figure 5F, H in the main article. A, carbon. B, calcium. C, iron. D, magnesium. E, manganese. F, oxygen. G, phosphorous. H, sulphur. I, silicon.

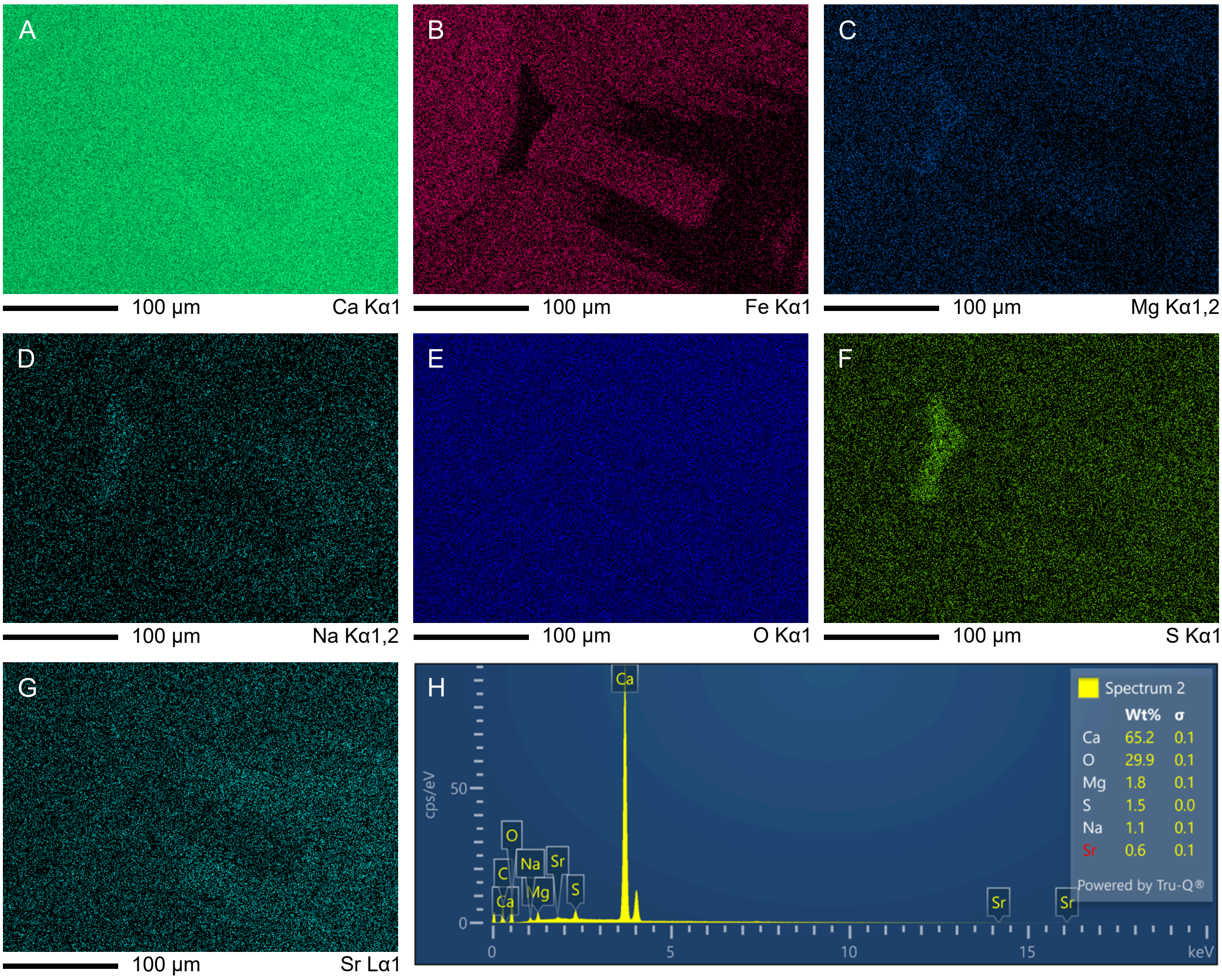


**Fig. S21.** EDS element mappings of *Trematoceras elegans*, PZO 16538-01, cross-section, at biconcave structure ventral to siphuncle (see Fig. 6J-L in the main article). Note that carbon was excluded from the measurements. A, calcium. B, iron. C, magnesium. D, sodium. E, oxygen. F, sulphur. G, strontium. H, point measurement in dorsal (upper left) area, compare Fig. 6L (sc) in the main article.

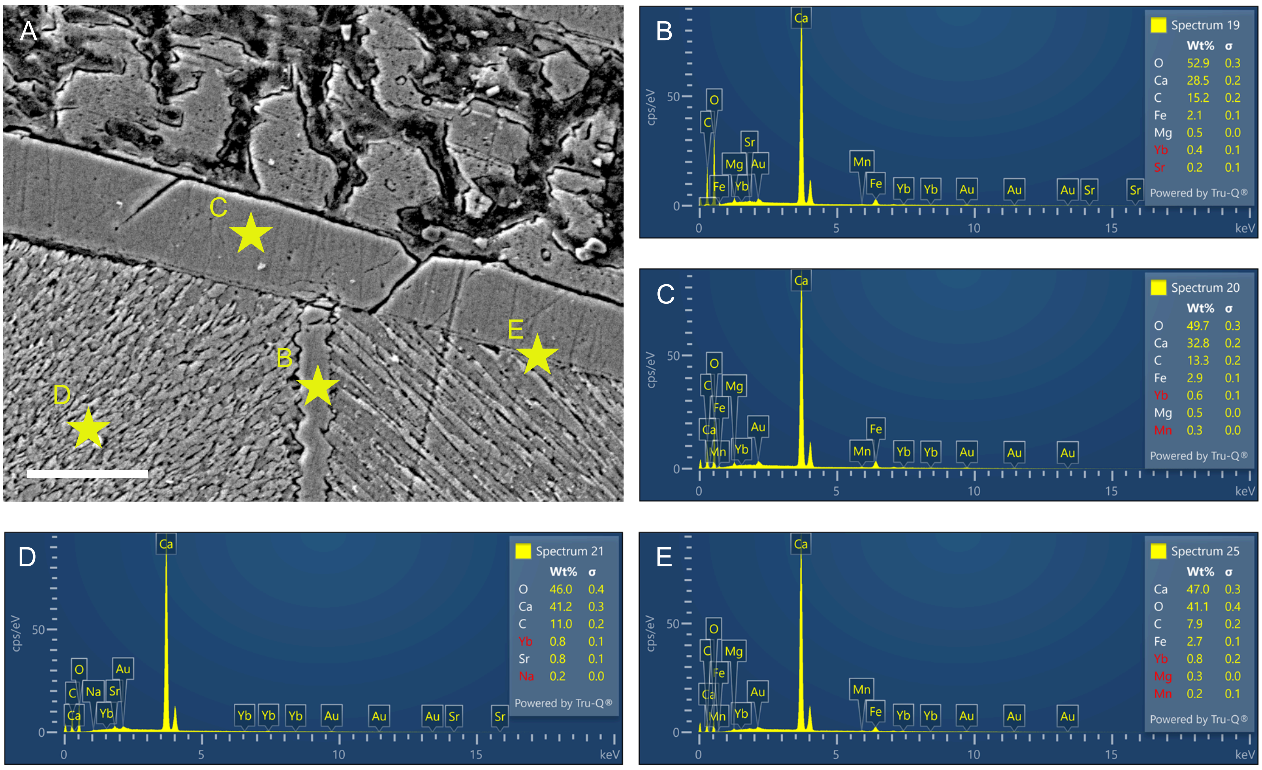


**Fig. S22.** EDS point measurements of *Trematoceras elegans*, PZO 16538-02, longitudinal section (see Fig. 7H). A, Overview of measured points, stars indicating position of point measurements. Scale bar represents 10 µm. B, Point spectrum of sheet-like structure between cameral deposits. C, Point spectrum of diagenetic calcite between broken septum and cameral deposits. D, Point spectrum of aragonitic cameral deposits. E, Point spectrum of sheet-like structure on septal surface of cameral deposits (= pellicle?).

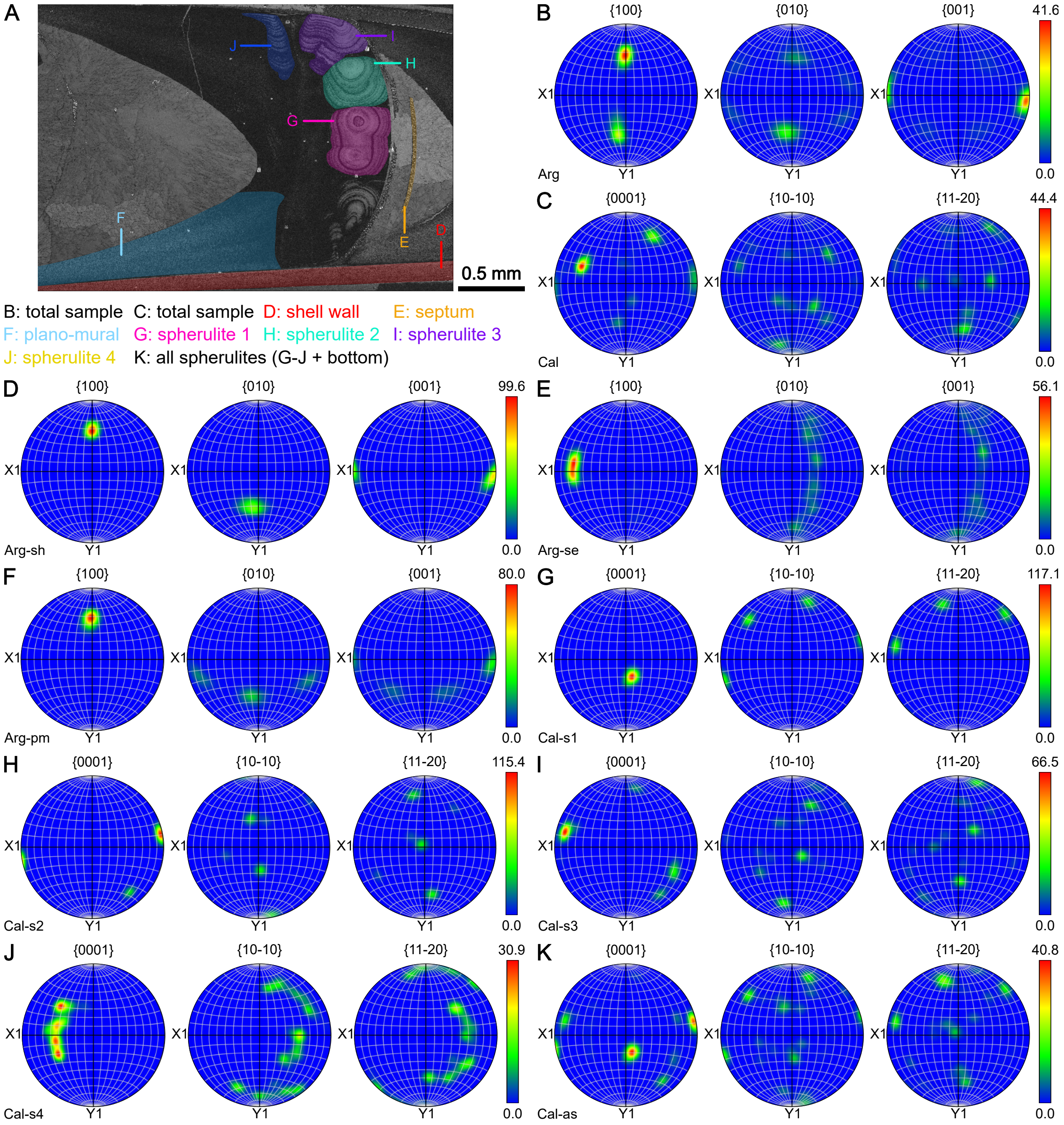


**Fig. S23.** EBSD pole figures of *Trematoceras elegans*, PZO 16538-02, longitudinal section. A, Band contrast image with subsets highlighted. B-K, pole figures, showing crystal orientations. For aragonite (B, D-F), a {100}, b {010} and c{001} axes are shown from left to right. For calcite (C, G-K), c {0001}, $a_1$ {10-10} and $a_2$ {11-20} axes are shown from left to right. B, total sample. C, total sample. D, shell wall. E, septum. F, plano-mural deposits. G-J, individual spherulites. K, all spherulites, including spherulite at bottom of image.

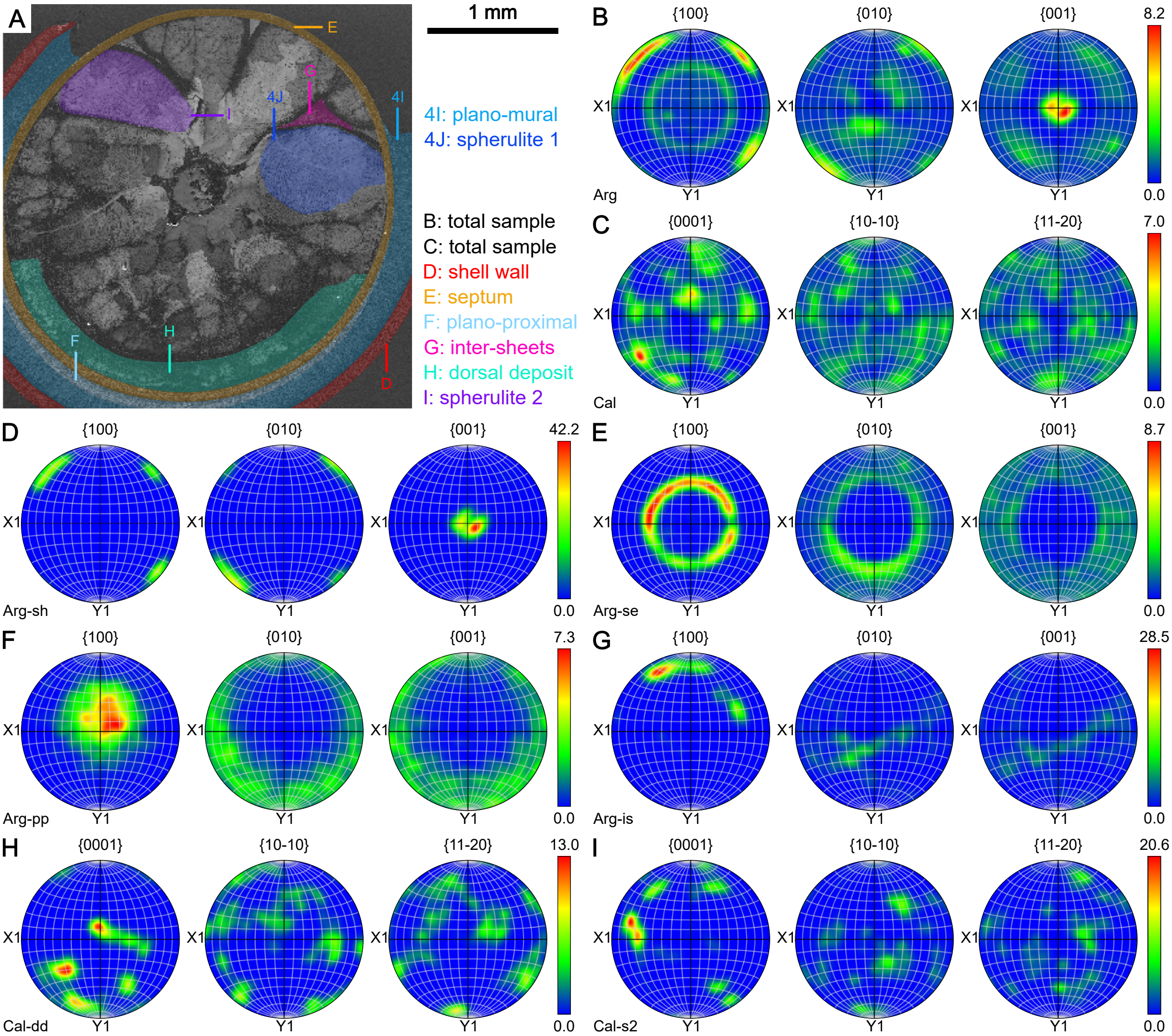


**Fig. S24.** EBSD pole figures of *Trematoceras elegans*, PZO 16535-01, cross section. A, Band contrast image with subsets highlighted. B-I, pole figures, showing crystal orientations. For aragonite (B, D-G), a {100}, b {010} and c{001} axes are shown from left to right. For calcite (C, H-I), c {0001}, $a_1$ {10-10} and $a_2$ {11-20} axes are shown from left to right. B, total sample. C, total sample. D, shell wall. E, septum. F, plano-proximal deposits. G inter-sheet-space. H, dorsal cameral deposits. H, individual spherulite.
